# Alkaloid avoidance in poison frog tadpoles

**DOI:** 10.1101/2022.01.12.476122

**Authors:** Eugenia Sanchez, Travis Ramirez, Lauren A. O’Connell

## Abstract

Animals show a spectrum of avoidance-tolerance to foods containing toxic secondary metabolites. However, this spectrum has not been evaluated in animals that may actively seek out these compounds as a chemical defense. Poison frogs sequester toxic and unpalatable alkaloids from their diet, and in some species, tadpoles are exposed to these toxins before the development of their skin granular glands, which are used for toxin compartmentalization. Here, we examined the effects of the alkaloid decahydroquinoline (DHQ) in tadpoles of the Mimetic poison frog, *Ranitomeya imitator*, using alkaloid supplemented food. We found that although their survival is lowered by the alkaloid, their development is only mildly affected, with no evident effects on their growth. Furthermore, locomotor activity and feeding behavior was altered in the first week of DHQ treatment, probably in part through nicotinic blockade. However, after two weeks, tadpoles learned to avoid the alkaloid by visiting the food area only when necessary to eat. Our results suggest that poison frogs navigate the avoidance-tolerance spectrum during the development of their sequestration machinery.

**Summary:** Animals avoid consuming toxic foods or have anti-toxin machinery to avoid food poisoning. Adult poison frogs eat toxic insects and store the toxins in their skin glands. Some poison frog species even feed their tadpoles with toxins to protect them from predation at the risk of poisoning them. In this study, we observed that toxic food did not affect the development of tadpoles because they quickly learned to eat just enough to grow without getting poisoned. Our results indicate that poison frogs use diverse ways to avoid food poisoning during development.

## Introduction

Animals make choices when approaching food to obtain nutrition and when avoiding food that potentially contain harmful compounds. This approach-avoidance trade-off has been used for decades in studying motivational behaviors linked to eating behavior and distinguishing appealing and aversive foods (Piqueras-Fiszman et al. 2014). Both taste and smell are important sensory inputs that may signal the quality or content of food. For instance, foods containing potentially harmful secondary metabolites might be avoided as they taste bitter (Muñoz et al. 2020). However, some animals rely on these compounds for acquired chemical defenses, such as butterflies that sequester cardenolides and poison frogs that sequester alkaloids (Ruxton et al. 2018). Although approach-avoidance behavior towards secondary metabolites has been investigated in animals that do not rely ecologically on these compounds, how secondary metabolites impact the feeding behavior of animals that actively seek out these compounds has not been investigated.

In the aposematic poison frogs of the family Dendrobatidae, alkaloid-based chemical defenses are obtained from a specialized diet on ants and mites (Daly et al. 1994; Darst et al. 2005; Saporito et al. 2009). Although the physiology and molecular basis of alkaloid processing is not fully understood, poison frogs use a variety of mechanisms to avoid intoxication, including the accumulation of mutations in target ion-channels or receptors (Tarvin et al. 2016, 2017, Yuan and Wang 2018, Márquez et al. 2019), and the compartmentalization of alkaloids mainly in skin granular glands (Saporito et al. 2012). Granular glands start developing after Gosner stage 33 (Stynoski et al. 2014a, Stynoski and O’Connell 2017) and are likely fully operational by the time froglets emerge from the water. However, in some species of poison frogs, tadpoles are exposed to alkaloids before this stage through extended parental care in which mothers feed them with unfertilized eggs that might contain alkaloids (Weygoldt 1987; Stynoski et al. 2014a, Fischer et al. 2019). In other vertebrates, the early consumption of certain alkaloids slows developmental rates, affects adult sizes, and alters behaviors in adulthood due to long-lasting modifications of neural circuits (Levitt et al. 1997; Parker and Connaughton 2007; Dwyer et al. 2008). While alkaloid provisioning gives an antipredator advantage to older tadpoles that can accumulate alkaloids (Stynoski et al. 2014b), it is unknown how early tadpoles lack poison glands for alkaloid compartmentalization may respond to alkaloid exposure.

Here, we examined how alkaloid provisioning affects the development, growth and behavior of poison frog tadpoles. We tested the hypothesis that alkaloid exposure during development would reduce growth rates and lead to deleterious health outcomes due to alkaloid toxicity. We raised freshly hatched tadpoles in either a control or an alkaloid dietary treatment. We used *Ranitomeya imitator* because this species presents maternal egg provisioning (Brown et al. 2008) and, unlike *Oophaga* species, their tadpoles are not obligate egg-feeders, enabling easier testing in controlled laboratory conditions. The alkaloid diet was supplemented with decahydroquinoline (DHQ), a commercially available alkaloid of the same class of major alkaloids found in this species in nature (Stuckert et al. 2014).

## Methods

### Feeding experiments

Freshly hatched, full-sibling tadpoles of *Ranitomeya imitator* were randomly allocated to control or alkaloid diet treatments in two separate experiments, one with 1.0% DHQ (N = 37 = 24 control + 13 DHQ) and another with 0.5% DHQ (N = 54 = 27 control + 27 DHQ). Control diet consisted in Brine shrimp flakes (Josh’s Frogs, Owosso, MI, USA), and alkaloid diet consisted of the same flakes with either 1.0% or 0.5% m/m DHQ (Sigma-Aldrich, St Louis, MO, USA). Feeding occurred twice per week. In the 1.0% DHQ experiment, we offered approx. 5 mg of flakes both times, while in the 0.5% DHQ experiment, we offered one pellet (made out of the flakes) of approx. 10 mg during the behavioral trial, and approximately 1 mg flakes two days later (Figure SM1). In both experiments, both treatments received control food in the first feeding on week 0, and the water was changed the day before the behavioral trials. The 1.0%-DHQ experiment lasted 10 weeks, and the 0.5%-DHQ experiment lasted 3 weeks. All animals were kept in individual transparent polypropylene cups with 60 ml of water at 28 ± 1 °C, and with shade and open areas (Figure SM1). These cups were placed in a box to provide a green background (Figure SM1), and kept in a room with a 12h hour light/dark cycle.

### Mortality, development and growth

Once a week, we recorded the mortality and developmental stage of each individual (Gosner 1960). Tadpoles were weighed and photographed in triplicate, from which we measured the snout-tail length (STL), snout-vent length (SVL), and head width (eye-width) with ImageJ (Schneider et al. 2012).

### Behavioral trials

Once per week, individuals were subjected to an open field trial and a feeding behavior trial. Individuals were placed in a 10×10 cm behavioral arena filled with water at a temperature of 31 ± 1 °C (Figure SM3). In the 1.0%-DHQ experiment, individuals were acclimatized to the arena for 5 min, then movement behavior in an “open field” was scored for 10 min, after which tadpoles were presented with food and behavior was scored for feeding 15 min. In the 0.5%-DHQ experiment, tadpoles were acclimatized to the arena for 15 min and then movement behavior in an “open field” was scored for 30 min, after which tadpoles were presented with food and behavior was scored for feeding 30 min. Both the open field and feeding behavioral trials were video recorded using a GoPro model 6 (GoPro, Inc., San Mateo, CA, USA) on top of the arenas, at 960p/120fps. Trials were performed between 9 am and 5 pm in the 1.0%-DHQ, and 1 pm and 5 pm in the 0.5%-DHQ experiment.

Individual’s locomotor activity during the open field trials was analyzed using ToxTrac (Rodriguez et al. 2018), and we calculated the average speed of movement, mobility rate (percent of time moving), exploration rate (number of areas visited when dividing the arena in a 10×10 grid), and the frequency individuals visited the arena’s center considering a 1.5 cm margin, using a customized python script (toxtrac_summary.py). Behavioral events during the feeding trials were counted and measured with BORIS v. 7.9.8 (Friard and Gamba 2016). We scored the following behaviors: smelling, when the tadpoles moved their mouths away from the food; testing, when the tadpole’s mouth was very close to the food or it was unclear whether they were eating; eating, when the tadpole’s mouth was touching the food; near, when the tadpole’s mouth was less than 5 mm from the food. From the Boris log files, we further estimated the latency to each behavior from the moment the food was placed, using a customized python script (boris_latency.py).

### Data analysis

Tadpole survival was compared between treatments using a Chi-square test with the R package *survival* (Therneau 2020). Longitudinal data (development, growth, movement and feeding behavior) were analyzed using Linear Mixed-Effects models with the R package *nlme* (Pinheiro et al. 2020), considering week, treatment, and parents as fixed factors, and individual as a random factor. Latencies were compared between treatments per week, using a Chi-square test with the R package *survival* (Therneau 2020).

## Results

### Survival, development and growth

The probability of survival did not differ between diet treatments in the 0.5% DHQ experiment (X^2^ (1, N = 54) = 0.6, p = 0.4; Figure 1), but more tadpoles died in the 1.0% DHQ treatment after 10 weeks of experiment (X^2^ (1, N= 37) = 4, p = 0.05; Figure 1). However, DHQ treatments did not affect the developmental rate, weight, or size of the tadpoles (LME treatment: p > 0.05). Figures and statistics are provided in the supplementary R Markdown.

**Figure 1.**
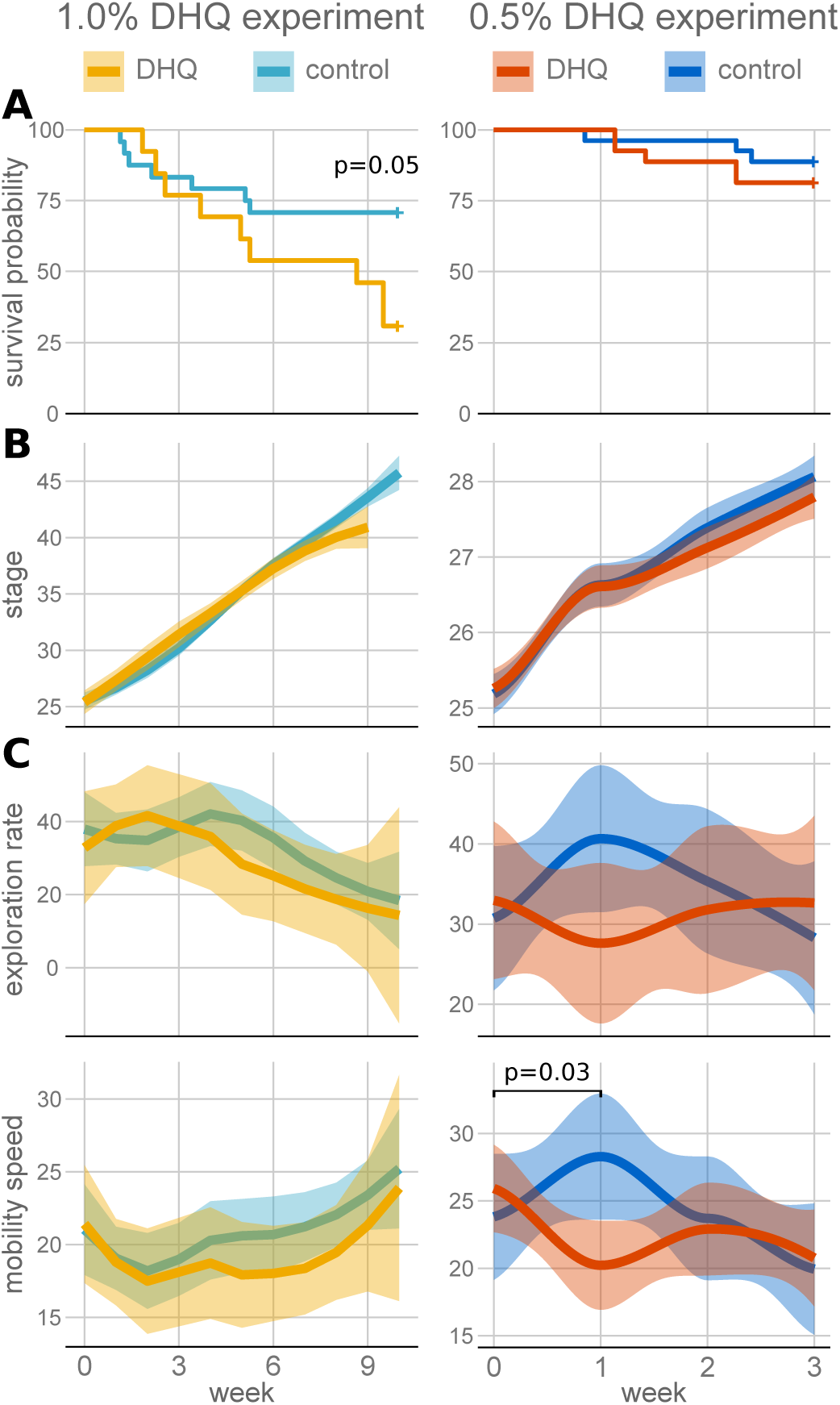
Survival (A), development (B) and locomotor activity (C) in the open field trials for both 1.0% DHQ (left column) and 0.5% DHQ (right column) experiments. The survival probability was significantly lower after 10 weeks of 1.0% DHQ treatment, but not after three weeks of 0.5% DHQ treatment. Although the mobility speed (mm/s), and exploration rate (% arena visited) were not significantly different across the whole experiments, the mobility speed was different between treatments in the first week of 0.5% DHQ treatment.

### Locomotor activity

In the first week of treatment, 0.5% DHQ and 1.0% DHQ tadpoles tended to move slower and explore a smaller area of the arena than the controls (Figure 1; LME treatment: p>0.01). Suggesting that initial exposure to alkaloids has a mild influence on locomotor activity.

### Feeding behavior

The behavior of 0.5% DHQ-fed tadpoles changed throughout the three weeks of experiment compared to controls (Figure 2). DHQ tadpoles had a higher latency to test the food (X^2^ (1, N = 44) = 3.96, p = 0.05) and eat it in the first week (X^2^ (1, N = 44) = 4.15, p = 0.04), and a higher latency to smell the food in the second week (X^2^ (1, N = 46) = 4.71, p = 0.03). These behavioral changes lowered the number of times DHQ tadpole tested and ate the food in the second and third weeks (testing LME treatment*week: beta = -0.38, t = -2.48, p = 0.015; eating LME treatment*week: beta = -0.43, t = -2.54, p = 0.013), and tended to influence the number of times and percent of time they spent close to the food (LME p > 0.05). Overall, tadpoles developed avoidance to alkaloid-containing food.

**Figure 2.**
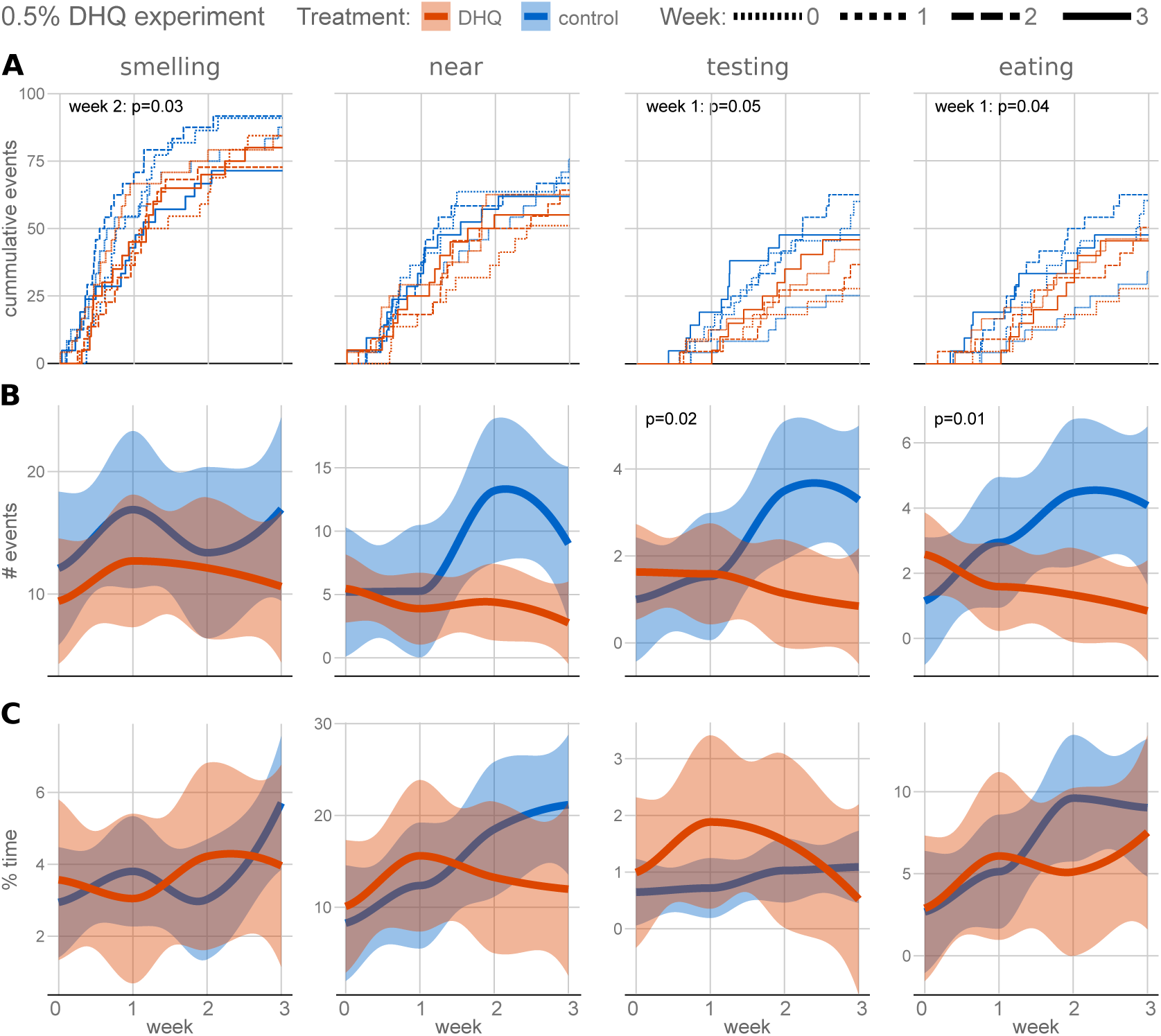
Changes in feeding behavior in the 0.5% DHQ experiment. We scored the behaviors smelling, when the tadpoles moved their mouths away from the food, near, when the tadpole was close to the food, testing, when they moved it close to the food, and eating, when the tadpole’s mouth was touching the food. The plots show **(A)** the latency to each behavior, **(B)** the number of events of each behavior, and **(C)** the percentage of time spent in each behavior, in the 30 min feeding trials conducted once a week.

## Discussion

Here, we examined how alkaloid provisioning may alter the growth and behavior of young poison frog tadpoles. We specifically tested the hypothesis that alkaloid exposure during development would reduce growth, increase activity level and eating behavior, and lead to deleterious health outcomes due to alkaloid toxicity from lack of granular glands for compartmentalization. Contrary to our initial hypotheses, we found alkaloid exposure through the diet did not influence growth, influenced the survival mildly, and instead induced learned avoidance of alkaloid-containing food. We discuss these findings in the context of poison frog alkaloid physiology, as well as within the larger literature of learned toxin avoidance in aquatic animals.

Alkaloid processing is considered costly for poison frogs as their consumption upregulates mitochondrial pathways (Sanchez et al. 2019, Caty et al. 2019) and is associated with the evolution of higher metabolic rates (Santos and Cannatella 2011). Based on these metabolic costs, we hypothesized that alkaloid provisioning would slow tadpole growth rates. However, we instead found that DHQ-exposure in the diet did not significantly alter the developmental rate of *R. imitator* tadpoles compared to controls. It is possible that tadpoles do not display the same physiological changes associated with alkaloid exposure that has been observed in adult poison frogs. Although we observed changes in the number of breaths taken by DHQ tadpoles, these are explained by the changes in activity level, and not by the time in the treatment (see R Markdown). Hence, future work focusing on gene expression or metabolic rate measurements would shed more light on how alkaloids may influence metabolism differently across amphibian life stages.

DHQs are non-competitive blockers of nicotinic acetylcholine receptors (Daly et al. 1991), and *R. imitator* does not present mutations in these receptors that would alter the receptor dynamics that has been reported in other poison frogs (Tarvin et al. 2017). In mammals, nicotinic blockade limits the behavioral effects of the peptide hormone ghrelin, which include locomotor activity and feeding behavior (Jerlhag et al. 2006; Dickson et al. 2010; Perelló and Zigman 2012). In agreement, we found that tadpoles reduced their activity levels, and showed a higher latency to start eating in the first week of DHQ treatment. Although tadpoles spent similar amounts of time eating, we consider latency to eat as a form of food avoidance because food avoidance *per se* would not persist as there were no food alternatives offered (e.g., Kimball et al. 2002). In this sense, our experiments do not measure preference but mimic the case of obligate egg-feeders with alkaloid provisioning. Nicotinic blockade decreases the food’s rewarding properties (Dickson et al. 2010), and DHQs are considered to have a noxious, bitter taste (Santos et al. 2016). Together, these effects seemed to have resulted in tadpoles learning to avoid the DHQ food after two weeks, as they visited the food area less times than the controls. Noticeably, tadpoles’ latency to smell was also increased in the second week of DHQ treatment. Such spatial avoidance, which has been observed previously in other aquatic organisms exposed to contaminants (Araujo et al. 2016), might serve as a reasonable compromise between necessary feeding and alkaloid exposure.

Our initial hypotheses about the influence of alkaloids on early tadpole physiology and behavior were based on their inability to store alkaloids in granular glands that have not yet developed and the lack of mutations in the target receptor that regulates locomotor activity and feeding behavior in other vertebrates. More recently, other toxin-resistance mechanisms have come to light, including the proposal of a carrier protein or “toxin sponge” that may protect the nervous system from alkaloids (Abderemane-Ali et al. 2021). Although these anti-toxin factors are still a hypothesis lacking empirical data, they represent a potential alternative explanation of how tadpole behavior and growth are relatively buffered against alkaloid toxicity.

Animals show a spectrum of avoidance-tolerance to foods containing secondary metabolites (Iason and Villalba 2006). This spectrum can also be considered as a behavior-physiology spectrum, as avoidance is achieved through limiting food amount or frequency of intake, while tolerance requires adaptive resistance mechanisms. On an evolutionary scale, we recognize poison frogs may occupy different parts of this continuum, with species specialized in alkaloid containing diets showing target mutations and the ability to sequester and compartmentalize alkaloids in one of the extremes (Toft 1995, Daly et al. 2003, Darst et al. 2005, Tarvin et al. 2016, Tarvin et al. 2017, Yuan and Wang 2018, Márquez et al. 2019, O’Connell et al. 2021). While target mutations might be expressed throughout the life time of individuals, compartmentalization depends on poison gland development, which leads poison frogs to navigate through the avoidance-tolerance spectrum during development. Future work investigating how alkaloids impact poison frog’s physiology and behavior across different life stages and across chemically defended and non-defended dendrobatids will shed light on the principles guiding avoidance-tolerance and its underlying molecular mechanisms.

## Funding

ES was funded through fellowships from the Deutsche Forschungsgemeinschaft (SA 3693/1-1 and SA 3693/2-1). This research was supported by the National Science Foundation (IOS-1845651), the National Institutes of Health (DP2HD102042), and a Hellman Fellow Award to LAO. LAO is a New York Stem Cell Foundation – Robertson Investigator.

## Ethics

All animal protocols were approved by the Stanford University Institutional Animal Care and Use Committee (protocol #33097).

## Competing interests

We have no competing interests.

## Data accessibility

Behavioral videos and pictures can be accessed upon request. Customized python scripts are available at https://github.com/eu-sanisa/DHQ_feeding_python. Datasets and R Markdown for data analysis are published as Supplementary Materials.

## Authors’ contributions

ES and LAO conceived the study and designed the experiments; ES performed feeding experiments and recorded data; ES and TR measured tadpoles from pictures; ES evaluated videos; ES analyzed data; ES and LAO wrote the manuscript with input from TR.

## Acknowledgments

We thank the Laboratory of Organismal Biology at Stanford University for their emotional and scientific support, particularly Jessica Nowicki and Billie Goolsby for their support implementing molecular protocols. We thank Marian Hebenbrock for his advice preparing the food treatments. ES thanks Heike Pröhl for her trust and support when transitioning back to Germany.

## SUPPLEMENTARY MATERIALS

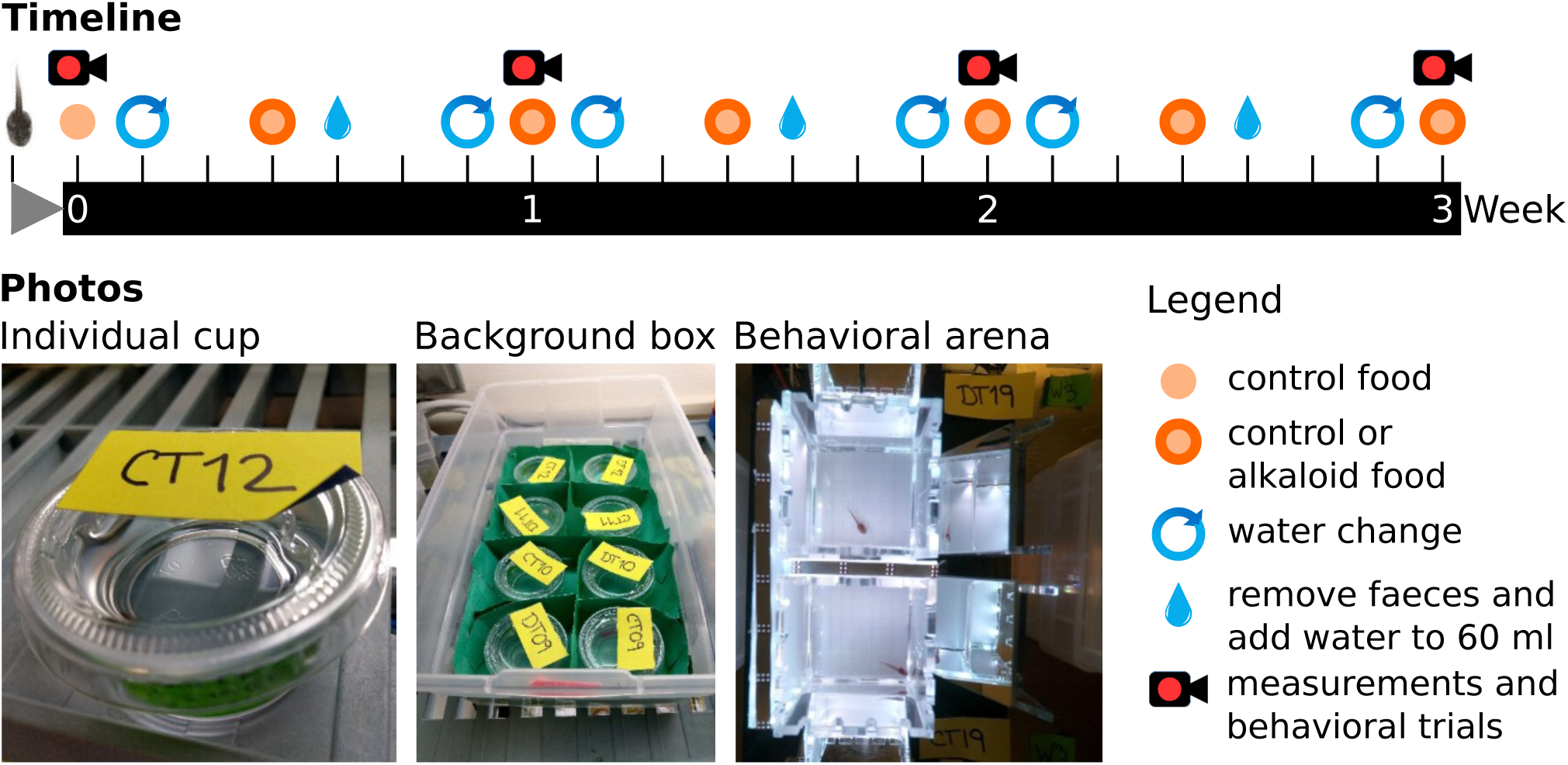

**SM1. Experimental protocol**. The timeline shows the protocol for the three weeks of 0.5% DHQ experiment. The 1.0% DHQ experiment continued similarly for other seven weeks.

**SM2. SM tables**. Excel document containing spread-sheets with data on tadpole survival, measurements, movement and feeding behavior.

**SM3. R Markdown**. HTML document with the R code used for data analysis, and the resulting graphics and statistics.

**SM4. Alignment**. Sequences of nicotinic acetylcholine receptor beta-2 (chrnb2) for R. imitator and R. ventrimaculata showing lack of desensitizing mutations.

## Notes

### Competing Interest Statement

The authors have declared no competing interest.

## References

Abderemane-Ali, F., Rossen, N.D., Kobiela, M.E., Craig, R.A., Garrison, C.E., Chen, Z., Colleran, C.M., O’Connell, L.A., Du Bois, J., Dumbacher, J.P. and Minor Jr, D.L., 2021. Evidence that toxin resistance in poison birds and frogs is not rooted in sodium channel mutations and may rely on “toxin sponge” proteins. Journal of General Physiology, 153(9), p.e202112872.

Araújo, C.V., Moreira-Santos, M. and Ribeiro, R., 2016. Active and passive spatial avoidance by aquatic organisms from environmental stressors: a complementary perspective and a critical review. Environment International, 92, pp.405–415.

Brown, J.L., Twomey, E., Morales, V. and Summers, K., 2008. Phytotelm size in relation to parental care and mating strategies in two species of Peruvian poison frogs. Behaviour, pp.1139–1165.

Caty, S.N., Alvarez-Buylla, A., Byrd, G.D., Vidoudez, C., Roland, A.B., Tapia, E.E., Budnik, B., Trauger, S.A., Coloma, L.A. and O’Connell, L.A., 2019. Molecular physiology of chemical defenses in a poison frog. Journal of Experimental Biology, 222(12), p.jeb204149.

Daly, J.W., Nishizawa, Y., Padgett, W.L., Tokuyama, T., McCloskey, P.J., Waykole, L., Schultz, A.G. and Aronstam, R.S., 1991. Decahydroquinoline alkaloids: Noncompetitive blockers for nicotinic acetylcholine receptor-channels in pheochromocytoma cells and Torpedo electroplax. Neurochemical Research, 16(11), pp.1207–1212.

Daly, J.W., Garraffo, H.M., Spande, T.F., Jaramillo, C. and Rand, A.S., 1994. Dietary source for skin alkaloids of poison frogs (Dendrobatidae)?. Journal of Chemical Ecology, 20(4), pp.943–955.

Daly, J.W., Garraffo, H.M., Spande, T.F., Clark, V.C., Ma, J., Ziffer, H. and Cover, J.F., 2003. Evidence for an enantioselective pumiliotoxin 7-hydroxylase in dendrobatid poison frogs of the genus Dendrobates. Proceedings of the National Academy of Sciences, 100(19), pp.11092–11097.

Darst, C.R., Menéndez-Guerrero, P.A., Coloma, L.A. and Cannatella, D.C., 2005. Evolution of dietary specialization and chemical defense in poison frogs (Dendrobatidae): a comparative analysis. The American Naturalist, 165(1), pp.56–69.

Dickson, S.L., Hrabovszky, E., Hansson, C., Jerlhag, E., Alvarez-Crespo, M., Skibicka, K.P., Molnar, C.S., Liposits, Z., Engel, J.A. and Egecioglu, E., 2010. Blockade of central nicotine acetylcholine receptor signaling attenuate ghrelin-induced food intake in rodents. Neuroscience, 171(4), pp.1180–1186.

Dwyer, J.B., Broide, R.S. and Leslie, F.M., 2008. Nicotine and brain development. Birth Defects Research Part C: Embryo Today: Reviews, 84(1), pp.30–44.

Fischer, E.K., Roland, A.B., Moskowitz, N.A., Tapia, E.E., Summers, K., Coloma, L.A. and O’Connell, L.A., 2019. The neural basis of tadpole transport in poison frogs. Proceedings of the Royal Society B, 286(1907), p.20191084.

Friard, O. and Gamba, M., 2016. BORIS: a free, versatile open-source event-logging software for video/audio coding and live observations. Methods in Ecology and Evolution, 7(11), pp.1325–1330.

Gosner, K.L., 1960. A simplified table for staging anuran embryos and larvae with notes on identification. Herpetologica, 16(3), pp.183–190.

Iason, G.R. and Villalba, J.J., 2006. Behavioral strategies of mammal herbivores against plant secondary metabolites: the avoidance–tolerance continuum. Journal of chemical ecology, 32(6), pp.11151132.

Jerlhag, E., Egecioglu, E., Dickson, S.L., Andersson, M., Svensson, L. and Engel, J.A., 2006. PRE-CLINICAL STUDY: Ghrelin stimulates locomotor activity and accumbal dopamine-overflow via central cholinergic systems in mice: implications for its involvement in brain reward. Addiction Biology, 11(1), pp.45–54.

Kimball, B.A., Provenza, F.D. and Burritt, E.A., 2002. Importance of alternative foods on the persistence of flavor aversions: implications for applied flavor avoidance learning. Applied Animal Behaviour Science, 76(3), pp.249–258.

Levitt, P., Harvey, J.A., Friedman, E., Simansky, K. and Murphy, E.H., 1997. New evidence for neurotransmitter influences on brain development. Trends in neurosciences, 20(6), pp.269–274.

Márquez, R., Ramírez-Castañeda, V. and Amézquita, A., 2019. Does batrachotoxin autoresistance coevolve with toxicity in Phyllobates poison-dart frogs?. Evolution, 73(2), pp.390–400.

Muñoz, I.J., Schilman, P.E. and Barrozo, R.B., 2020. Impact of alkaloids in food consumption, metabolism and survival in a blood-sucking insect. Scientific Reports, 10(1), pp.1–10.

O’Connell, L.A., LS50: Integrated Science Laboratory Course, O’Connell, J.D., Paulo, J.A., Trauger, S.A., Gygi, S.P. and Murray, A.W., 2021. Rapid toxin sequestration modifies poison frog physiology. Journal of Experimental Biology, 224(3), p.jeb230342.

Parker, B. and Connaughton, V.P., 2007. Effects of nicotine on growth and development in larval zebrafish. Zebrafish, 4(1), pp.59–68.

Perelló, M. and Zigman, J.M., 2012. The role of ghrelin in reward-based eating. Biological psychiatry, 72(5), pp.347–353.

Pinheiro J, Bates D, DebRoy S, Sarkar D, R Core Team (2020). nlme: Linear and Nonlinear Mixed Effects Models. R package version 3.1-150, https://CRAN.R-project.org/package=nlme.

Piqueras-Fiszman, B., Kraus, A.A. and Spence, C., 2014. “Yummy” versus “Yucky”! Explicit and implicit approach–avoidance motivations towards appealing and disgusting foods. Appetite, 78, pp.193–202.

Rodriguez, A., Zhang, H., Klaminder, J., Brodin, T., Andersson, P.L. and Andersson, M., 2018. ToxTrac: a fast and robust software for tracking organisms. Methods in Ecology and Evolution, 9(3), pp.460–464.

Ruxton, G.D., Allen, W.L., Sherratt, T.N. and Speed, M.P., 2018. Avoiding attack: the evolutionary ecology of crypsis, aposematism, and mimicry. Second edition. Oxford University Press. Pp. 278.

Sanchez, E., Rodríguez, A., Grau, J.H., Lötters, S., Künzel, S., Saporito, R.A., Ringler, E., Schulz, S., Wollenberg Valero, K.C. and Vences, M., 2019. Transcriptomic signatures of experimental alkaloid consumption in a poison frog. Genes, 10(10), p.733.

Santos, J.C. and Cannatella, D.C., 2011. Phenotypic integration emerges from aposematism and scale in poison frogs. Proceedings of the National Academy of Sciences, 108(15), pp.6175–6180.

Santos, J.C., Tarvin, R.D. and O’Connell, L.A., 2016. A review of chemical defense in poison frogs (Dendrobatidae): ecology, pharmacokinetics, and autoresistance. Chemical signals in vertebrates 13, pp.305–337.

Saporito, R.A., Spande, T.F. and Garraffo, H.M., 2009. Arthropod alkaloids in poison frogs: a review of the’dietary hypothesis’. Heterocycles, 79(1), pp.277–297.

Saporito, R.A., Donnelly, M.A., Spande, T.F. and Garraffo, H.M., 2012. A review of chemical ecology in poison frogs. Chemoecology, 22(3), pp.159–168.

Schneider, C.A., Rasband, W.S. and Eliceiri, K.W., 2012. NIH Image to ImageJ: 25 years of image analysis. Nature Methods, 9(7), pp.671–675.

Stuckert, A.M., Saporito, R.A., Venegas, P.J. and Summers, K., 2014. Alkaloid defenses of co-mimics in a putative Müllerian mimetic radiation. BMC Evolutionary Biology, 14(1), pp.1–8.

Stynoski, J.L. and O’Connell, L.A., 2017. Developmental morphology of granular skin glands in premetamorphic egg-eating poison frogs. Zoomorphology, 136(2), pp.219–224.

Stynoski, J.L., Torres-Mendoza, Y., Sasa-Marin, M. and Saporito, R.A., 2014a. Evidence of maternal provisioning of alkaloid-based chemical defenses in the strawberry poison frog Oophaga pumilio. Ecology, 95(3), pp.587–593.

Stynoski, J.L., Shelton, G. and Stynoski, P., 2014b. Maternally derived chemical defences are an effective deterrent against some predators of poison frog tadpoles (Oophaga pumilio). Biology Letters, 10(5), p.20140187

Tarvin, R.D., Santos, J.C., O’Connell, L.A., Zakon, H.H. and Cannatella, D.C., 2016. Convergent substitutions in a sodium channel suggest multiple origins of toxin resistance in poison frogs. Molecular Biology and Evolution, 33(4), pp.1068–1080.

Tarvin, R.D., Borghese, C.M., Sachs, W., Santos, J.C., Lu, Y., O’Connell, L.A., Cannatella, D.C., Harris, R.A. and Zakon, H.H., 2017. Interacting amino acid replacements allow poison frogs to evolve epibatidine resistance. Science, 357(6357), pp.1261–1266.

Therneau T (2020). A Package for Survival Analysis in R. R package version 3.2-7, https://CRAN.R-project.org/package=survival.

Toft, C.A., 1995. Evolution of diet specialization in poison-dart frogs (Dendrobatidae). Herpetologica, pp.202–216.

Weygoldt, P., 1987. Evolution of parental care in dart poison frogs (Amphibia: Anura: Dendrobatidae). Journal of Zoological Systematics and Evolutionary Research, 25(1), pp.51–67.

Yuan, M.L. and Wang, I.J., 2018. Sodium ion channel alkaloid resistance does not vary with toxicity in aposematic Dendrobates poison frogs: An examination of correlated trait evolution. PloS One, 13(3), p.e0194265.

