## Supplementary material for "Alkaloid avoidance in poison frog tadpoles": SM3

#### 2021

Packages used in this Markdown:

```
library(tidyr)
library(FSA)
library(ggplot2)
library(ggthemes)
library(survival)
library(survminer)
library(lme4)
library(lmerTest)
library(rstatix)
library(nlme)
library(kableExtra)
```

The data used in this markdown are provided as tables (spread\_sheets) in the SM1\_tables.xlsx document.

### Survival

The survival table looks like this:

```
survival = read.table('D:/STANFORD/DHQ_FEEDING/code/SM_tables/survival.csv', header=TRUE, sep=',')
kbl(head(survival), caption = "Head of survival table") %>% kable_paper("hover", full_width = F)
```

Head of survival table

| individual | treatment | experiment | tank | female | male | hatch\_date | end\_date | days | censored | comments |
| --- | --- | --- | --- | --- | --- | --- | --- | --- | --- | --- |
| CT01 | control | p05 | B15 | F40 | M38 | 2019-08-05 | 2019-08-26 | 21 | 1 |  |
| CT02 | control | p05 | C10 | F35 | M68 | 2019-08-05 | 2019-08-26 | 21 | 1 |  |
| CT03 | control | p05 | A14 | F50 | M51 | 2019-08-07 | 2019-08-28 | 21 | 1 |  |
| CT04 | control | p05 | B10 | F52 | M92 | 2019-08-12 | 2019-09-02 | 21 | 1 |  |
| CT05 | control | p05 | B16 | F21 | M14 | 2019-08-12 | 2019-09-02 | 21 | 1 |  |
| CT06 | control | p05 | A15 | F54 | M47 | 2019-08-12 | 2019-09-02 | 21 | 1 |  |

##### Plots and statistics

For analysis and plotting we separate the 0.5% DHQ and 1.0% DHQ experiments.

```
survival05 = subset(survival, experiment == 'p05')
survival05$days = ifelse(survival05$days > 21, survival05$days == 21, survival05$days)
survdiff(Surv(days, censored) ~ treatment, data = survival05)
```

```
## Call:
## survdiff(formula = Surv(days, censored) ~ treatment, data = survival05)
## 
##                    N Observed Expected (O-E)^2/E (O-E)^2/V
## treatment=control 27        3     4.08     0.287     0.599
## treatment=dhq     27        5     3.92     0.299     0.599
## 
##  Chisq= 0.6  on 1 degrees of freedom, p= 0.4
```

```
fit05 <- survfit(Surv(days, censored) ~ treatment, data = survival05)
ggsurvplot(fit05, data=survival05, xlab = "days", ylab = 'survival probability', title = '0.5% DHQ experiment', palette = c("#0064c8", "#dc4600"))
```

```
survival10 = subset(survival, experiment == 'p10')
survival10$days = ifelse(survival10$days > 70, survival10$days == 70, survival10$days)
survdiff(Surv(days, censored) ~ treatment, data = survival10)
```

```
## Call:
## survdiff(formula = Surv(days, censored) ~ treatment, data = survival10)
## 
##                    N Observed Expected (O-E)^2/E (O-E)^2/V
## treatment=control 24        7    10.72      1.29      3.97
## treatment=dhq     13        9     5.28      2.61      3.97
## 
##  Chisq= 4  on 1 degrees of freedom, p= 0.05
```

```
fit10 <- survfit(Surv(days, censored) ~ treatment, data = survival10)
ggsurvplot(fit10, data=survival10, xlab = "days", ylab = "survival probability", title = '1.0% DHQ experiment', palette = c("#3caac8", "#f0aa00"))
```

```
survival = rbind(survival05, survival10)
fit <- survfit(Surv(days, censored) ~ treatment + experiment, data = survival)
ggsurvplot(fit, data=survival, xlab = "days", ylab = "survival probability", palette = c("#0064c8", "#3caac8", "#dc4600", "#f0aa00"), xlim = c(0, 70), ggtheme = theme_gdocs())
```

### Development and growth

The measurements table looks like this:

```
tad_data = read.table('D:/STANFORD/DHQ_FEEDING/code/SM_tables/measurements.csv', header=TRUE, sep=',')
kbl(head(tad_data), caption = "Head of measurements table") %>% kable_paper("hover", full_width = F)
```

Head of measurements table

| id | treatment | individual | week | stage | tank | stl | svl | weight | width | tail | area | experiment |
| --- | --- | --- | --- | --- | --- | --- | --- | --- | --- | --- | --- | --- |
| CT01W0 | control | CT01 | 0 | 25.0 | B15 | 10.6 | 4.6 | 25.7 | 2.4 | 6.1 | 34.5 | p05 |
| CT01W1 | control | CT01 | 1 | NA | B15 | 14.0 | 5.2 | 38.0 | 2.8 | 8.8 | 45.6 | p05 |
| CT01W2 | control | CT01 | 2 | 28.0 | B15 | 14.2 | 4.7 | 52.3 | 2.3 | 9.6 | 33.4 | p05 |
| CT01W3 | control | CT01 | 3 | 28.5 | B15 | 16.8 | 6.1 | 58.3 | 2.4 | 10.8 | 46.1 | p05 |
| CT02W0 | control | CT02 | 0 | 25.0 | C10 | 13.4 | 4.9 | 33.7 | 2.7 | 8.6 | 41.5 | p05 |
| CT02W1 | control | CT02 | 1 | NA | C10 | 16.1 | 5.8 | 43.0 | 2.9 | 10.3 | 53.5 | p05 |

We split this table by experiment.

```
tads05 = subset(tad_data, experiment == 'p05')
tads10 = subset(tad_data, experiment == 'p10')
```

#### Development

##### Plots and statistics

```
# lmer1: random intercepts for different individuals but week and treatment effects are fixed 
lmer1 <- lmer(stage ~ week * treatment + tank + (1 | individual), tads05)
# lmer2: random intercepts and treatment effects for different individuals, but the week effect is still fixed
lmer2 <- lmer(stage ~ week * treatment + tank + (1 + treatment | individual), tads05)
# If P<0.5, the best fit is the one with the lowest logLik or lowest AIC/BIC, else, use lmer1
anova(lmer1, lmer2)
```

```
## Data: tads05
## Models:
## lmer1: stage ~ week * treatment + tank + (1 | individual)
## lmer2: stage ~ week * treatment + tank + (1 + treatment | individual)
##       npar    AIC    BIC  logLik deviance  Chisq Df Pr(>Chisq)
## lmer1   19 387.98 450.36 -174.99   349.98                     
## lmer2   21 391.80 460.75 -174.90   349.80 0.1711  2      0.918
```

```
# lmer3: same as lmer1 but removing tank effects
lmer3 <- lmer(stage ~ week * treatment + (1 | individual), tads05)
anova(lmer1, lmer3)
```

```
## Data: tads05
## Models:
## lmer3: stage ~ week * treatment + (1 | individual)
## lmer1: stage ~ week * treatment + tank + (1 | individual)
##       npar    AIC    BIC  logLik deviance  Chisq Df Pr(>Chisq)    
## lmer3    6 405.96 425.66 -196.98   393.96                         
## lmer1   19 387.98 450.36 -174.99   349.98 43.981 13  3.091e-05 ***
## ---
## Signif. codes:  0 '***' 0.001 '**' 0.01 '*' 0.05 '.' 0.1 ' ' 1
```

```
dev.lmer = lmer1
summary(dev.lmer)
```

```
## Linear mixed model fit by REML. t-tests use Satterthwaite's method [
## lmerModLmerTest]
## Formula: stage ~ week * treatment + tank + (1 | individual)
##    Data: tads05
## 
## REML criterion at convergence: 378.6
## 
## Scaled residuals: 
##     Min      1Q  Median      3Q     Max 
## -2.2728 -0.6280 -0.0838  0.5445  3.3588 
## 
## Random effects:
##  Groups     Name        Variance Std.Dev.
##  individual (Intercept) 0.07115  0.2667  
##  Residual               0.32570  0.5707  
## Number of obs: 197, groups:  individual, 54
## 
## Fixed effects:
##                    Estimate Std. Error        df t value Pr(>|t|)    
## (Intercept)        24.86459    0.29157  41.32975  85.278  < 2e-16 ***
## week                0.95758    0.05086 145.20412  18.827  < 2e-16 ***
## treatmentdhq        0.06627    0.15171 116.67859   0.437  0.66303    
## tankA07             1.37561    0.41138  42.16510   3.344  0.00174 ** 
## tankA09            -0.73359    0.40766  42.53161  -1.800  0.07904 .  
## tankA14             0.01641    0.40766  42.53161   0.040  0.96807    
## tankA15             0.18750    0.39061  35.89338   0.480  0.63413    
## tankB07             0.87500    0.30881  35.89338   2.833  0.00751 ** 
## tankB10             0.12500    0.39061  35.89338   0.320  0.75082    
## tankB15             0.62022    0.29745  36.59446   2.085  0.04409 *  
## tankB16             0.56250    0.39061  35.89338   1.440  0.15852    
## tankC07             0.53125    0.33828  35.89338   1.570  0.12509    
## tankC10             0.54909    0.32219  37.26156   1.704  0.09666 .  
## tankC11             0.67682    0.39871  38.70103   1.698  0.09763 .  
## tankC13             0.32838    0.34054  36.75022   0.964  0.34120    
## tankC15            -0.06250    0.39061  35.89338  -0.160  0.87377    
## week:treatmentdhq  -0.11212    0.07307 147.50880  -1.534  0.12708    
## ---
## Signif. codes:  0 '***' 0.001 '**' 0.01 '*' 0.05 '.' 0.1 ' ' 1
```

```
ggplot(tads05, aes(week, stage, colour = treatment, group = individual)) +
  geom_smooth(method=loess, aes(group = treatment, fill=treatment), size = 2) +
  theme_gdocs() + ggtitle('0.5% DHQ experiment') +
  scale_fill_manual(values = c("dhq" = "#dc4600", "control" = "#0064c8")) + 
  scale_color_manual(values = c("dhq" = "#dc4600", "control" = "#0064c8"))
```

```
## `geom_smooth()` using formula 'y ~ x'
```

```
lmer1 <- lmer(stage ~ week * treatment + tank + (1 | individual), tads10)
lmer2 <- lmer(stage ~ week * treatment + tank + (1 + treatment | individual), tads10)
# If P<0.5, the best fit is the one with the lowest logLik or lowest AIC/BIC, else, use lmer1
anova(lmer1, lmer2)
```

```
## Data: tads10
## Models:
## lmer1: stage ~ week * treatment + tank + (1 | individual)
## lmer2: stage ~ week * treatment + tank + (1 + treatment | individual)
##       npar    AIC    BIC  logLik deviance  Chisq Df Pr(>Chisq)  
## lmer1   14 803.44 849.61 -387.72   775.44                       
## lmer2   16 800.55 853.32 -384.27   768.55 6.8882  2    0.03193 *
## ---
## Signif. codes:  0 '***' 0.001 '**' 0.01 '*' 0.05 '.' 0.1 ' ' 1
```

```
# lmer3: same as lmer1 but removing tank effects
lmer3 <- lmer(stage ~ week * treatment + (1 + treatment | individual), tads10)
anova(lmer2, lmer3)
```

```
## Data: tads10
## Models:
## lmer3: stage ~ week * treatment + (1 + treatment | individual)
## lmer2: stage ~ week * treatment + tank + (1 + treatment | individual)
##       npar    AIC    BIC  logLik deviance  Chisq Df Pr(>Chisq)
## lmer3    8 795.91 822.30 -389.96   779.91                     
## lmer2   16 800.55 853.32 -384.27   768.55 11.365  8     0.1819
```

```
dev.lmer = lmer2
summary(dev.lmer)
```

```
## Linear mixed model fit by REML. t-tests use Satterthwaite's method [
## lmerModLmerTest]
## Formula: stage ~ week * treatment + tank + (1 + treatment | individual)
##    Data: tads10
## 
## REML criterion at convergence: 770.2
## 
## Scaled residuals: 
##     Min      1Q  Median      3Q     Max 
## -4.3874 -0.5273  0.0764  0.5949  1.7736 
## 
## Random effects:
##  Groups     Name         Variance Std.Dev. Corr 
##  individual (Intercept)  2.3504   1.533         
##             treatmentdhq 0.6084   0.780    -0.98
##  Residual                2.2687   1.506         
## Number of obs: 200, groups:  individual, 37
## 
## Fixed effects:
##                    Estimate Std. Error        df t value Pr(>|t|)    
## (Intercept)        23.67590    0.79179  17.24228  29.902 2.67e-16 ***
## week                1.98157    0.04989 178.22792  39.717  < 2e-16 ***
## treatmentdhq        0.12027    0.58286  45.76168   0.206   0.8374    
## tankA14             1.75687    1.04380   9.22470   1.683   0.1258    
## tankA15             2.69549    1.05943   8.28285   2.544   0.0336 *  
## tankB12             1.33922    1.04380   9.22470   1.283   0.2308    
## tankB14             1.41119    1.05688   9.00855   1.335   0.2146    
## tankB15             1.46209    0.97483  15.37024   1.500   0.1539    
## tankB16             2.47570    1.13742  14.91998   2.177   0.0460 *  
## tankC10             1.23573    0.94605  12.56417   1.306   0.2149    
## tankC15             1.52508    0.91087  10.29089   1.674   0.1241    
## week:treatmentdhq  -0.03355    0.09136 162.39059  -0.367   0.7139    
## ---
## Signif. codes:  0 '***' 0.001 '**' 0.01 '*' 0.05 '.' 0.1 ' ' 1
## 
## Correlation of Fixed Effects:
##             (Intr) week   trtmnt tnkA14 tnkA15 tnkB12 tnkB14 tnkB15 tnkB16
## week        -0.183                                                        
## treatmntdhq -0.307  0.344                                                 
## tankA14     -0.625 -0.082 -0.016                                          
## tankA15     -0.596 -0.074 -0.163  0.484                                   
## tankB12     -0.625 -0.082 -0.016  0.497  0.484                            
## tankB14     -0.629 -0.023 -0.061  0.482  0.488  0.482                     
## tankB15     -0.702 -0.054 -0.043  0.523  0.531  0.523  0.526              
## tankB16     -0.589 -0.047 -0.041  0.450  0.453  0.450  0.449  0.491       
## tankC10     -0.715 -0.073 -0.012  0.543  0.538  0.543  0.537  0.589  0.502
## tankC15     -0.732 -0.058 -0.039  0.563  0.561  0.563  0.558  0.609  0.520
## wk:trtmntdh  0.052 -0.543 -0.594  0.012  0.144  0.012  0.074  0.134  0.080
##             tnkC10 tnkC15
## week                     
## treatmntdhq              
## tankA14                  
## tankA15                  
## tankB12                  
## tankB14                  
## tankB15                  
## tankB16                  
## tankC10                  
## tankC15      0.624       
## wk:trtmntdh  0.079  0.070
## optimizer (nloptwrap) convergence code: 0 (OK)
## Model is nearly unidentifiable: large eigenvalue ratio
##  - Rescale variables?
```

```
ggplot(tads10, aes(week, stage, colour = treatment, group = individual)) +
  geom_smooth(method=loess, aes(group = treatment, fill=treatment), size = 2) +
  theme_gdocs() + ggtitle('1.0% DHQ experiment') + scale_x_discrete(limits = c(0,3,6,9)) +
  scale_fill_manual(values = c("dhq" = "#f0aa00", "control" = "#3caac8")) + 
  scale_color_manual(values = c("dhq" = "#f0aa00", "control" = "#3caac8"))
```

```
## `geom_smooth()` using formula 'y ~ x'
```

#### Body measurements

##### Plots

```
tads05.gather = gather(tads05, type, measurement, stl, svl, width, weight, tail, area)
ggplot(tads05.gather, aes(week, measurement, colour = treatment, group = individual)) +
  geom_smooth(method=loess, aes(group = treatment, fill=treatment), size = 2) +
  theme_gdocs() + ggtitle('0.5% DHQ experiment') +
  facet_wrap(~type, scales = "free_y") +
  scale_fill_manual(values = c("dhq" = "#dc4600", "control" = "#0064c8")) + 
  scale_color_manual(values = c("dhq" = "#dc4600", "control" = "#0064c8"))
```

```
tads10.gather = gather(tads10, type, measurement, stl, svl, width, weight, tail, area)
ggplot(tads10.gather, aes(week, measurement, colour = treatment, group = individual)) +
  geom_smooth(method=loess, aes(group = treatment, fill=treatment), size = 2) +
  theme_gdocs() + ggtitle('1.0% DHQ experiment') + scale_x_discrete(limits = c(0,3,6,9)) +
  facet_wrap(~type, scales = "free_y")  +
  scale_fill_manual(values = c("dhq" = "#f0aa00", "control" = "#3caac8")) + 
  scale_color_manual(values = c("dhq" = "#f0aa00", "control" = "#3caac8"))
```

as you can notice, we do not have data for week 10 in the DHQ 1.0% experiment, hence, we subset this dataset up to week 9 for the statistical analyses.

```
tads10 = subset(tads10, week <= 9)
```

##### Statistics for Weight

```
# NOTE: in Shapiro tests, P > 0.5 means the distribution is normal
desc = Summarize(weight ~ treatment + week, data=tads05)
kbl(desc, caption = "Descriptive statistics for weight in 0.5% Experiment") %>% kable_paper("hover", full_width = F)
```

Descriptive statistics for weight in 0.5% Experiment

| treatment | week | n | nvalid | mean | sd | min | Q1 | median | Q3 | max |
| --- | --- | --- | --- | --- | --- | --- | --- | --- | --- | --- |
| control | 0 | 27 | 27 | 23.84444 | 3.759842 | 17.3 | 20.700 | 24.00 | 26.000 | 33.7 |
| dhq | 0 | 27 | 27 | 24.46667 | 5.560575 | 15.7 | 21.000 | 24.30 | 26.500 | 36.7 |
| control | 1 | 27 | 26 | 38.16154 | 7.483640 | 26.3 | 31.775 | 38.65 | 42.825 | 52.3 |
| dhq | 1 | 27 | 26 | 37.05769 | 8.215117 | 22.7 | 32.000 | 35.50 | 42.925 | 55.7 |
| control | 2 | 27 | 27 | 47.30370 | 11.210074 | 26.7 | 40.300 | 45.00 | 55.000 | 75.0 |
| dhq | 2 | 27 | 24 | 45.15000 | 9.409662 | 27.3 | 38.050 | 43.50 | 51.850 | 62.7 |
| control | 3 | 27 | 24 | 54.17083 | 14.705854 | 29.3 | 44.750 | 52.70 | 67.075 | 84.7 |
| dhq | 3 | 27 | 22 | 49.20909 | 10.722158 | 30.3 | 41.425 | 49.15 | 55.025 | 73.3 |

```
ggplot(tads05, aes(weight, fill = treatment)) + 
  geom_density(alpha = 0.5) + facet_wrap(~week) + ggtitle('0.5% DHQ experiment') +
  scale_fill_manual(values = c("dhq" = "#dc4600", "control" = "#0064c8")) + 
  scale_color_manual(values = c("dhq" = "#dc4600", "control" = "#0064c8"))
```

```
tads05 %>% group_by(treatment, week) %>% shapiro_test(weight)
```

```
## # A tibble: 8 x 5
##   treatment  week variable statistic     p
##   <chr>     <int> <chr>        <dbl> <dbl>
## 1 control       0 weight       0.974 0.718
## 2 control       1 weight       0.940 0.132
## 3 control       2 weight       0.978 0.816
## 4 control       3 weight       0.968 0.618
## 5 dhq           0 weight       0.966 0.510
## 6 dhq           1 weight       0.983 0.935
## 7 dhq           2 weight       0.976 0.805
## 8 dhq           3 weight       0.986 0.980
```

```
lmer1 <- lmer(weight ~ week * treatment + tank + (1 | individual), tads05)
lmer2 <- lmer(weight ~ week * treatment + (1 | individual), tads05)
anova(lmer1, lmer2)
```

```
## Data: tads05
## Models:
## lmer2: weight ~ week * treatment + (1 | individual)
## lmer1: weight ~ week * treatment + tank + (1 | individual)
##       npar    AIC    BIC  logLik deviance  Chisq Df Pr(>Chisq)    
## lmer2    6 1444.4 1464.3 -716.22   1432.4                         
## lmer1   19 1418.0 1481.0 -690.00   1380.0 52.428 13  1.136e-06 ***
## ---
## Signif. codes:  0 '***' 0.001 '**' 0.01 '*' 0.05 '.' 0.1 ' ' 1
```

```
weight.lmer = lmer1
summary(weight.lmer)
```

```
## Linear mixed model fit by REML. t-tests use Satterthwaite's method [
## lmerModLmerTest]
## Formula: weight ~ week * treatment + tank + (1 | individual)
##    Data: tads05
## 
## REML criterion at convergence: 1321
## 
## Scaled residuals: 
##      Min       1Q   Median       3Q      Max 
## -2.70975 -0.57796  0.03482  0.47973  2.31242 
## 
## Random effects:
##  Groups     Name        Variance Std.Dev.
##  individual (Intercept) 16.24    4.029   
##  Residual               46.22    6.798   
## Number of obs: 203, groups:  individual, 54
## 
## Fixed effects:
##                   Estimate Std. Error       df t value Pr(>|t|)    
## (Intercept)        18.2563     3.9021  42.7054   4.679 2.91e-05 ***
## week               10.1307     0.6045 150.7059  16.760  < 2e-16 ***
## treatmentdhq        0.7336     1.9107 103.4841   0.384  0.70181    
## tankA07             6.1817     5.5016  44.0237   1.124  0.26727    
## tankA09            -8.0169     5.4517  43.7583  -1.471  0.14858    
## tankA14            10.4125     5.2717  38.3287   1.975  0.05548 .  
## tankA15             8.1750     5.2717  38.3287   1.551  0.12918    
## tankB07             2.2781     4.1676  38.3287   0.547  0.58781    
## tankB10            11.9000     5.2717  38.3287   2.257  0.02976 *  
## tankB15             4.3037     4.0019  38.8456   1.075  0.28882    
## tankB16            14.9125     5.2717  38.3287   2.829  0.00739 ** 
## tankC07            10.5938     4.5654  38.3287   2.320  0.02574 *  
## tankC10            14.8847     4.3157  38.6918   3.449  0.00137 ** 
## tankC11             3.5147     5.3585  40.7227   0.656  0.51557    
## tankC13            14.4933     4.5898  39.0728   3.158  0.00306 ** 
## tankC15            15.5250     5.2717  38.3287   2.945  0.00546 ** 
## week:treatmentdhq  -1.8089     0.8698 152.8680  -2.080  0.03923 *  
## ---
## Signif. codes:  0 '***' 0.001 '**' 0.01 '*' 0.05 '.' 0.1 ' ' 1
```

```
desc = Summarize(weight ~ treatment + week, data=tads10)
kbl(desc, caption = "Descriptive statistics for weight in 1.0% Experiment") %>% kable_paper("hover", full_width = F)
```

Descriptive statistics for weight in 1.0% Experiment

| treatment | week | n | nvalid | mean | sd | min | Q1 | median | Q3 | max |
| --- | --- | --- | --- | --- | --- | --- | --- | --- | --- | --- |
| control | 0 | 24 | 23 | 26.44783 | 9.863702 | 16.3 | 19.000 | 22.7 | 29.250 | 49.7 |
| dhq | 0 | 13 | 11 | 29.34545 | 16.937199 | 14.0 | 20.650 | 23.0 | 32.150 | 73.0 |
| control | 1 | 24 | 15 | 45.90667 | 18.737909 | 15.7 | 34.500 | 38.3 | 58.500 | 90.3 |
| dhq | 1 | 13 | 7 | 59.31429 | 29.304233 | 24.5 | 41.350 | 50.7 | 72.150 | 113.0 |
| control | 2 | 24 | 21 | 72.64762 | 28.991716 | 22.3 | 62.700 | 68.7 | 84.300 | 165.3 |
| dhq | 2 | 13 | 12 | 80.91667 | 27.988564 | 41.3 | 57.750 | 74.4 | 96.450 | 132.5 |
| control | 3 | 24 | 20 | 96.76500 | 23.000281 | 44.8 | 84.875 | 94.5 | 109.250 | 155.0 |
| dhq | 3 | 13 | 10 | 97.97000 | 36.164594 | 50.3 | 72.000 | 86.8 | 120.375 | 162.3 |
| control | 4 | 24 | 19 | 128.61579 | 23.016426 | 79.3 | 113.800 | 130.5 | 143.300 | 183.0 |
| dhq | 4 | 13 | 9 | 126.54444 | 31.391882 | 90.0 | 105.500 | 124.0 | 143.500 | 193.8 |
| control | 5 | 24 | 19 | 143.78421 | 23.550189 | 87.3 | 137.250 | 141.5 | 151.000 | 190.7 |
| dhq | 5 | 13 | 8 | 138.67500 | 22.585631 | 109.3 | 122.850 | 139.0 | 147.975 | 178.0 |
| control | 6 | 24 | 17 | 169.01176 | 17.688446 | 134.8 | 160.800 | 168.0 | 179.500 | 202.3 |
| dhq | 6 | 13 | 7 | 165.52857 | 21.852133 | 140.0 | 145.000 | 174.3 | 180.200 | 194.0 |
| control | 7 | 24 | 13 | 182.03846 | 14.994418 | 159.3 | 170.700 | 181.0 | 193.000 | 205.3 |
| dhq | 7 | 13 | 5 | 173.60000 | 21.773723 | 149.0 | 157.700 | 169.0 | 195.800 | 196.5 |
| control | 8 | 24 | 12 | 171.10000 | 21.683467 | 146.5 | 152.525 | 163.9 | 186.475 | 207.3 |
| dhq | 8 | 13 | 5 | 175.66000 | 24.547362 | 156.7 | 158.300 | 159.0 | 196.300 | 208.0 |
| control | 9 | 24 | 8 | 141.57500 | 41.383459 | 103.5 | 109.175 | 129.7 | 158.900 | 204.6 |
| dhq | 9 | 13 | 3 | 179.36667 | 10.789501 | 170.3 | 173.400 | 176.5 | 183.900 | 191.3 |

```
ggplot(tads10, aes(weight, fill = treatment)) + 
  geom_density(alpha = 0.5) + facet_wrap(~week) + ggtitle('1.0% DHQ experiment')  +
  scale_fill_manual(values = c("dhq" = "#f0aa00", "control" = "#3caac8")) + 
  scale_color_manual(values = c("dhq" = "#f0aa00", "control" = "#3caac8"))
```

```
tads10 %>% group_by(treatment, week) %>% shapiro_test(weight)
```

```
## # A tibble: 20 x 5
##    treatment  week variable statistic       p
##    <chr>     <int> <chr>        <dbl>   <dbl>
##  1 control       0 weight       0.839 0.00172
##  2 control       1 weight       0.937 0.341  
##  3 control       2 weight       0.880 0.0148 
##  4 control       3 weight       0.963 0.605  
##  5 control       4 weight       0.982 0.959  
##  6 control       5 weight       0.936 0.221  
##  7 control       6 weight       0.989 0.999  
##  8 control       7 weight       0.957 0.708  
##  9 control       8 weight       0.906 0.188  
## 10 control       9 weight       0.815 0.0410 
## 11 dhq           0 weight       0.750 0.00206
## 12 dhq           1 weight       0.932 0.571  
## 13 dhq           2 weight       0.948 0.611  
## 14 dhq           3 weight       0.949 0.661  
## 15 dhq           4 weight       0.907 0.294  
## 16 dhq           5 weight       0.965 0.859  
## 17 dhq           6 weight       0.888 0.266  
## 18 dhq           7 weight       0.876 0.293  
## 19 dhq           8 weight       0.777 0.0516 
## 20 dhq           9 weight       0.947 0.557
```

```
lmer1 <- lmer(weight ~ week * treatment + tank + (1 | individual), tads10)
lmer2 <- lmer(weight ~ week * treatment + (1 | individual), tads10)
anova(lmer1, lmer2)
```

```
## Data: tads10
## Models:
## lmer2: weight ~ week * treatment + (1 | individual)
## lmer1: weight ~ week * treatment + tank + (1 | individual)
##       npar    AIC    BIC  logLik deviance  Chisq Df Pr(>Chisq)
## lmer2    6 2295.9 2316.9 -1142.0   2283.9                     
## lmer1   14 2304.3 2353.3 -1138.2   2276.3 7.5867  8     0.4748
```

```
weight.lmer = lmer2
summary(weight.lmer)
```

```
## Linear mixed model fit by REML. t-tests use Satterthwaite's method [
## lmerModLmerTest]
## Formula: weight ~ week * treatment + (1 | individual)
##    Data: tads10
## 
## REML criterion at convergence: 2270.8
## 
## Scaled residuals: 
##     Min      1Q  Median      3Q     Max 
## -4.3494 -0.4776  0.0184  0.5895  2.7862 
## 
## Random effects:
##  Groups     Name        Variance Std.Dev.
##  individual (Intercept) 277.7    16.66   
##  Residual               562.5    23.72   
## Number of obs: 244, groups:  individual, 35
## 
## Fixed effects:
##                   Estimate Std. Error       df t value Pr(>|t|)    
## (Intercept)        38.6154     4.7918  59.0268   8.059 4.34e-11 ***
## week               17.7625     0.7280 223.9360  24.398  < 2e-16 ***
## treatmentdhq       -0.5042     8.2861  61.9143  -0.061   0.9517    
## week:treatmentdhq   2.2930     1.3540 230.1722   1.694   0.0917 .  
## ---
## Signif. codes:  0 '***' 0.001 '**' 0.01 '*' 0.05 '.' 0.1 ' ' 1
## 
## Correlation of Fixed Effects:
##             (Intr) week   trtmnt
## week        -0.548              
## treatmntdhq -0.578  0.317       
## wk:trtmntdh  0.295 -0.538 -0.555
```

##### Statistics for STL

```
desc = Summarize(stl ~ treatment + week, data=tads05)
kbl(desc, caption = "Descriptive statistics for STL in 0.5% Experiment") %>% kable_paper("hover", full_width = F)
```

Descriptive statistics for STL in 0.5% Experiment

| treatment | week | n | nvalid | mean | sd | min | Q1 | median | Q3 | max |
| --- | --- | --- | --- | --- | --- | --- | --- | --- | --- | --- |
| control | 0 | 27 | 27 | 11.22963 | 0.6910498 | 9.8 | 10.850 | 11.20 | 11.500 | 13.4 |
| dhq | 0 | 27 | 27 | 11.38148 | 1.0532138 | 9.5 | 10.700 | 11.40 | 11.850 | 13.9 |
| control | 1 | 27 | 26 | 13.65385 | 1.1710613 | 11.6 | 12.925 | 13.50 | 14.300 | 16.1 |
| dhq | 1 | 27 | 26 | 13.67692 | 1.0226662 | 11.7 | 13.000 | 13.70 | 14.675 | 15.5 |
| control | 2 | 27 | 27 | 14.60370 | 1.1853210 | 13.0 | 13.850 | 14.50 | 15.000 | 17.5 |
| dhq | 2 | 27 | 24 | 14.44583 | 1.4613809 | 11.3 | 13.500 | 14.35 | 15.400 | 17.7 |
| control | 3 | 27 | 23 | 15.03043 | 1.5921497 | 12.6 | 13.700 | 15.10 | 16.500 | 17.4 |
| dhq | 3 | 27 | 22 | 14.92727 | 1.3617852 | 11.8 | 14.125 | 14.60 | 15.825 | 18.3 |

```
ggplot(tads05, aes(stl, fill = treatment)) + 
  geom_density(alpha = 0.5) + facet_wrap(~week) + ggtitle('0.5% DHQ experiment') +
  scale_fill_manual(values = c("dhq" = "#dc4600", "control" = "#0064c8")) + 
  scale_color_manual(values = c("dhq" = "#dc4600", "control" = "#0064c8"))
```

```
tads05 %>% group_by(treatment, week) %>% shapiro_test(stl)
```

```
## # A tibble: 8 x 5
##   treatment  week variable statistic      p
##   <chr>     <int> <chr>        <dbl>  <dbl>
## 1 control       0 stl          0.939 0.117 
## 2 control       1 stl          0.975 0.748 
## 3 control       2 stl          0.907 0.0199
## 4 control       3 stl          0.930 0.110 
## 5 dhq           0 stl          0.979 0.837 
## 6 dhq           1 stl          0.964 0.465 
## 7 dhq           2 stl          0.971 0.690 
## 8 dhq           3 stl          0.967 0.648
```

```
lmer1 <- lmer(stl ~ week * treatment + tank + (1 | individual), tads05)
lmer2 <- lmer(stl ~ week * treatment + (1 | individual), tads05)
anova(lmer1, lmer2)
```

```
## Data: tads05
## Models:
## lmer2: stl ~ week * treatment + (1 | individual)
## lmer1: stl ~ week * treatment + tank + (1 | individual)
##       npar    AIC    BIC  logLik deviance  Chisq Df Pr(>Chisq)    
## lmer2    6 665.23 685.08 -326.62   653.23                         
## lmer1   19 651.12 713.97 -306.56   613.12 40.116 13  0.0001324 ***
## ---
## Signif. codes:  0 '***' 0.001 '**' 0.01 '*' 0.05 '.' 0.1 ' ' 1
```

```
stl.lmer = lmer1
summary(stl.lmer)
```

```
## Linear mixed model fit by REML. t-tests use Satterthwaite's method [
## lmerModLmerTest]
## Formula: stl ~ week * treatment + tank + (1 | individual)
##    Data: tads05
## 
## REML criterion at convergence: 621
## 
## Scaled residuals: 
##      Min       1Q   Median       3Q      Max 
## -2.86707 -0.60852  0.01512  0.63122  2.23043 
## 
## Random effects:
##  Groups     Name        Variance Std.Dev.
##  individual (Intercept) 0.2289   0.4785  
##  Residual               1.1582   1.0762  
## Number of obs: 202, groups:  individual, 54
## 
## Fixed effects:
##                    Estimate Std. Error        df t value Pr(>|t|)    
## (Intercept)        10.98544    0.53805  42.21852  20.417  < 2e-16 ***
## week                1.28555    0.09663 150.01308  13.304  < 2e-16 ***
## treatmentdhq        0.12095    0.27991 120.00703   0.432  0.66645    
## tankA07             0.29653    0.75969  43.02986   0.390  0.69822    
## tankA09            -1.45879    0.75291  43.56416  -1.938  0.05918 .  
## tankA14             0.75000    0.72006  36.49096   1.042  0.30446    
## tankA15             0.77500    0.72006  36.49096   1.076  0.28886    
## tankB07             0.25312    0.56926  36.49096   0.445  0.65919    
## tankB10             1.27500    0.72006  36.49096   1.771  0.08497 .  
## tankB15             0.53506    0.54721  37.07837   0.978  0.33451    
## tankB16             1.05000    0.72006  36.49096   1.458  0.15334    
## tankC07             1.12500    0.62359  36.49096   1.804  0.07948 .  
## tankC10             1.75841    0.58995  36.92835   2.981  0.00507 ** 
## tankC11             0.53935    0.73562  39.45858   0.733  0.46777    
## tankC13             1.18983    0.63258  38.44765   1.881  0.06757 .  
## tankC15             1.68750    0.72006  36.49096   2.344  0.02466 *  
## week:treatmentdhq  -0.11564    0.13808 152.26051  -0.837  0.40363    
## ---
## Signif. codes:  0 '***' 0.001 '**' 0.01 '*' 0.05 '.' 0.1 ' ' 1
```

```
desc = Summarize(stl ~ treatment + week, data=tads10)
kbl(desc, caption = "Descriptive statistics for STL in 1.0% Experiment") %>% kable_paper("hover", full_width = F)
```

Descriptive statistics for STL in 1.0% Experiment

| treatment | week | n | nvalid | mean | sd | min | Q1 | median | Q3 | max |
| --- | --- | --- | --- | --- | --- | --- | --- | --- | --- | --- |
| control | 0 | 24 | 24 | 11.26667 | 1.9771888 | 8.6 | 10.075 | 11.05 | 12.250 | 16.1 |
| dhq | 0 | 13 | 13 | 12.20769 | 2.6399058 | 9.4 | 10.200 | 12.10 | 12.800 | 18.9 |
| control | 1 | 24 | 15 | 12.57333 | 1.8652716 | 9.7 | 11.200 | 12.80 | 13.500 | 17.0 |
| dhq | 1 | 13 | 8 | 13.55000 | 2.3603874 | 10.8 | 11.875 | 13.30 | 15.175 | 17.2 |
| control | 2 | 24 | 21 | 15.37143 | 2.7495714 | 9.2 | 13.500 | 15.10 | 17.000 | 20.0 |
| dhq | 2 | 13 | 12 | 16.05000 | 2.8937236 | 12.2 | 14.275 | 15.60 | 16.900 | 22.6 |
| control | 3 | 24 | 19 | 17.06316 | 2.9456579 | 11.3 | 15.450 | 16.30 | 18.700 | 22.2 |
| dhq | 3 | 13 | 10 | 18.66000 | 3.7689374 | 14.5 | 15.700 | 17.40 | 21.450 | 25.8 |
| control | 4 | 24 | 19 | 20.07895 | 2.6889855 | 15.7 | 17.300 | 20.80 | 22.350 | 23.5 |
| dhq | 4 | 13 | 9 | 19.64444 | 3.4641417 | 16.2 | 17.100 | 18.40 | 22.000 | 26.6 |
| control | 5 | 24 | 19 | 22.47895 | 1.2817431 | 20.7 | 21.750 | 22.10 | 23.250 | 25.3 |
| dhq | 5 | 13 | 8 | 22.80000 | 1.5629642 | 21.4 | 21.850 | 22.05 | 23.400 | 25.9 |
| control | 6 | 24 | 17 | 22.38824 | 2.8718205 | 16.8 | 22.000 | 22.90 | 24.000 | 26.3 |
| dhq | 6 | 13 | 7 | 22.68571 | 2.7637104 | 18.2 | 21.200 | 23.30 | 24.800 | 25.3 |
| control | 7 | 24 | 13 | 24.25385 | 2.4144065 | 17.2 | 24.000 | 24.70 | 25.800 | 27.1 |
| dhq | 7 | 13 | 5 | 25.44000 | 0.9423375 | 24.3 | 24.600 | 25.70 | 26.300 | 26.3 |
| control | 8 | 24 | 12 | 20.51667 | 4.0505518 | 11.9 | 18.525 | 20.10 | 24.050 | 25.5 |
| dhq | 8 | 13 | 5 | 23.64000 | 2.0305172 | 20.4 | 23.300 | 23.80 | 25.300 | 25.4 |
| control | 9 | 24 | 6 | 16.30000 | 5.2199617 | 9.2 | 12.925 | 16.60 | 20.275 | 22.2 |
| dhq | 9 | 13 | 3 | 18.96667 | 0.4163332 | 18.5 | 18.800 | 19.10 | 19.200 | 19.3 |

```
ggplot(tads10, aes(stl, fill = treatment)) + 
  geom_density(alpha = 0.5) + facet_wrap(~week) + ggtitle('1.0% DHQ experiment') +
  scale_fill_manual(values = c("dhq" = "#f0aa00", "control" = "#3caac8")) + 
  scale_color_manual(values = c("dhq" = "#f0aa00", "control" = "#3caac8"))
```

```
tads10 %>% group_by(treatment, week) %>% shapiro_test(weight)
```

```
## # A tibble: 20 x 5
##    treatment  week variable statistic       p
##    <chr>     <int> <chr>        <dbl>   <dbl>
##  1 control       0 weight       0.839 0.00172
##  2 control       1 weight       0.937 0.341  
##  3 control       2 weight       0.880 0.0148 
##  4 control       3 weight       0.963 0.605  
##  5 control       4 weight       0.982 0.959  
##  6 control       5 weight       0.936 0.221  
##  7 control       6 weight       0.989 0.999  
##  8 control       7 weight       0.957 0.708  
##  9 control       8 weight       0.906 0.188  
## 10 control       9 weight       0.815 0.0410 
## 11 dhq           0 weight       0.750 0.00206
## 12 dhq           1 weight       0.932 0.571  
## 13 dhq           2 weight       0.948 0.611  
## 14 dhq           3 weight       0.949 0.661  
## 15 dhq           4 weight       0.907 0.294  
## 16 dhq           5 weight       0.965 0.859  
## 17 dhq           6 weight       0.888 0.266  
## 18 dhq           7 weight       0.876 0.293  
## 19 dhq           8 weight       0.777 0.0516 
## 20 dhq           9 weight       0.947 0.557
```

```
tads10 %>% group_by(treatment, week) %>% shapiro_test(stl)
```

```
## # A tibble: 20 x 5
##    treatment  week variable statistic       p
##    <chr>     <int> <chr>        <dbl>   <dbl>
##  1 control       0 stl          0.912 0.0387 
##  2 control       1 stl          0.936 0.330  
##  3 control       2 stl          0.980 0.922  
##  4 control       3 stl          0.966 0.698  
##  5 control       4 stl          0.868 0.0133 
##  6 control       5 stl          0.944 0.316  
##  7 control       6 stl          0.900 0.0691 
##  8 control       7 stl          0.778 0.00375
##  9 control       8 stl          0.919 0.278  
## 10 control       9 stl          0.900 0.375  
## 11 dhq           0 stl          0.878 0.0660 
## 12 dhq           1 stl          0.925 0.469  
## 13 dhq           2 stl          0.935 0.434  
## 14 dhq           3 stl          0.903 0.235  
## 15 dhq           4 stl          0.889 0.195  
## 16 dhq           5 stl          0.832 0.0625 
## 17 dhq           6 stl          0.866 0.170  
## 18 dhq           7 stl          0.852 0.201  
## 19 dhq           8 stl          0.878 0.302  
## 20 dhq           9 stl          0.923 0.463
```

```
lmer1 <- lmer(stl ~ week * treatment + tank + (1 | individual), tads10)
lmer2 <- lmer(stl ~ week * treatment + (1 | individual), tads10)
anova(lmer1, lmer2)
```

```
## Data: tads10
## Models:
## lmer2: stl ~ week * treatment + (1 | individual)
## lmer1: stl ~ week * treatment + tank + (1 | individual)
##       npar    AIC    BIC  logLik deviance Chisq Df Pr(>Chisq)
## lmer2    6 1312.6 1333.6 -650.31   1300.6                    
## lmer1   14 1325.3 1374.3 -648.63   1297.3 3.346  8     0.9108
```

```
stl.lmer = lmer2
summary(stl.lmer)
```

```
## Linear mixed model fit by REML. t-tests use Satterthwaite's method [
## lmerModLmerTest]
## Formula: stl ~ week * treatment + (1 | individual)
##    Data: tads10
## 
## REML criterion at convergence: 1306.2
## 
## Scaled residuals: 
##     Min      1Q  Median      3Q     Max 
## -4.5262 -0.6555  0.0717  0.7260  2.3720 
## 
## Random effects:
##  Groups     Name        Variance Std.Dev.
##  individual (Intercept)  0.00    0.000   
##  Residual               12.03    3.468   
## Number of obs: 245, groups:  individual, 37
## 
## Fixed effects:
##                    Estimate Std. Error        df t value Pr(>|t|)    
## (Intercept)        12.78304    0.47209 241.00000  27.078   <2e-16 ***
## week                1.34602    0.10175 241.00000  13.229   <2e-16 ***
## treatmentdhq        0.45338    0.80127 241.00000   0.566    0.572    
## week:treatmentdhq   0.09968    0.17885 241.00000   0.557    0.578    
## ---
## Signif. codes:  0 '***' 0.001 '**' 0.01 '*' 0.05 '.' 0.1 ' ' 1
## 
## Correlation of Fixed Effects:
##             (Intr) week   trtmnt
## week        -0.820              
## treatmntdhq -0.589  0.483       
## wk:trtmntdh  0.467 -0.569 -0.807
## optimizer (nloptwrap) convergence code: 0 (OK)
## boundary (singular) fit: see ?isSingular
```

##### Statistics for SVL

```
desc = Summarize(svl ~ treatment + week, data=tads05)
kbl(desc, caption = "Descriptive statistics for SVL in 0.5% Experiment") %>% kable_paper("hover", full_width = F)
```

Descriptive statistics for SVL in 0.5% Experiment

| treatment | week | n | nvalid | mean | sd | min | Q1 | median | Q3 | max |
| --- | --- | --- | --- | --- | --- | --- | --- | --- | --- | --- |
| control | 0 | 27 | 27 | 4.211111 | 0.3651484 | 3.5 | 4.000 | 4.20 | 4.500 | 4.9 |
| dhq | 0 | 27 | 27 | 4.159259 | 0.3785186 | 3.6 | 3.900 | 4.10 | 4.300 | 5.0 |
| control | 1 | 27 | 26 | 4.953846 | 0.5069365 | 4.0 | 4.700 | 5.00 | 5.275 | 5.9 |
| dhq | 1 | 27 | 26 | 4.873077 | 0.5040299 | 3.5 | 4.600 | 5.00 | 5.200 | 5.8 |
| control | 2 | 27 | 27 | 5.362963 | 0.5492289 | 4.5 | 4.800 | 5.40 | 5.800 | 6.4 |
| dhq | 2 | 27 | 24 | 5.295833 | 0.5384997 | 4.5 | 4.775 | 5.35 | 5.700 | 6.4 |
| control | 3 | 27 | 23 | 5.730435 | 0.6255432 | 4.5 | 5.350 | 5.60 | 6.200 | 7.0 |
| dhq | 3 | 27 | 22 | 5.572727 | 0.5128648 | 4.8 | 5.125 | 5.65 | 6.000 | 6.4 |

```
ggplot(tads05, aes(svl, fill = treatment)) + 
  geom_density(alpha = 0.5) + facet_wrap(~week) + ggtitle('0.5% DHQ experiment') +
  scale_fill_manual(values = c("dhq" = "#dc4600", "control" = "#0064c8")) + 
  scale_color_manual(values = c("dhq" = "#dc4600", "control" = "#0064c8"))
```

```
tads05 %>% group_by(treatment, week) %>% shapiro_test(svl)
```

```
## # A tibble: 8 x 5
##   treatment  week variable statistic     p
##   <chr>     <int> <chr>        <dbl> <dbl>
## 1 control       0 svl          0.968 0.539
## 2 control       1 svl          0.976 0.780
## 3 control       2 svl          0.947 0.178
## 4 control       3 svl          0.982 0.934
## 5 dhq           0 svl          0.939 0.116
## 6 dhq           1 svl          0.947 0.200
## 7 dhq           2 svl          0.953 0.318
## 8 dhq           3 svl          0.932 0.136
```

```
lmer1 <- lmer(svl ~ week * treatment + tank + (1 | individual), tads05)
lmer2 <- lmer(svl ~ week * treatment + (1 | individual), tads05)
anova(lmer1, lmer2)
```

```
## Data: tads05
## Models:
## lmer2: svl ~ week * treatment + (1 | individual)
## lmer1: svl ~ week * treatment + tank + (1 | individual)
##       npar    AIC    BIC  logLik deviance  Chisq Df Pr(>Chisq)    
## lmer2    6 282.83 302.68 -135.41   270.83                         
## lmer1   19 261.20 324.06 -111.60   223.20 47.625 13  7.574e-06 ***
## ---
## Signif. codes:  0 '***' 0.001 '**' 0.01 '*' 0.05 '.' 0.1 ' ' 1
```

```
svl.lmer = lmer1
summary(svl.lmer)
```

```
## Linear mixed model fit by REML. t-tests use Satterthwaite's method [
## lmerModLmerTest]
## Formula: svl ~ week * treatment + tank + (1 | individual)
##    Data: tads05
## 
## REML criterion at convergence: 265
## 
## Scaled residuals: 
##      Min       1Q   Median       3Q      Max 
## -2.66643 -0.56847 -0.00957  0.69203  1.98225 
## 
## Random effects:
##  Groups     Name        Variance Std.Dev.
##  individual (Intercept) 0.0266   0.1631  
##  Residual               0.1725   0.4154  
## Number of obs: 202, groups:  individual, 54
## 
## Fixed effects:
##                    Estimate Std. Error        df t value Pr(>|t|)    
## (Intercept)         3.83310    0.19807  41.99303  19.352  < 2e-16 ***
## week                0.50986    0.03728 149.80551  13.678  < 2e-16 ***
## treatmentdhq       -0.04243    0.10542 126.20302  -0.402  0.68801    
## tankA07             0.61982    0.27977  42.53398   2.215  0.03213 *  
## tankA09            -0.01535    0.27738  43.48123  -0.055  0.95613    
## tankA14             0.77500    0.26407  35.74356   2.935  0.00580 ** 
## tankA15             0.66250    0.26407  35.74356   2.509  0.01679 *  
## tankB07             0.11562    0.20876  35.74356   0.554  0.58313    
## tankB10             0.62500    0.26407  35.74356   2.367  0.02348 *  
## tankB15             0.35424    0.20076  36.35214   1.765  0.08605 .  
## tankB16             0.60000    0.26407  35.74356   2.272  0.02920 *  
## tankC07             0.72500    0.22869  35.74356   3.170  0.00312 ** 
## tankC10             0.85628    0.21642  36.20660   3.957  0.00034 ***
## tankC11             0.53190    0.27033  38.92180   1.968  0.05627 .  
## tankC13             0.53635    0.23230  37.83683   2.309  0.02651 *  
## tankC15             0.86250    0.26407  35.74356   3.266  0.00241 ** 
## week:treatmentdhq  -0.03555    0.05325 152.15909  -0.668  0.50533    
## ---
## Signif. codes:  0 '***' 0.001 '**' 0.01 '*' 0.05 '.' 0.1 ' ' 1
```

```
desc = Summarize(svl ~ treatment + week, data=tads10)
kbl(desc, caption = "Descriptive statistics for SVL in 1.0% Experiment") %>% kable_paper("hover", full_width = F)
```

Descriptive statistics for SVL in 1.0% Experiment

| treatment | week | n | nvalid | mean | sd | min | Q1 | median | Q3 | max |
| --- | --- | --- | --- | --- | --- | --- | --- | --- | --- | --- |
| control | 0 | 24 | 24 | 4.220833 | 0.6775783 | 3.2 | 3.800 | 4.20 | 4.525 | 6.0 |
| dhq | 0 | 13 | 13 | 4.538462 | 0.9412212 | 3.6 | 4.100 | 4.20 | 4.700 | 7.1 |
| control | 1 | 24 | 15 | 5.026667 | 0.9512899 | 3.8 | 4.500 | 4.80 | 5.200 | 7.6 |
| dhq | 1 | 13 | 8 | 4.962500 | 0.9560895 | 3.4 | 4.575 | 4.85 | 5.250 | 6.7 |
| control | 2 | 24 | 21 | 5.714286 | 1.1363475 | 3.2 | 5.000 | 5.60 | 6.500 | 7.9 |
| dhq | 2 | 13 | 12 | 6.100000 | 1.1321741 | 4.8 | 5.375 | 5.70 | 6.850 | 8.6 |
| control | 3 | 24 | 19 | 6.452632 | 1.0404958 | 4.6 | 5.800 | 6.40 | 7.000 | 8.5 |
| dhq | 3 | 13 | 10 | 7.020000 | 1.4649991 | 5.5 | 5.825 | 6.20 | 8.650 | 8.8 |
| control | 4 | 24 | 19 | 7.800000 | 0.9018500 | 6.5 | 6.900 | 7.90 | 8.450 | 9.2 |
| dhq | 4 | 13 | 9 | 7.544444 | 1.2923923 | 6.4 | 6.600 | 6.80 | 9.000 | 9.6 |
| control | 5 | 24 | 19 | 8.668421 | 0.5426812 | 7.8 | 8.350 | 8.60 | 8.850 | 9.8 |
| dhq | 5 | 13 | 8 | 8.975000 | 0.6584614 | 7.9 | 8.550 | 9.10 | 9.275 | 10.0 |
| control | 6 | 24 | 17 | 8.617647 | 1.0560749 | 6.8 | 8.100 | 8.80 | 9.200 | 10.5 |
| dhq | 6 | 13 | 7 | 8.800000 | 0.9504385 | 7.5 | 8.150 | 9.10 | 9.400 | 9.9 |
| control | 7 | 24 | 13 | 9.453846 | 0.9820178 | 6.7 | 9.100 | 9.60 | 9.900 | 10.7 |
| dhq | 7 | 13 | 5 | 9.680000 | 0.3346640 | 9.1 | 9.700 | 9.80 | 9.900 | 9.9 |
| control | 8 | 24 | 12 | 8.908333 | 1.1835681 | 7.0 | 7.800 | 9.20 | 9.875 | 10.3 |
| dhq | 8 | 13 | 5 | 9.220000 | 1.2316655 | 7.7 | 8.600 | 8.90 | 10.100 | 10.8 |
| control | 9 | 24 | 8 | 8.900000 | 2.1895857 | 7.1 | 7.475 | 8.15 | 9.525 | 13.8 |
| dhq | 9 | 13 | 3 | 8.066667 | 0.4509250 | 7.6 | 7.850 | 8.10 | 8.300 | 8.5 |

```
ggplot(tads10, aes(svl, fill = treatment)) + 
  geom_density(alpha = 0.5) + facet_wrap(~week) + ggtitle('1.0% DHQ experiment') +
  scale_fill_manual(values = c("dhq" = "#f0aa00", "control" = "#3caac8")) + 
  scale_color_manual(values = c("dhq" = "#f0aa00", "control" = "#3caac8"))
```

```
tads10 %>% group_by(treatment, week) %>% shapiro_test(weight)
```

```
## # A tibble: 20 x 5
##    treatment  week variable statistic       p
##    <chr>     <int> <chr>        <dbl>   <dbl>
##  1 control       0 weight       0.839 0.00172
##  2 control       1 weight       0.937 0.341  
##  3 control       2 weight       0.880 0.0148 
##  4 control       3 weight       0.963 0.605  
##  5 control       4 weight       0.982 0.959  
##  6 control       5 weight       0.936 0.221  
##  7 control       6 weight       0.989 0.999  
##  8 control       7 weight       0.957 0.708  
##  9 control       8 weight       0.906 0.188  
## 10 control       9 weight       0.815 0.0410 
## 11 dhq           0 weight       0.750 0.00206
## 12 dhq           1 weight       0.932 0.571  
## 13 dhq           2 weight       0.948 0.611  
## 14 dhq           3 weight       0.949 0.661  
## 15 dhq           4 weight       0.907 0.294  
## 16 dhq           5 weight       0.965 0.859  
## 17 dhq           6 weight       0.888 0.266  
## 18 dhq           7 weight       0.876 0.293  
## 19 dhq           8 weight       0.777 0.0516 
## 20 dhq           9 weight       0.947 0.557
```

```
tads10 %>% group_by(treatment, week) %>% shapiro_test(svl)
```

```
## # A tibble: 20 x 5
##    treatment  week variable statistic       p
##    <chr>     <int> <chr>        <dbl>   <dbl>
##  1 control       0 svl          0.943 0.194  
##  2 control       1 svl          0.861 0.0252 
##  3 control       2 svl          0.970 0.728  
##  4 control       3 svl          0.970 0.779  
##  5 control       4 svl          0.924 0.136  
##  6 control       5 svl          0.940 0.259  
##  7 control       6 svl          0.946 0.394  
##  8 control       7 svl          0.836 0.0187 
##  9 control       8 svl          0.898 0.151  
## 10 control       9 svl          0.784 0.0191 
## 11 dhq           0 svl          0.812 0.00955
## 12 dhq           1 svl          0.953 0.742  
## 13 dhq           2 svl          0.904 0.179  
## 14 dhq           3 svl          0.762 0.00503
## 15 dhq           4 svl          0.781 0.0124 
## 16 dhq           5 svl          0.979 0.957  
## 17 dhq           6 svl          0.870 0.187  
## 18 dhq           7 svl          0.751 0.0303 
## 19 dhq           8 svl          0.967 0.856  
## 20 dhq           9 svl          0.996 0.878
```

```
lmer1 <- lmer(svl ~ week * treatment + tank + (1 | individual), tads10)
lmer2 <- lmer(svl ~ week * treatment + (1 | individual), tads10)
anova(lmer1, lmer2)
```

```
## Data: tads10
## Models:
## lmer2: svl ~ week * treatment + (1 | individual)
## lmer1: svl ~ week * treatment + tank + (1 | individual)
##       npar    AIC    BIC  logLik deviance  Chisq Df Pr(>Chisq)
## lmer2    6 779.18 800.24 -383.59   767.18                     
## lmer1   14 788.97 838.10 -380.48   760.97 6.2169  8     0.6229
```

```
svl.lmer = lmer2
summary(svl.lmer)
```

```
## Linear mixed model fit by REML. t-tests use Satterthwaite's method [
## lmerModLmerTest]
## Formula: svl ~ week * treatment + (1 | individual)
##    Data: tads10
## 
## REML criterion at convergence: 780.3
## 
## Scaled residuals: 
##      Min       1Q   Median       3Q      Max 
## -2.67015 -0.61677 -0.00406  0.72722  2.39028 
## 
## Random effects:
##  Groups     Name        Variance Std.Dev.
##  individual (Intercept) 0.1983   0.4453  
##  Residual               1.1933   1.0924  
## Number of obs: 247, groups:  individual, 37
## 
## Fixed effects:
##                    Estimate Std. Error        df t value Pr(>|t|)    
## (Intercept)       4.600e+00  1.755e-01 9.571e+01  26.207   <2e-16 ***
## week              6.258e-01  3.266e-02 2.370e+02  19.162   <2e-16 ***
## treatmentdhq      2.091e-01  2.978e-01 9.889e+01   0.702    0.484    
## week:treatmentdhq 8.515e-03  5.933e-02 2.427e+02   0.144    0.886    
## ---
## Signif. codes:  0 '***' 0.001 '**' 0.01 '*' 0.05 '.' 0.1 ' ' 1
## 
## Correlation of Fixed Effects:
##             (Intr) week   trtmnt
## week        -0.686              
## treatmntdhq -0.589  0.405       
## wk:trtmntdh  0.378 -0.551 -0.677
```

##### Statistics for Width

```
desc = Summarize(width ~ treatment + week, data=tads05)
kbl(desc, caption = "Descriptive statistics for Width in 0.5% Experiment") %>% kable_paper("hover", full_width = F)
```

Descriptive statistics for Width in 0.5% Experiment

| treatment | week | n | nvalid | mean | sd | min | Q1 | median | Q3 | max |
| --- | --- | --- | --- | --- | --- | --- | --- | --- | --- | --- |
| control | 0 | 27 | 27 | 2.474074 | 0.2767845 | 1.8 | 2.300 | 2.50 | 2.700 | 3.0 |
| dhq | 0 | 27 | 27 | 2.466667 | 0.2218801 | 2.2 | 2.300 | 2.40 | 2.600 | 3.0 |
| control | 1 | 27 | 25 | 2.736000 | 0.2736786 | 2.2 | 2.600 | 2.70 | 2.900 | 3.2 |
| dhq | 1 | 27 | 26 | 2.880769 | 0.2884708 | 2.4 | 2.625 | 2.90 | 3.000 | 3.5 |
| control | 2 | 27 | 26 | 2.957692 | 0.3251745 | 2.3 | 2.700 | 3.00 | 3.200 | 3.4 |
| dhq | 2 | 27 | 24 | 2.887500 | 0.3893277 | 2.1 | 2.575 | 2.90 | 3.200 | 3.6 |
| control | 3 | 27 | 23 | 3.043478 | 0.4132203 | 2.1 | 2.750 | 3.10 | 3.300 | 3.9 |
| dhq | 3 | 27 | 22 | 3.059091 | 0.3246377 | 2.4 | 2.900 | 3.05 | 3.275 | 3.7 |

```
ggplot(tads05, aes(width, fill = treatment)) + 
  geom_density(alpha = 0.5) + facet_wrap(~week) + ggtitle('0.5% DHQ experiment') +
  scale_fill_manual(values = c("dhq" = "#dc4600", "control" = "#0064c8")) + 
  scale_color_manual(values = c("dhq" = "#dc4600", "control" = "#0064c8"))
```

```
tads05 %>% group_by(treatment, week) %>% shapiro_test(width)
```

```
## # A tibble: 8 x 5
##   treatment  week variable statistic       p
##   <chr>     <int> <chr>        <dbl>   <dbl>
## 1 control       0 width        0.975 0.728  
## 2 control       1 width        0.948 0.224  
## 3 control       2 width        0.942 0.153  
## 4 control       3 width        0.985 0.974  
## 5 dhq           0 width        0.886 0.00654
## 6 dhq           1 width        0.953 0.270  
## 7 dhq           2 width        0.979 0.871  
## 8 dhq           3 width        0.986 0.980
```

```
lmer1 <- lmer(width ~ week * treatment + tank + (1 | individual), tads05)
lmer2 <- lmer(width ~ week * treatment + (1 | individual), tads05)
anova(lmer1, lmer2)
```

```
## Data: tads05
## Models:
## lmer2: width ~ week * treatment + (1 | individual)
## lmer1: width ~ week * treatment + tank + (1 | individual)
##       npar    AIC    BIC  logLik deviance  Chisq Df Pr(>Chisq)   
## lmer2    6 114.63 134.42 -51.317  102.634                        
## lmer1   19 111.10 173.77 -36.551   73.103 29.532 13   0.005498 **
## ---
## Signif. codes:  0 '***' 0.001 '**' 0.01 '*' 0.05 '.' 0.1 ' ' 1
```

```
width.lmer = lmer1
summary(width.lmer)
```

```
## Linear mixed model fit by REML. t-tests use Satterthwaite's method [
## lmerModLmerTest]
## Formula: width ~ week * treatment + tank + (1 | individual)
##    Data: tads05
## 
## REML criterion at convergence: 128.3
## 
## Scaled residuals: 
##      Min       1Q   Median       3Q      Max 
## -2.99332 -0.57811  0.04156  0.69909  2.03897 
## 
## Random effects:
##  Groups     Name        Variance Std.Dev.
##  individual (Intercept) 0.00991  0.09955 
##  Residual               0.08447  0.29064 
## Number of obs: 200, groups:  individual, 54
## 
## Fixed effects:
##                     Estimate Std. Error         df t value Pr(>|t|)    
## (Intercept)         2.403498   0.132674  41.646264  18.116  < 2e-16 ***
## week                0.198886   0.026129 148.121923   7.612 2.92e-12 ***
## treatmentdhq        0.043832   0.072326 131.518217   0.606   0.5455    
## tankA07             0.355834   0.187382  41.831454   1.899   0.0645 .  
## tankA09            -0.144398   0.185896  43.231867  -0.777   0.4415    
## tankA14             0.263563   0.180607  38.170639   1.459   0.1527    
## tankA15             0.125000   0.176147  34.865329   0.710   0.4827    
## tankB07            -0.008278   0.139588  35.146778  -0.059   0.9530    
## tankB10             0.225000   0.176147  34.865329   1.277   0.2099    
## tankB15             0.001974   0.133973  35.488944   0.015   0.9883    
## tankB16             0.200000   0.176147  34.865329   1.135   0.2639    
## tankC07             0.356250   0.152548  34.865329   2.335   0.0254 *  
## tankC10             0.190440   0.144408  35.349714   1.319   0.1957    
## tankC11             0.010692   0.180694  38.228666   0.059   0.9531    
## tankC13             0.132863   0.155173  37.077804   0.856   0.3974    
## tankC15             0.162500   0.176147  34.865329   0.923   0.3626    
## week:treatmentdhq  -0.014990   0.037267 150.416102  -0.402   0.6881    
## ---
## Signif. codes:  0 '***' 0.001 '**' 0.01 '*' 0.05 '.' 0.1 ' ' 1
```

```
desc = Summarize(width ~ treatment + week, data=tads10)
kbl(desc, caption = "Descriptive statistics for Width in 1.0% Experiment") %>% kable_paper("hover", full_width = F)
```

Descriptive statistics for Width in 1.0% Experiment

| treatment | week | n | nvalid | mean | sd | min | Q1 | median | Q3 | max |
| --- | --- | --- | --- | --- | --- | --- | --- | --- | --- | --- |
| control | 0 | 24 | 24 | 2.370833 | 0.4206405 | 1.7 | 2.100 | 2.25 | 2.600 | 3.5 |
| dhq | 0 | 13 | 13 | 2.430769 | 0.4973004 | 1.9 | 2.100 | 2.20 | 2.700 | 3.7 |
| control | 1 | 24 | 15 | 2.526667 | 0.2491892 | 2.0 | 2.350 | 2.60 | 2.700 | 2.9 |
| dhq | 1 | 13 | 8 | 2.562500 | 0.4405759 | 2.0 | 2.275 | 2.50 | 2.750 | 3.2 |
| control | 2 | 24 | 21 | 2.985714 | 0.4150731 | 2.1 | 2.700 | 3.00 | 3.200 | 4.0 |
| dhq | 2 | 13 | 12 | 3.091667 | 0.6141636 | 2.3 | 2.700 | 2.95 | 3.275 | 4.3 |
| control | 3 | 24 | 19 | 3.289474 | 0.6090929 | 2.1 | 2.900 | 3.20 | 3.650 | 4.6 |
| dhq | 3 | 13 | 10 | 3.470000 | 0.5945119 | 2.9 | 3.025 | 3.25 | 3.950 | 4.6 |
| control | 4 | 24 | 19 | 3.889474 | 0.5404676 | 2.8 | 3.450 | 3.90 | 4.350 | 4.8 |
| dhq | 4 | 13 | 9 | 3.766667 | 0.6041523 | 3.0 | 3.400 | 3.60 | 4.300 | 4.8 |
| control | 5 | 24 | 19 | 4.194737 | 0.2504966 | 3.6 | 4.050 | 4.20 | 4.300 | 4.6 |
| dhq | 5 | 13 | 8 | 4.025000 | 0.3845220 | 3.6 | 3.775 | 3.95 | 4.125 | 4.7 |
| control | 6 | 24 | 17 | 4.241176 | 0.3890033 | 3.5 | 3.900 | 4.40 | 4.500 | 4.8 |
| dhq | 6 | 13 | 7 | 4.028571 | 0.6129554 | 2.8 | 3.900 | 4.20 | 4.350 | 4.7 |
| control | 7 | 24 | 13 | 4.230769 | 0.4714952 | 3.1 | 4.100 | 4.30 | 4.400 | 4.9 |
| dhq | 7 | 13 | 5 | 4.380000 | 0.1788854 | 4.1 | 4.400 | 4.40 | 4.400 | 4.6 |
| control | 8 | 24 | 12 | 3.975000 | 0.3621276 | 3.3 | 3.850 | 4.00 | 4.050 | 4.7 |
| dhq | 8 | 13 | 5 | 4.160000 | 0.3435113 | 3.7 | 4.000 | 4.10 | 4.500 | 4.5 |
| control | 9 | 24 | 8 | 3.862500 | 0.9379880 | 3.2 | 3.200 | 3.65 | 3.950 | 6.0 |
| dhq | 9 | 13 | 3 | 3.733333 | 0.1527525 | 3.6 | 3.650 | 3.70 | 3.800 | 3.9 |

```
ggplot(tads10, aes(width, fill = treatment)) + 
  geom_density(alpha = 0.5) + facet_wrap(~week) + ggtitle('1.0% DHQ experiment') +
  scale_fill_manual(values = c("dhq" = "#f0aa00", "control" = "#3caac8")) + 
  scale_color_manual(values = c("dhq" = "#f0aa00", "control" = "#3caac8"))
```

```
tads10 %>% group_by(treatment, week) %>% shapiro_test(weight)
```

```
## # A tibble: 20 x 5
##    treatment  week variable statistic       p
##    <chr>     <int> <chr>        <dbl>   <dbl>
##  1 control       0 weight       0.839 0.00172
##  2 control       1 weight       0.937 0.341  
##  3 control       2 weight       0.880 0.0148 
##  4 control       3 weight       0.963 0.605  
##  5 control       4 weight       0.982 0.959  
##  6 control       5 weight       0.936 0.221  
##  7 control       6 weight       0.989 0.999  
##  8 control       7 weight       0.957 0.708  
##  9 control       8 weight       0.906 0.188  
## 10 control       9 weight       0.815 0.0410 
## 11 dhq           0 weight       0.750 0.00206
## 12 dhq           1 weight       0.932 0.571  
## 13 dhq           2 weight       0.948 0.611  
## 14 dhq           3 weight       0.949 0.661  
## 15 dhq           4 weight       0.907 0.294  
## 16 dhq           5 weight       0.965 0.859  
## 17 dhq           6 weight       0.888 0.266  
## 18 dhq           7 weight       0.876 0.293  
## 19 dhq           8 weight       0.777 0.0516 
## 20 dhq           9 weight       0.947 0.557
```

```
tads10 %>% group_by(treatment, week) %>% shapiro_test(width)
```

```
## # A tibble: 20 x 5
##    treatment  week variable statistic       p
##    <chr>     <int> <chr>        <dbl>   <dbl>
##  1 control       0 width        0.924 0.0700 
##  2 control       1 width        0.940 0.379  
##  3 control       2 width        0.965 0.623  
##  4 control       3 width        0.984 0.981  
##  5 control       4 width        0.958 0.538  
##  6 control       5 width        0.950 0.401  
##  7 control       6 width        0.936 0.278  
##  8 control       7 width        0.915 0.218  
##  9 control       8 width        0.957 0.739  
## 10 control       9 width        0.736 0.00577
## 11 dhq           0 width        0.849 0.0279 
## 12 dhq           1 width        0.901 0.293  
## 13 dhq           2 width        0.910 0.214  
## 14 dhq           3 width        0.865 0.0879 
## 15 dhq           4 width        0.950 0.685  
## 16 dhq           5 width        0.889 0.230  
## 17 dhq           6 width        0.879 0.224  
## 18 dhq           7 width        0.863 0.238  
## 19 dhq           8 width        0.902 0.419  
## 20 dhq           9 width        0.964 0.637
```

```
lmer1 <- lmer(width ~ week * treatment + tank + (1 | individual), tads10)
lmer2 <- lmer(width ~ week * treatment + (1 | individual), tads10)
anova(lmer1, lmer2)
```

```
## Data: tads10
## Models:
## lmer2: width ~ week * treatment + (1 | individual)
## lmer1: width ~ week * treatment + tank + (1 | individual)
##       npar    AIC    BIC  logLik deviance  Chisq Df Pr(>Chisq)
## lmer2    6 420.94 441.99 -204.47   408.94                     
## lmer1   14 431.59 480.72 -201.79   403.59 5.3521  8     0.7194
```

```
width.lmer = lmer2
summary(width.lmer)
```

```
## Linear mixed model fit by REML. t-tests use Satterthwaite's method [
## lmerModLmerTest]
## Formula: width ~ week * treatment + (1 | individual)
##    Data: tads10
## 
## REML criterion at convergence: 428.5
## 
## Scaled residuals: 
##      Min       1Q   Median       3Q      Max 
## -2.55900 -0.65616  0.01001  0.66695  2.40147 
## 
## Random effects:
##  Groups     Name        Variance Std.Dev.
##  individual (Intercept) 0.02367  0.1538  
##  Residual               0.29267  0.5410  
## Number of obs: 247, groups:  individual, 37
## 
## Fixed effects:
##                    Estimate Std. Error        df t value Pr(>|t|)    
## (Intercept)       2.563e+00  8.032e-02 1.127e+02  31.918   <2e-16 ***
## week              2.316e-01  1.594e-02 2.405e+02  14.533   <2e-16 ***
## treatmentdhq      2.178e-02  1.365e-01 1.170e+02   0.160    0.874    
## week:treatmentdhq 2.736e-03  2.876e-02 2.425e+02   0.095    0.924    
## ---
## Signif. codes:  0 '***' 0.001 '**' 0.01 '*' 0.05 '.' 0.1 ' ' 1
## 
## Correlation of Fixed Effects:
##             (Intr) week   trtmnt
## week        -0.746              
## treatmntdhq -0.588  0.439       
## wk:trtmntdh  0.413 -0.554 -0.734
```

##### Statistic for Tail length

```
desc = Summarize(tail ~ treatment + week, data=tads05)
kbl(desc, caption = "Descriptive statistics for Tail length in 0.5% Experiment") %>% kable_paper("hover", full_width = F)
```

Descriptive statistics for Tail length in 0.5% Experiment

| treatment | week | n | nvalid | mean | sd | min | Q1 | median | Q3 | max |
| --- | --- | --- | --- | --- | --- | --- | --- | --- | --- | --- |
| control | 0 | 27 | 27 | 7.033333 | 0.5449065 | 6.1 | 6.650 | 7.00 | 7.400 | 8.6 |
| dhq | 0 | 27 | 27 | 7.225926 | 0.7788515 | 5.6 | 6.700 | 7.30 | 7.800 | 9.0 |
| control | 1 | 27 | 26 | 8.696154 | 0.8359334 | 6.9 | 8.125 | 8.60 | 9.300 | 10.3 |
| dhq | 1 | 27 | 26 | 8.811538 | 0.7279159 | 7.3 | 8.200 | 8.80 | 9.400 | 10.1 |
| control | 2 | 27 | 27 | 9.229630 | 0.8887393 | 7.9 | 8.700 | 9.10 | 9.650 | 11.2 |
| dhq | 2 | 27 | 24 | 9.141667 | 1.1201384 | 6.8 | 8.175 | 9.15 | 9.925 | 11.3 |
| control | 3 | 27 | 23 | 9.317391 | 1.0994429 | 7.4 | 8.400 | 9.60 | 10.150 | 11.0 |
| dhq | 3 | 27 | 22 | 9.368182 | 1.0426095 | 7.1 | 8.800 | 9.45 | 10.000 | 12.1 |

```
ggplot(tads05, aes(tail, fill = treatment)) + 
  geom_density(alpha = 0.5) + facet_wrap(~week) + ggtitle('0.5% DHQ experiment') +
  scale_fill_manual(values = c("dhq" = "#dc4600", "control" = "#0064c8")) + 
  scale_color_manual(values = c("dhq" = "#dc4600", "control" = "#0064c8"))
```

```
tads05 %>% group_by(treatment, week) %>% shapiro_test(tail)
```

```
## # A tibble: 8 x 5
##   treatment  week variable statistic      p
##   <chr>     <int> <chr>        <dbl>  <dbl>
## 1 control       0 tail         0.954 0.274 
## 2 control       1 tail         0.989 0.989 
## 3 control       2 tail         0.924 0.0505
## 4 control       3 tail         0.948 0.266 
## 5 dhq           0 tail         0.981 0.883 
## 6 dhq           1 tail         0.973 0.695 
## 7 dhq           2 tail         0.978 0.862 
## 8 dhq           3 tail         0.965 0.606
```

```
lmer1 <- lmer(tail ~ week * treatment + tank + (1 | individual), tads05)
lmer2 <- lmer(tail ~ week * treatment + (1 | individual), tads05)
anova(lmer1, lmer2)
```

```
## Data: tads05
## Models:
## lmer2: tail ~ week * treatment + (1 | individual)
## lmer1: tail ~ week * treatment + tank + (1 | individual)
##       npar    AIC    BIC  logLik deviance  Chisq Df Pr(>Chisq)   
## lmer2    6 547.74 567.59 -267.87   535.74                        
## lmer1   19 540.72 603.58 -251.36   502.72 33.021 13   0.001692 **
## ---
## Signif. codes:  0 '***' 0.001 '**' 0.01 '*' 0.05 '.' 0.1 ' ' 1
```

```
tail.lmer = lmer1
summary(tail.lmer)
```

```
## Linear mixed model fit by REML. t-tests use Satterthwaite's method [
## lmerModLmerTest]
## Formula: tail ~ week * treatment + tank + (1 | individual)
##    Data: tads05
## 
## REML criterion at convergence: 519.1
## 
## Scaled residuals: 
##      Min       1Q   Median       3Q      Max 
## -2.42293 -0.62559  0.04904  0.61152  1.97188 
## 
## Random effects:
##  Groups     Name        Variance Std.Dev.
##  individual (Intercept) 0.1550   0.3937  
##  Residual               0.6564   0.8102  
## Number of obs: 202, groups:  individual, 54
## 
## Fixed effects:
##                     Estimate Std. Error         df t value Pr(>|t|)    
## (Intercept)        7.140e+00  4.209e-01  4.273e+01  16.962   <2e-16 ***
## week               7.757e-01  7.277e-02  1.504e+02  10.660   <2e-16 ***
## treatmentdhq       1.613e-01  2.151e-01  1.156e+02   0.750   0.4550    
## tankA07           -3.089e-01  5.941e-01  4.371e+01  -0.520   0.6057    
## tankA09           -1.431e+00  5.887e-01  4.397e+01  -2.431   0.0192 *  
## tankA14            5.174e-13  5.649e-01  3.736e+01   0.000   1.0000    
## tankA15            1.000e-01  5.649e-01  3.736e+01   0.177   0.8604    
## tankB07            1.500e-01  4.466e-01  3.736e+01   0.336   0.7388    
## tankB10            6.500e-01  5.649e-01  3.736e+01   1.151   0.2572    
## tankB15            2.041e-01  4.292e-01  3.793e+01   0.476   0.6371    
## tankB16            4.625e-01  5.649e-01  3.736e+01   0.819   0.4181    
## tankC07            4.125e-01  4.892e-01  3.736e+01   0.843   0.4045    
## tankC10            9.276e-01  4.627e-01  3.778e+01   2.005   0.0522 .  
## tankC11            1.978e-02  5.762e-01  4.018e+01   0.034   0.9728    
## tankC13            6.867e-01  4.958e-01  3.922e+01   1.385   0.1738    
## tankC15            8.375e-01  5.649e-01  3.736e+01   1.483   0.1466    
## week:treatmentdhq -7.862e-02  1.040e-01  1.526e+02  -0.756   0.4510    
## ---
## Signif. codes:  0 '***' 0.001 '**' 0.01 '*' 0.05 '.' 0.1 ' ' 1
```

```
desc = Summarize(tail ~ treatment + week, data=tads10)
kbl(desc, caption = "Descriptive statistics for Tail length in 1.0% Experiment") %>% kable_paper("hover", full_width = F)
```

Descriptive statistics for Tail length in 1.0% Experiment

| treatment | week | n | nvalid | mean | sd | min | Q1 | median | Q3 | max |
| --- | --- | --- | --- | --- | --- | --- | --- | --- | --- | --- |
| control | 0 | 24 | 24 | 7.045833 | 1.3354625 | 5.4 | 6.225 | 6.75 | 7.700 | 10.3 |
| dhq | 0 | 13 | 13 | 7.676923 | 1.7823925 | 5.5 | 6.100 | 7.60 | 8.600 | 11.8 |
| control | 1 | 24 | 15 | 7.566667 | 1.1836183 | 5.8 | 6.500 | 7.90 | 8.400 | 9.5 |
| dhq | 1 | 13 | 8 | 8.600000 | 1.5399443 | 6.4 | 7.400 | 8.60 | 9.925 | 10.5 |
| control | 2 | 24 | 21 | 9.661905 | 1.6671761 | 6.0 | 8.700 | 9.70 | 10.900 | 12.6 |
| dhq | 2 | 13 | 12 | 9.958333 | 1.8637491 | 6.8 | 9.100 | 9.95 | 10.300 | 14.0 |
| control | 3 | 24 | 19 | 10.605263 | 1.9909444 | 6.7 | 9.300 | 10.30 | 12.000 | 14.1 |
| dhq | 3 | 13 | 10 | 11.650000 | 2.4627221 | 8.8 | 9.800 | 11.30 | 12.725 | 17.0 |
| control | 4 | 24 | 19 | 12.300000 | 1.8571184 | 9.2 | 10.500 | 13.00 | 13.700 | 14.9 |
| dhq | 4 | 13 | 9 | 12.111111 | 2.2734580 | 9.6 | 10.700 | 11.60 | 13.000 | 17.0 |
| control | 5 | 24 | 19 | 13.836842 | 0.8693615 | 12.4 | 13.350 | 13.60 | 14.450 | 16.0 |
| dhq | 5 | 13 | 8 | 13.825000 | 1.3123044 | 12.2 | 12.800 | 13.80 | 14.350 | 16.4 |
| control | 6 | 24 | 17 | 13.764706 | 1.9713388 | 10.1 | 13.600 | 13.90 | 15.200 | 16.7 |
| dhq | 6 | 13 | 7 | 13.885714 | 1.9827830 | 10.7 | 12.900 | 13.80 | 15.400 | 16.1 |
| control | 7 | 24 | 13 | 14.815385 | 1.6777198 | 10.5 | 14.300 | 15.00 | 15.100 | 17.1 |
| dhq | 7 | 13 | 5 | 15.760000 | 1.1282730 | 14.4 | 14.900 | 15.90 | 16.400 | 17.2 |
| control | 8 | 24 | 12 | 11.633333 | 3.8360333 | 1.8 | 10.150 | 11.65 | 14.475 | 16.1 |
| dhq | 8 | 13 | 5 | 14.480000 | 1.0281051 | 12.7 | 14.600 | 14.80 | 15.000 | 15.3 |
| control | 9 | 24 | 6 | 7.633333 | 3.9292069 | 2.2 | 5.450 | 7.15 | 10.725 | 12.5 |
| dhq | 9 | 13 | 3 | 10.933333 | 0.5131601 | 10.5 | 10.650 | 10.80 | 11.150 | 11.5 |

```
ggplot(tads10, aes(tail, fill = treatment)) + 
  geom_density(alpha = 0.5) + facet_wrap(~week) + ggtitle('1.0% DHQ experiment') +
  scale_fill_manual(values = c("dhq" = "#f0aa00", "control" = "#3caac8")) + 
  scale_color_manual(values = c("dhq" = "#f0aa00", "control" = "#3caac8"))
```

```
tads10 %>% group_by(treatment, week) %>% shapiro_test(weight)
```

```
## # A tibble: 20 x 5
##    treatment  week variable statistic       p
##    <chr>     <int> <chr>        <dbl>   <dbl>
##  1 control       0 weight       0.839 0.00172
##  2 control       1 weight       0.937 0.341  
##  3 control       2 weight       0.880 0.0148 
##  4 control       3 weight       0.963 0.605  
##  5 control       4 weight       0.982 0.959  
##  6 control       5 weight       0.936 0.221  
##  7 control       6 weight       0.989 0.999  
##  8 control       7 weight       0.957 0.708  
##  9 control       8 weight       0.906 0.188  
## 10 control       9 weight       0.815 0.0410 
## 11 dhq           0 weight       0.750 0.00206
## 12 dhq           1 weight       0.932 0.571  
## 13 dhq           2 weight       0.948 0.611  
## 14 dhq           3 weight       0.949 0.661  
## 15 dhq           4 weight       0.907 0.294  
## 16 dhq           5 weight       0.965 0.859  
## 17 dhq           6 weight       0.888 0.266  
## 18 dhq           7 weight       0.876 0.293  
## 19 dhq           8 weight       0.777 0.0516 
## 20 dhq           9 weight       0.947 0.557
```

```
tads10 %>% group_by(treatment, week) %>% shapiro_test(tail)
```

```
## # A tibble: 20 x 5
##    treatment  week variable statistic      p
##    <chr>     <int> <chr>        <dbl>  <dbl>
##  1 control       0 tail         0.905 0.0281
##  2 control       1 tail         0.922 0.206 
##  3 control       2 tail         0.977 0.882 
##  4 control       3 tail         0.967 0.715 
##  5 control       4 tail         0.889 0.0304
##  6 control       5 tail         0.953 0.444 
##  7 control       6 tail         0.909 0.0980
##  8 control       7 tail         0.870 0.0518
##  9 control       8 tail         0.869 0.0629
## 10 control       9 tail         0.954 0.776 
## 11 dhq           0 tail         0.933 0.368 
## 12 dhq           1 tail         0.889 0.231 
## 13 dhq           2 tail         0.945 0.561 
## 14 dhq           3 tail         0.905 0.251 
## 15 dhq           4 tail         0.898 0.243 
## 16 dhq           5 tail         0.926 0.483 
## 17 dhq           6 tail         0.937 0.614 
## 18 dhq           7 tail         0.971 0.879 
## 19 dhq           8 tail         0.787 0.0637
## 20 dhq           9 tail         0.949 0.567
```

```
tads10$t.tail = log10(tads10$tail)
tads10 %>% group_by(treatment, week) %>% shapiro_test(t.tail)
```

```
## # A tibble: 20 x 5
##    treatment  week variable statistic        p
##    <chr>     <int> <chr>        <dbl>    <dbl>
##  1 control       0 t.tail       0.945 0.207   
##  2 control       1 t.tail       0.904 0.110   
##  3 control       2 t.tail       0.952 0.378   
##  4 control       3 t.tail       0.967 0.712   
##  5 control       4 t.tail       0.868 0.0133  
##  6 control       5 t.tail       0.963 0.633   
##  7 control       6 t.tail       0.881 0.0333  
##  8 control       7 t.tail       0.822 0.0127  
##  9 control       8 t.tail       0.625 0.000176
## 10 control       9 t.tail       0.913 0.459   
## 11 dhq           0 t.tail       0.965 0.823   
## 12 dhq           1 t.tail       0.890 0.234   
## 13 dhq           2 t.tail       0.962 0.815   
## 14 dhq           3 t.tail       0.944 0.599   
## 15 dhq           4 t.tail       0.939 0.575   
## 16 dhq           5 t.tail       0.943 0.643   
## 17 dhq           6 t.tail       0.925 0.509   
## 18 dhq           7 t.tail       0.969 0.871   
## 19 dhq           8 t.tail       0.772 0.0466  
## 20 dhq           9 t.tail       0.954 0.587
```

```
ggplot(tads10, aes(t.tail, fill = treatment)) + 
  geom_density(alpha = 0.5) + facet_wrap(~week) + ggtitle('1.0% DHQ experiment') +
  scale_fill_manual(values = c("dhq" = "#f0aa00", "control" = "#3caac8")) + 
  scale_color_manual(values = c("dhq" = "#f0aa00", "control" = "#3caac8"))
```

```
tads10 %>% group_by(treatment, week) %>% shapiro_test(weight)
```

```
## # A tibble: 20 x 5
##    treatment  week variable statistic       p
##    <chr>     <int> <chr>        <dbl>   <dbl>
##  1 control       0 weight       0.839 0.00172
##  2 control       1 weight       0.937 0.341  
##  3 control       2 weight       0.880 0.0148 
##  4 control       3 weight       0.963 0.605  
##  5 control       4 weight       0.982 0.959  
##  6 control       5 weight       0.936 0.221  
##  7 control       6 weight       0.989 0.999  
##  8 control       7 weight       0.957 0.708  
##  9 control       8 weight       0.906 0.188  
## 10 control       9 weight       0.815 0.0410 
## 11 dhq           0 weight       0.750 0.00206
## 12 dhq           1 weight       0.932 0.571  
## 13 dhq           2 weight       0.948 0.611  
## 14 dhq           3 weight       0.949 0.661  
## 15 dhq           4 weight       0.907 0.294  
## 16 dhq           5 weight       0.965 0.859  
## 17 dhq           6 weight       0.888 0.266  
## 18 dhq           7 weight       0.876 0.293  
## 19 dhq           8 weight       0.777 0.0516 
## 20 dhq           9 weight       0.947 0.557
```

```
lmer1 <- lmer(t.tail ~ week * treatment + tank + (1 | individual), tads10)
lmer2 <- lmer(t.tail ~ week * treatment + (1 | individual), tads10)
anova(lmer1, lmer2)
```

```
## Data: tads10
## Models:
## lmer2: t.tail ~ week * treatment + (1 | individual)
## lmer1: t.tail ~ week * treatment + tank + (1 | individual)
##       npar     AIC     BIC logLik deviance  Chisq Df Pr(>Chisq)
## lmer2    6 -319.59 -298.58 165.79  -331.59                     
## lmer1   14 -307.93 -258.92 167.97  -335.93 4.3469  8     0.8245
```

```
tail.lmer = lmer2
summary(tail.lmer)
```

```
## Linear mixed model fit by REML. t-tests use Satterthwaite's method [
## lmerModLmerTest]
## Formula: t.tail ~ week * treatment + (1 | individual)
##    Data: tads10
## 
## REML criterion at convergence: -299.3
## 
## Scaled residuals: 
##     Min      1Q  Median      3Q     Max 
## -7.0189 -0.4751  0.1474  0.6471  1.7057 
## 
## Random effects:
##  Groups     Name        Variance Std.Dev.
##  individual (Intercept) 0.00000  0.000   
##  Residual               0.01538  0.124   
## Number of obs: 245, groups:  individual, 37
## 
## Fixed effects:
##                    Estimate Std. Error        df t value Pr(>|t|)    
## (Intercept)       9.102e-01  1.688e-02 2.410e+02  53.920  < 2e-16 ***
## week              2.694e-02  3.638e-03 2.410e+02   7.404 2.21e-12 ***
## treatmentdhq      6.105e-03  2.865e-02 2.410e+02   0.213    0.831    
## week:treatmentdhq 7.283e-03  6.395e-03 2.410e+02   1.139    0.256    
## ---
## Signif. codes:  0 '***' 0.001 '**' 0.01 '*' 0.05 '.' 0.1 ' ' 1
## 
## Correlation of Fixed Effects:
##             (Intr) week   trtmnt
## week        -0.820              
## treatmntdhq -0.589  0.483       
## wk:trtmntdh  0.467 -0.569 -0.807
## optimizer (nloptwrap) convergence code: 0 (OK)
## boundary (singular) fit: see ?isSingular
```

##### Statistics for Body area

```
desc = Summarize(area ~ treatment + week, data=tads05)
kbl(desc, caption = "Descriptive statistics for Body area in 0.5% Experiment") %>% kable_paper("hover", full_width = F)
```

Descriptive statistics for Body area in 0.5% Experiment

| treatment | week | n | nvalid | mean | sd | min | Q1 | median | Q3 | max |
| --- | --- | --- | --- | --- | --- | --- | --- | --- | --- | --- |
| control | 0 | 27 | 27 | 32.87778 | 5.788074 | 19.8 | 28.500 | 32.80 | 37.500 | 43.0 |
| dhq | 0 | 27 | 27 | 32.52593 | 5.702469 | 25.2 | 28.750 | 31.40 | 34.750 | 46.1 |
| control | 1 | 27 | 25 | 43.09600 | 7.301740 | 27.4 | 40.000 | 42.50 | 45.800 | 60.5 |
| dhq | 1 | 27 | 26 | 44.21923 | 7.473902 | 28.9 | 39.050 | 44.35 | 47.525 | 60.0 |
| control | 2 | 27 | 26 | 50.03846 | 9.745238 | 33.3 | 41.225 | 49.85 | 57.700 | 66.6 |
| dhq | 2 | 27 | 24 | 48.63333 | 10.293462 | 31.9 | 40.500 | 48.65 | 56.675 | 64.2 |
| control | 3 | 27 | 23 | 55.35217 | 12.761372 | 29.8 | 45.950 | 52.50 | 65.650 | 85.1 |
| dhq | 3 | 27 | 22 | 53.85909 | 9.778823 | 37.7 | 46.925 | 52.00 | 60.075 | 71.4 |

```
ggplot(tads05, aes(area, fill = treatment)) + 
  geom_density(alpha = 0.5) + facet_wrap(~week) + ggtitle('0.5% DHQ experiment') +
  scale_fill_manual(values = c("dhq" = "#dc4600", "control" = "#0064c8")) + 
  scale_color_manual(values = c("dhq" = "#dc4600", "control" = "#0064c8"))
```

```
tads05 %>% group_by(treatment, week) %>% shapiro_test(area)
```

```
## # A tibble: 8 x 5
##   treatment  week variable statistic      p
##   <chr>     <int> <chr>        <dbl>  <dbl>
## 1 control       0 area         0.976 0.753 
## 2 control       1 area         0.961 0.427 
## 3 control       2 area         0.959 0.370 
## 4 control       3 area         0.959 0.434 
## 5 dhq           0 area         0.909 0.0211
## 6 dhq           1 area         0.985 0.963 
## 7 dhq           2 area         0.932 0.109 
## 8 dhq           3 area         0.950 0.313
```

```
lmer1 <- lmer(area ~ week * treatment + tank + (1 | individual), tads05)
lmer2 <- lmer(area ~ week * treatment + (1 | individual), tads05)
anova(lmer1, lmer2)
```

```
## Data: tads05
## Models:
## lmer2: area ~ week * treatment + (1 | individual)
## lmer1: area ~ week * treatment + tank + (1 | individual)
##       npar    AIC    BIC  logLik deviance  Chisq Df Pr(>Chisq)    
## lmer2    6 1433.4 1453.2 -710.68   1421.4                         
## lmer1   19 1417.8 1480.4 -689.88   1379.8 41.595 13  7.625e-05 ***
## ---
## Signif. codes:  0 '***' 0.001 '**' 0.01 '*' 0.05 '.' 0.1 ' ' 1
```

```
area.lmer = lmer1
summary(area.lmer)
```

```
## Linear mixed model fit by REML. t-tests use Satterthwaite's method [
## lmerModLmerTest]
## Formula: area ~ week * treatment + tank + (1 | individual)
##    Data: tads05
## 
## REML criterion at convergence: 1323.9
## 
## Scaled residuals: 
##      Min       1Q   Median       3Q      Max 
## -2.85561 -0.59428 -0.07403  0.68863  2.77790 
## 
## Random effects:
##  Groups     Name        Variance Std.Dev.
##  individual (Intercept)  6.878   2.623   
##  Residual               58.036   7.618   
## Number of obs: 200, groups:  individual, 54
## 
## Fixed effects:
##                   Estimate Std. Error       df t value Pr(>|t|)    
## (Intercept)        28.1989     3.4827  41.8835   8.097 4.18e-10 ***
## week                7.6535     0.6849 148.3286  11.175  < 2e-16 ***
## treatmentdhq        0.4569     1.8971 131.5845   0.241   0.8101    
## tankA07            10.2915     4.9188  42.0812   2.092   0.0425 *  
## tankA09            -2.5220     4.8797  43.4700  -0.517   0.6079    
## tankA14            11.6105     4.7413  38.4055   2.449   0.0190 *  
## tankA15             7.2000     4.6246  35.0928   1.557   0.1285    
## tankB07             0.8451     3.6647  35.3750   0.231   0.8190    
## tankB10             9.3750     4.6246  35.0928   2.027   0.0503 .  
## tankB15             3.0512     3.5173  35.7186   0.867   0.3915    
## tankB16             8.4250     4.6246  35.0928   1.822   0.0770 .  
## tankC07            12.1250     4.0050  35.0928   3.027   0.0046 ** 
## tankC10            10.7105     3.7912  35.5785   2.825   0.0077 ** 
## tankC11             4.0500     4.7436  38.4637   0.854   0.3985    
## tankC13             6.5495     4.0737  37.3104   1.608   0.1163    
## tankC15            10.3875     4.6246  35.0928   2.246   0.0311 *  
## week:treatmentdhq  -0.6934     0.9768 150.6097  -0.710   0.4789    
## ---
## Signif. codes:  0 '***' 0.001 '**' 0.01 '*' 0.05 '.' 0.1 ' ' 1
```

```
desc = Summarize(area ~ treatment + week, data=tads10)
kbl(desc, caption = "Descriptive statistics for Body area in 1.0% Experiment") %>% kable_paper("hover", full_width = F)
```

Descriptive statistics for Body area in 1.0% Experiment

| treatment | week | n | nvalid | mean | sd | min | Q1 | median | Q3 | max |
| --- | --- | --- | --- | --- | --- | --- | --- | --- | --- | --- |
| control | 0 | 24 | 24 | 31.97500 | 11.068807 | 16.9 | 24.875 | 29.30 | 36.700 | 61.0 |
| dhq | 0 | 13 | 13 | 35.44615 | 15.985859 | 23.0 | 25.400 | 28.10 | 37.300 | 81.8 |
| control | 1 | 24 | 15 | 40.32667 | 10.710440 | 24.3 | 33.100 | 39.70 | 42.200 | 69.6 |
| dhq | 1 | 13 | 8 | 41.05000 | 14.442794 | 21.9 | 32.900 | 38.95 | 44.700 | 67.0 |
| control | 2 | 24 | 21 | 54.59048 | 17.663094 | 20.9 | 44.600 | 49.00 | 67.400 | 94.9 |
| dhq | 2 | 13 | 12 | 60.83333 | 23.819905 | 39.3 | 42.275 | 53.75 | 69.925 | 114.8 |
| control | 3 | 24 | 19 | 67.87895 | 22.040205 | 30.4 | 52.450 | 61.90 | 82.000 | 117.2 |
| dhq | 3 | 13 | 10 | 78.39000 | 29.428726 | 52.5 | 55.200 | 60.80 | 104.625 | 128.1 |
| control | 4 | 24 | 19 | 96.45789 | 23.158951 | 57.5 | 72.050 | 92.40 | 115.950 | 131.2 |
| dhq | 4 | 13 | 9 | 90.66667 | 30.272966 | 62.1 | 71.500 | 77.50 | 120.500 | 144.8 |
| control | 5 | 24 | 19 | 114.11579 | 12.414833 | 89.2 | 107.400 | 113.60 | 120.900 | 138.3 |
| dhq | 5 | 13 | 8 | 113.66250 | 19.035676 | 88.8 | 104.500 | 110.95 | 118.950 | 147.6 |
| control | 6 | 24 | 17 | 115.18824 | 20.734991 | 75.5 | 99.900 | 122.00 | 127.100 | 141.5 |
| dhq | 6 | 13 | 7 | 112.20000 | 26.275337 | 66.0 | 99.200 | 121.10 | 131.050 | 137.8 |
| control | 7 | 24 | 13 | 126.53846 | 21.906564 | 65.1 | 118.600 | 135.00 | 137.100 | 153.2 |
| dhq | 7 | 13 | 5 | 132.56000 | 9.262991 | 117.6 | 133.100 | 134.00 | 135.000 | 143.1 |
| control | 8 | 24 | 12 | 111.60000 | 21.207374 | 77.1 | 95.550 | 117.30 | 125.150 | 142.2 |
| dhq | 8 | 13 | 5 | 120.40000 | 25.911291 | 88.4 | 107.100 | 113.10 | 141.800 | 151.6 |
| control | 9 | 24 | 8 | 113.08750 | 62.521778 | 70.5 | 77.325 | 90.80 | 119.800 | 259.3 |
| dhq | 9 | 13 | 3 | 93.66667 | 4.368448 | 90.0 | 91.250 | 92.50 | 95.500 | 98.5 |

```
ggplot(tads10, aes(area, fill = treatment)) + 
  geom_density(alpha = 0.5) + facet_wrap(~week) + ggtitle('1.0% DHQ experiment') +
  scale_fill_manual(values = c("dhq" = "#f0aa00", "control" = "#3caac8")) + 
  scale_color_manual(values = c("dhq" = "#f0aa00", "control" = "#3caac8"))
```

```
tads10 %>% group_by(treatment, week) %>% shapiro_test(weight)
```

```
## # A tibble: 20 x 5
##    treatment  week variable statistic       p
##    <chr>     <int> <chr>        <dbl>   <dbl>
##  1 control       0 weight       0.839 0.00172
##  2 control       1 weight       0.937 0.341  
##  3 control       2 weight       0.880 0.0148 
##  4 control       3 weight       0.963 0.605  
##  5 control       4 weight       0.982 0.959  
##  6 control       5 weight       0.936 0.221  
##  7 control       6 weight       0.989 0.999  
##  8 control       7 weight       0.957 0.708  
##  9 control       8 weight       0.906 0.188  
## 10 control       9 weight       0.815 0.0410 
## 11 dhq           0 weight       0.750 0.00206
## 12 dhq           1 weight       0.932 0.571  
## 13 dhq           2 weight       0.948 0.611  
## 14 dhq           3 weight       0.949 0.661  
## 15 dhq           4 weight       0.907 0.294  
## 16 dhq           5 weight       0.965 0.859  
## 17 dhq           6 weight       0.888 0.266  
## 18 dhq           7 weight       0.876 0.293  
## 19 dhq           8 weight       0.777 0.0516 
## 20 dhq           9 weight       0.947 0.557
```

```
tads10 %>% group_by(treatment, week) %>% shapiro_test(area)
```

```
## # A tibble: 20 x 5
##    treatment  week variable statistic        p
##    <chr>     <int> <chr>        <dbl>    <dbl>
##  1 control       0 area         0.893 0.0155  
##  2 control       1 area         0.892 0.0725  
##  3 control       2 area         0.947 0.297   
##  4 control       3 area         0.964 0.654   
##  5 control       4 area         0.932 0.188   
##  6 control       5 area         0.983 0.972   
##  7 control       6 area         0.924 0.175   
##  8 control       7 area         0.786 0.00468 
##  9 control       8 area         0.944 0.555   
## 10 control       9 area         0.699 0.00220 
## 11 dhq           0 area         0.720 0.000878
## 12 dhq           1 area         0.931 0.527   
## 13 dhq           2 area         0.848 0.0347  
## 14 dhq           3 area         0.808 0.0179  
## 15 dhq           4 area         0.841 0.0600  
## 16 dhq           5 area         0.940 0.615   
## 17 dhq           6 area         0.893 0.291   
## 18 dhq           7 area         0.883 0.322   
## 19 dhq           8 area         0.945 0.702   
## 20 dhq           9 area         0.947 0.554
```

```
lmer1 <- lmer(area ~ week * treatment + tank + (1 | individual), tads10)
lmer2 <- lmer(area ~ week * treatment + (1 | individual), tads10)
anova(lmer1, lmer2)
```

```
## Data: tads10
## Models:
## lmer2: area ~ week * treatment + (1 | individual)
## lmer1: area ~ week * treatment + tank + (1 | individual)
##       npar    AIC    BIC  logLik deviance  Chisq Df Pr(>Chisq)
## lmer2    6 2293.4 2314.5 -1140.7   2281.4                     
## lmer1   14 2304.5 2353.6 -1138.2   2276.5 4.9241  8     0.7657
```

```
area.lmer = lmer2
summary(area.lmer)
```

```
## Linear mixed model fit by REML. t-tests use Satterthwaite's method [
## lmerModLmerTest]
## Formula: area ~ week * treatment + (1 | individual)
##    Data: tads10
## 
## REML criterion at convergence: 2270.6
## 
## Scaled residuals: 
##     Min      1Q  Median      3Q     Max 
## -2.7854 -0.6089 -0.1171  0.6928  4.3955 
## 
## Random effects:
##  Groups     Name        Variance Std.Dev.
##  individual (Intercept)  49.4     7.029  
##  Residual               571.8    23.912  
## Number of obs: 247, groups:  individual, 37
## 
## Fixed effects:
##                   Estimate Std. Error       df t value Pr(>|t|)    
## (Intercept)        36.2486     3.5699 123.1357  10.154   <2e-16 ***
## week               11.6724     0.7054 240.7085  16.548   <2e-16 ***
## treatmentdhq        2.9224     6.0666 127.3620   0.482    0.631    
## week:treatmentdhq  -0.1106     1.2734 242.6916  -0.087    0.931    
## ---
## Signif. codes:  0 '***' 0.001 '**' 0.01 '*' 0.05 '.' 0.1 ' ' 1
## 
## Correlation of Fixed Effects:
##             (Intr) week   trtmnt
## week        -0.742              
## treatmntdhq -0.588  0.436       
## wk:trtmntdh  0.411 -0.554 -0.730
```

### Movement in open field trials

The movement table looks like this:

```
movement = read.table('D:/STANFORD/DHQ_FEEDING/code/SM_tables/movement.csv', header=TRUE, sep=',')
kbl(head(movement), caption = "Head of movement table") %>% kable_paper("hover", full_width = F)
```

Head of movement table

| id | visible | center\_freq | exp\_rate | mob\_speed | mob\_rate | treatment | experiment | individual | week | tank |
| --- | --- | --- | --- | --- | --- | --- | --- | --- | --- | --- |
| CT01W0 | 1 | 0.1237 | 44 | 24.79 | 0.0857 | control | p05 | CT01 | 0 | B15 |
| CT01W1 | 1 | 0.0000 | 16 | 19.86 | 0.0141 | control | p05 | CT01 | 1 | B15 |
| CT01W2 | 1 | 0.0516 | 20 | 16.52 | 0.0664 | control | p05 | CT01 | 2 | B15 |
| CT01W3 | 1 | 1.0000 | 2 | 14.21 | 0.0286 | control | p05 | CT01 | 3 | B15 |
| CT02W0 | 1 | 0.6532 | 13 | 37.29 | 0.0027 | control | p05 | CT02 | 0 | C10 |
| CT02W1 | 1 | 0.2722 | 5 | 16.00 | 0.0207 | control | p05 | CT02 | 1 | C10 |

##### Plots

We keep only those samples for which the individuals were visible >95% of the time in the video, and separate the 0.5% DHQ and 1.0% DHQ experiments for plotting.

```
movement = subset(movement, visible >= 0.95)
movement05 = subset(movement, experiment == 'p05')
movement10 = subset(movement, experiment == 'p10')
```

```
movement05.gather = gather(movement05, type, measurement, center_freq, exp_rate, mob_speed, mob_rate)
ggplot(movement05.gather, aes(week, measurement, colour = treatment, group = individual)) +
  geom_smooth(method=loess, aes(group = treatment, fill=treatment), size = 2) +
  theme_gdocs() + ggtitle('0.5% DHQ experiment') +
  facet_wrap(~type, scales = "free_y") +
  scale_fill_manual(values = c("dhq" = "#dc4600", "control" = "#0064c8")) + 
  scale_color_manual(values = c("dhq" = "#dc4600", "control" = "#0064c8"))
```

```
movement10.gather = gather(movement10, type, measurement, center_freq, exp_rate, mob_speed, mob_rate)
ggplot(movement10.gather, aes(week, measurement, colour = treatment, group = individual)) +
  geom_smooth(method=loess, aes(group = treatment, fill=treatment), size = 2) +
  theme_gdocs() + ggtitle('1.0% DHQ experiment') + scale_x_discrete(limits = c(0,3,6,9)) +
  facet_wrap(~type, scales = "free_y") +
  scale_fill_manual(values = c("dhq" = "#f0aa00", "control" = "#3caac8")) + 
  scale_color_manual(values = c("dhq" = "#f0aa00", "control" = "#3caac8"))
```

```
ggplot(movement10.gather, aes(week, measurement, colour = treatment, group = individual)) +
  geom_smooth(method=loess, aes(group = treatment, fill=treatment), size = 2) +
  theme_gdocs() + ggtitle('1.0% DHQ experiment') + xlim(0,3) +
  facet_wrap(~type, scales = "free_y") +
  scale_fill_manual(values = c("dhq" = "#f0aa00", "control" = "#3caac8")) + 
  scale_color_manual(values = c("dhq" = "#f0aa00", "control" = "#3caac8"))
```

##### Statistics for Frequency in arena center

```
desc = Summarize(center_freq ~ treatment + week, data=movement05)
kbl(desc, caption = "Descriptive statistics for Frequency in arena center in 0.5% Experiment") %>% kable_paper("hover", full_width = F)
```

Descriptive statistics for Frequency in arena center in 0.5% Experiment

| treatment | week | n | mean | sd | min | Q1 | median | Q3 | max | percZero |
| --- | --- | --- | --- | --- | --- | --- | --- | --- | --- | --- |
| control | 0 | 27 | 0.3394481 | 0.2946449 | 0 | 0.089600 | 0.27630 | 0.531850 | 1.0000 | 3.703704 |
| dhq | 0 | 27 | 0.2552889 | 0.2388927 | 0 | 0.055550 | 0.21550 | 0.377000 | 0.9674 | 14.814815 |
| control | 1 | 26 | 0.1434192 | 0.1037004 | 0 | 0.064775 | 0.12800 | 0.233150 | 0.3317 | 7.692308 |
| dhq | 1 | 26 | 0.2151808 | 0.2692504 | 0 | 0.059250 | 0.09835 | 0.267600 | 1.0000 | 15.384615 |
| control | 2 | 27 | 0.1999444 | 0.1970231 | 0 | 0.048150 | 0.15280 | 0.320650 | 0.7468 | 7.407407 |
| dhq | 2 | 24 | 0.2726958 | 0.3014120 | 0 | 0.071150 | 0.17270 | 0.366250 | 1.0000 | 12.500000 |
| control | 3 | 24 | 0.2817542 | 0.3353498 | 0 | 0.027000 | 0.16670 | 0.324625 | 1.0000 | 12.500000 |
| dhq | 3 | 22 | 0.2829318 | 0.2951889 | 0 | 0.062350 | 0.15330 | 0.420550 | 1.0000 | 4.545454 |

```
ggplot(movement05, aes(center_freq, fill = treatment)) + 
  geom_density(alpha = 0.5) + facet_wrap(~week) + ggtitle('0.5% DHQ experiment') +
  scale_fill_manual(values = c("dhq" = "#dc4600", "control" = "#0064c8")) + 
  scale_color_manual(values = c("dhq" = "#dc4600", "control" = "#0064c8"))
```

```
movement05 %>% group_by(treatment, week) %>% shapiro_test(center_freq)
```

```
## # A tibble: 8 x 5
##   treatment  week variable    statistic         p
##   <chr>     <int> <chr>           <dbl>     <dbl>
## 1 control       0 center_freq     0.902 0.0151   
## 2 control       1 center_freq     0.938 0.119    
## 3 control       2 center_freq     0.852 0.00127  
## 4 control       3 center_freq     0.764 0.0000809
## 5 dhq           0 center_freq     0.893 0.00928  
## 6 dhq           1 center_freq     0.725 0.0000120
## 7 dhq           2 center_freq     0.800 0.000298 
## 8 dhq           3 center_freq     0.837 0.00202
```

```
lmer1 <- lmer(center_freq ~ week * treatment + tank + (1 | individual), movement05)
lmer2 <- lmer(center_freq ~ week * treatment + (1 | individual), movement05)
anova(lmer1, lmer2)
```

```
## Data: movement05
## Models:
## lmer2: center_freq ~ week * treatment + (1 | individual)
## lmer1: center_freq ~ week * treatment + tank + (1 | individual)
##       npar    AIC     BIC   logLik deviance  Chisq Df Pr(>Chisq)
## lmer2    6 39.936  59.815 -13.9681   27.936                     
## lmer1   19 48.187 111.138  -5.0935   10.187 17.749 13     0.1673
```

```
center.lmer = lmer2
summary(center.lmer)
```

```
## Linear mixed model fit by REML. t-tests use Satterthwaite's method [
## lmerModLmerTest]
## Formula: center_freq ~ week * treatment + (1 | individual)
##    Data: movement05
## 
## REML criterion at convergence: 49.8
## 
## Scaled residuals: 
##     Min      1Q  Median      3Q     Max 
## -1.3263 -0.6440 -0.3523  0.2962  3.0906 
## 
## Random effects:
##  Groups     Name        Variance Std.Dev.
##  individual (Intercept) 0.01010  0.1005  
##  Residual               0.06021  0.2454  
## Number of obs: 203, groups:  individual, 54
## 
## Fixed effects:
##                    Estimate Std. Error        df t value Pr(>|t|)    
## (Intercept)         0.26057    0.04429 154.43436   5.883 2.41e-08 ***
## week               -0.01347    0.02177 150.42004  -0.619    0.537    
## treatmentdhq       -0.02534    0.06272 154.78139  -0.404    0.687    
## week:treatmentdhq   0.02668    0.03123 153.40659   0.854    0.394    
## ---
## Signif. codes:  0 '***' 0.001 '**' 0.01 '*' 0.05 '.' 0.1 ' ' 1
## 
## Correlation of Fixed Effects:
##             (Intr) week   trtmnt
## week        -0.716              
## treatmntdhq -0.706  0.506       
## wk:trtmntdh  0.499 -0.697 -0.711
```

```
desc = Summarize(center_freq ~ treatment + week, data=movement10)
kbl(desc, caption = "Descriptive statistics for Frequency in arena center in 1.0% Experiment") %>% kable_paper("hover", full_width = F)
```

Descriptive statistics for Frequency in arena center in 1.0% Experiment

| treatment | week | n | mean | sd | min | Q1 | median | Q3 | max | percZero |
| --- | --- | --- | --- | --- | --- | --- | --- | --- | --- | --- |
| control | 0 | 24 | 0.2490792 | 0.2473198 | 0.000000 | 0.1073000 | 0.1613500 | 0.3357000 | 0.9468000 | 12.500000 |
| dhq | 0 | 13 | 0.3919308 | 0.4273527 | 0.000000 | 0.0934000 | 0.2143000 | 0.9931000 | 1.0000000 | 7.692308 |
| control | 1 | 14 | 0.2946020 | 0.3350959 | 0.000000 | 0.0725645 | 0.1344690 | 0.4074038 | 1.0000000 | 14.285714 |
| dhq | 1 | 7 | 0.5676813 | 0.4585398 | 0.000000 | 0.1455207 | 0.6835000 | 0.9996138 | 1.0000000 | 14.285714 |
| control | 2 | 18 | 0.4398831 | 0.3563915 | 0.000000 | 0.1517250 | 0.3396591 | 0.6622600 | 1.0000000 | 5.555556 |
| dhq | 2 | 11 | 0.2117851 | 0.1796266 | 0.000000 | 0.0775299 | 0.1697937 | 0.3376703 | 0.5221183 | 9.090909 |
| control | 3 | 4 | 0.2951000 | 0.1385699 | 0.147900 | 0.1923750 | 0.3014500 | 0.4041750 | 0.4296000 | 0.000000 |
| control | 4 | 16 | 0.3050131 | 0.3198458 | 0.000000 | 0.0499250 | 0.1320000 | 0.4973198 | 0.8326340 | 6.250000 |
| dhq | 4 | 8 | 0.3951054 | 0.3910997 | 0.000000 | 0.0403822 | 0.2949500 | 0.7478666 | 0.9995000 | 12.500000 |
| control | 5 | 15 | 0.2058067 | 0.2532781 | 0.049300 | 0.0731500 | 0.0989000 | 0.1953500 | 1.0000000 | 0.000000 |
| dhq | 5 | 7 | 0.4777467 | 0.4408865 | 0.008527 | 0.1289000 | 0.2653000 | 0.9063000 | 1.0000000 | 0.000000 |
| control | 6 | 11 | 0.5100727 | 0.2905275 | 0.025500 | 0.3313500 | 0.4189000 | 0.7043000 | 1.0000000 | 0.000000 |
| dhq | 6 | 4 | 0.1948250 | 0.3241201 | 0.000000 | 0.0020250 | 0.0517000 | 0.2445000 | 0.6759000 | 25.000000 |
| control | 7 | 17 | 0.4581294 | 0.3488698 | 0.013800 | 0.1468000 | 0.4460000 | 0.7354000 | 1.0000000 | 0.000000 |
| dhq | 7 | 7 | 0.5743286 | 0.3517231 | 0.106600 | 0.3096000 | 0.6582000 | 0.8181500 | 1.0000000 | 0.000000 |
| control | 8 | 16 | 0.5068524 | 0.3542205 | 0.016100 | 0.1626000 | 0.5355500 | 0.8018250 | 0.9973000 | 0.000000 |
| dhq | 8 | 6 | 0.4848667 | 0.3618649 | 0.069300 | 0.2865750 | 0.3522000 | 0.7411500 | 1.0000000 | 0.000000 |
| control | 9 | 13 | 0.6755683 | 0.3284740 | 0.076800 | 0.4648000 | 0.6941000 | 1.0000000 | 1.0000000 | 0.000000 |
| dhq | 9 | 4 | 0.4370500 | 0.4079058 | 0.000000 | 0.1454250 | 0.4385000 | 0.7301250 | 0.8712000 | 25.000000 |
| control | 10 | 11 | 0.6838182 | 0.3606141 | 0.031800 | 0.4292000 | 0.8671000 | 0.9623500 | 1.0000000 | 0.000000 |
| dhq | 10 | 2 | 0.3027000 | 0.2453661 | 0.129200 | 0.2159500 | 0.3027000 | 0.3894500 | 0.4762000 | 0.000000 |

```
ggplot(movement10, aes(center_freq, fill = treatment)) + 
  geom_density(alpha = 0.5) + facet_wrap(~week) + ggtitle('1.0% DHQ experiment')  +
  scale_fill_manual(values = c("dhq" = "#f0aa00", "control" = "#3caac8")) + 
  scale_color_manual(values = c("dhq" = "#f0aa00", "control" = "#3caac8"))
```

```
#movement10 %>% group_by(treatment, week) %>% shapiro_test(center_freq)
lmer1 <- lmer(center_freq ~ week * treatment + tank + (1 | individual), movement10)
lmer2 <- lmer(center_freq ~ week * treatment + (1 | individual), movement10)
anova(lmer1, lmer2)
```

```
## Data: movement10
## Models:
## lmer2: center_freq ~ week * treatment + (1 | individual)
## lmer1: center_freq ~ week * treatment + tank + (1 | individual)
##       npar    AIC    BIC  logLik deviance  Chisq Df Pr(>Chisq)
## lmer2    6 159.58 180.15 -73.789   147.58                     
## lmer1   14 171.71 219.72 -71.854   143.71 3.8697  8     0.8687
```

```
center.lmer = lmer2
summary(center.lmer)
```

```
## Linear mixed model fit by REML. t-tests use Satterthwaite's method [
## lmerModLmerTest]
## Formula: center_freq ~ week * treatment + (1 | individual)
##    Data: movement10
## 
## REML criterion at convergence: 172.2
## 
## Scaled residuals: 
##     Min      1Q  Median      3Q     Max 
## -1.7010 -0.8213 -0.2447  0.8544  2.1528 
## 
## Random effects:
##  Groups     Name        Variance Std.Dev.
##  individual (Intercept) 0.0000   0.0000  
##  Residual               0.1138   0.3374  
## Number of obs: 228, groups:  individual, 37
## 
## Fixed effects:
##                    Estimate Std. Error        df t value Pr(>|t|)    
## (Intercept)         0.23671    0.04654 224.00000   5.086 7.71e-07 ***
## week                0.03690    0.00817 224.00000   4.517 1.02e-05 ***
## treatmentdhq        0.13767    0.08050 224.00000   1.710   0.0886 .  
## week:treatmentdhq  -0.02908    0.01539 224.00000  -1.889   0.0601 .  
## ---
## Signif. codes:  0 '***' 0.001 '**' 0.01 '*' 0.05 '.' 0.1 ' ' 1
## 
## Correlation of Fixed Effects:
##             (Intr) week   trtmnt
## week        -0.818              
## treatmntdhq -0.578  0.473       
## wk:trtmntdh  0.434 -0.531 -0.794
## optimizer (nloptwrap) convergence code: 0 (OK)
## boundary (singular) fit: see ?isSingular
```

##### Statistics for Exploration rate

```
desc = Summarize(exp_rate ~ treatment + week, data=movement05)
kbl(desc, caption = "Descriptive statistics for Exploration rate in 0.5% Experiment") %>% kable_paper("hover", full_width = F)
```

Descriptive statistics for Exploration rate in 0.5% Experiment

| treatment | week | n | mean | sd | min | Q1 | median | Q3 | max |
| --- | --- | --- | --- | --- | --- | --- | --- | --- | --- |
| control | 0 | 27 | 30.74074 | 21.12161 | 1 | 15.5 | 27.0 | 42.50 | 90 |
| dhq | 0 | 27 | 32.96296 | 24.03760 | 1 | 15.0 | 27.0 | 45.50 | 83 |
| control | 1 | 26 | 40.65385 | 25.31868 | 1 | 25.0 | 35.0 | 54.00 | 98 |
| dhq | 1 | 26 | 27.61538 | 26.46519 | 1 | 7.0 | 25.0 | 38.25 | 85 |
| control | 2 | 27 | 35.33333 | 21.17691 | 1 | 19.5 | 34.0 | 46.50 | 81 |
| dhq | 2 | 24 | 31.79167 | 30.01591 | 1 | 8.5 | 21.0 | 46.00 | 100 |
| control | 3 | 24 | 28.29167 | 26.37766 | 1 | 7.0 | 18.5 | 41.50 | 88 |
| dhq | 3 | 22 | 32.63636 | 21.07706 | 1 | 13.5 | 32.5 | 47.75 | 77 |

```
ggplot(movement05, aes(exp_rate, fill = treatment)) + 
  geom_density(alpha = 0.5) + facet_wrap(~week) + ggtitle('0.5% DHQ experiment') +
  scale_fill_manual(values = c("dhq" = "#dc4600", "control" = "#0064c8")) + 
  scale_color_manual(values = c("dhq" = "#dc4600", "control" = "#0064c8"))
```

```
movement05 %>% group_by(treatment, week) %>% shapiro_test(exp_rate)
```

```
## # A tibble: 8 x 5
##   treatment  week variable statistic       p
##   <chr>     <int> <chr>        <dbl>   <dbl>
## 1 control       0 exp_rate     0.937 0.105  
## 2 control       1 exp_rate     0.940 0.133  
## 3 control       2 exp_rate     0.965 0.471  
## 4 control       3 exp_rate     0.862 0.00372
## 5 dhq           0 exp_rate     0.944 0.150  
## 6 dhq           1 exp_rate     0.858 0.00206
## 7 dhq           2 exp_rate     0.874 0.00633
## 8 dhq           3 exp_rate     0.959 0.462
```

```
lmer1 <- lmer(exp_rate ~ week * treatment + tank + (1 | individual), movement05)
lmer2 <- lmer(exp_rate ~ week * treatment + (1 | individual), movement05)
anova(lmer1, lmer2)
```

```
## Data: movement05
## Models:
## lmer2: exp_rate ~ week * treatment + (1 | individual)
## lmer1: exp_rate ~ week * treatment + tank + (1 | individual)
##       npar    AIC    BIC  logLik deviance  Chisq Df Pr(>Chisq)
## lmer2    6 1875.2 1895.0 -931.58   1863.2                     
## lmer1   19 1887.4 1950.4 -924.71   1849.4 13.741 13     0.3924
```

```
exp.lmer = lmer2
summary(exp.lmer)
```

```
## Linear mixed model fit by REML. t-tests use Satterthwaite's method [
## lmerModLmerTest]
## Formula: exp_rate ~ week * treatment + (1 | individual)
##    Data: movement05
## 
## REML criterion at convergence: 1848.7
## 
## Scaled residuals: 
##     Min      1Q  Median      3Q     Max 
## -1.5138 -0.7693 -0.1126  0.6170  2.3659 
## 
## Random effects:
##  Groups     Name        Variance Std.Dev.
##  individual (Intercept) 120.1    10.96   
##  Residual               486.0    22.05   
## Number of obs: 203, groups:  individual, 54
## 
## Fixed effects:
##                   Estimate Std. Error      df t value Pr(>|t|)    
## (Intercept)         35.479      4.155 143.838   8.538 1.77e-14 ***
## week                -1.187      1.957 151.520  -0.607    0.545    
## treatmentdhq        -4.578      5.884 144.179  -0.778    0.438    
## week:treatmentdhq    1.164      2.810 154.295   0.414    0.679    
## ---
## Signif. codes:  0 '***' 0.001 '**' 0.01 '*' 0.05 '.' 0.1 ' ' 1
## 
## Correlation of Fixed Effects:
##             (Intr) week   trtmnt
## week        -0.686              
## treatmntdhq -0.706  0.485       
## wk:trtmntdh  0.478 -0.696 -0.680
```

```
desc = Summarize(exp_rate ~ treatment + week, data=movement10)
kbl(desc, caption = "Descriptive statistics for Exploration rate in 1.0% Experiment") %>% kable_paper("hover", full_width = F)
```

Descriptive statistics for Exploration rate in 1.0% Experiment

| treatment | week | n | nvalid | mean | sd | min | Q1 | median | Q3 | max |
| --- | --- | --- | --- | --- | --- | --- | --- | --- | --- | --- |
| control | 0 | 24 | 23 | 40.13043 | 30.751651 | 2 | 15.00 | 38.0 | 67.50 | 95 |
| dhq | 0 | 13 | 13 | 37.46154 | 34.558201 | 1 | 4.00 | 27.0 | 67.00 | 96 |
| control | 1 | 14 | 14 | 27.07143 | 29.047611 | 1 | 5.75 | 19.5 | 38.00 | 97 |
| dhq | 1 | 7 | 5 | 6.60000 | 10.922454 | 1 | 1.00 | 1.0 | 4.00 | 26 |
| control | 2 | 18 | 18 | 36.72222 | 32.388824 | 1 | 4.50 | 30.0 | 65.50 | 91 |
| dhq | 2 | 11 | 11 | 52.72727 | 34.690318 | 2 | 26.00 | 47.0 | 86.00 | 99 |
| control | 3 | 4 | 3 | 57.00000 | 26.457513 | 27 | 47.00 | 67.0 | 72.00 | 77 |
| control | 4 | 16 | 16 | 34.25000 | 24.430855 | 1 | 20.50 | 25.0 | 50.00 | 91 |
| dhq | 4 | 8 | 8 | 33.50000 | 28.495614 | 1 | 7.75 | 34.0 | 48.50 | 80 |
| control | 5 | 15 | 15 | 47.20000 | 28.013262 | 1 | 31.00 | 43.0 | 64.00 | 97 |
| dhq | 5 | 7 | 7 | 27.57143 | 33.230795 | 2 | 7.50 | 20.0 | 28.50 | 99 |
| control | 6 | 11 | 11 | 35.81818 | 28.767406 | 2 | 16.00 | 26.0 | 53.50 | 91 |
| dhq | 6 | 4 | 4 | 25.50000 | 32.868425 | 1 | 10.00 | 13.5 | 29.00 | 74 |
| control | 7 | 17 | 17 | 25.94118 | 19.440520 | 1 | 11.00 | 24.0 | 35.00 | 63 |
| dhq | 7 | 7 | 7 | 25.71429 | 16.790020 | 14 | 14.00 | 18.0 | 34.00 | 52 |
| control | 8 | 16 | 16 | 22.68750 | 19.985724 | 2 | 10.50 | 17.0 | 27.25 | 85 |
| dhq | 8 | 6 | 6 | 17.16667 | 8.863784 | 7 | 10.75 | 17.0 | 21.00 | 31 |
| control | 9 | 13 | 13 | 22.00000 | 19.727308 | 2 | 3.00 | 18.0 | 37.00 | 55 |
| dhq | 9 | 4 | 4 | 13.00000 | 7.348469 | 2 | 12.50 | 16.5 | 17.00 | 17 |
| control | 10 | 11 | 11 | 18.81818 | 17.690778 | 2 | 4.00 | 16.0 | 30.50 | 51 |
| dhq | 10 | 2 | 2 | 18.00000 | 14.142136 | 8 | 13.00 | 18.0 | 23.00 | 28 |

```
ggplot(movement10, aes(exp_rate, fill = treatment)) + 
  geom_density(alpha = 0.5) + facet_wrap(~week) + ggtitle('1.0% DHQ experiment')  +
  scale_fill_manual(values = c("dhq" = "#f0aa00", "control" = "#3caac8")) + 
  scale_color_manual(values = c("dhq" = "#f0aa00", "control" = "#3caac8"))
```

```
#movement10 %>% group_by(treatment, week) %>% shapiro_test(exp_rate)
lmer1 <- lmer(exp_rate ~ week * treatment + tank + (1 | individual), movement10)
lmer2 <- lmer(exp_rate ~ week * treatment + (1 | individual), movement10)
anova(lmer1, lmer2)
```

```
## Data: movement10
## Models:
## lmer2: exp_rate ~ week * treatment + (1 | individual)
## lmer1: exp_rate ~ week * treatment + tank + (1 | individual)
##       npar    AIC    BIC  logLik deviance  Chisq Df Pr(>Chisq)
## lmer2    6 2112.3 2132.8 -1050.2   2100.3                     
## lmer1   14 2119.8 2167.5 -1045.9   2091.8 8.5837  8     0.3786
```

```
exp.lmer = lmer2
summary(exp.lmer)
```

```
## Linear mixed model fit by REML. t-tests use Satterthwaite's method [
## lmerModLmerTest]
## Formula: exp_rate ~ week * treatment + (1 | individual)
##    Data: movement10
## 
## REML criterion at convergence: 2088.7
## 
## Scaled residuals: 
##     Min      1Q  Median      3Q     Max 
## -1.9208 -0.7207 -0.1810  0.6528  2.5578 
## 
## Random effects:
##  Groups     Name        Variance Std.Dev.
##  individual (Intercept) 121.7    11.03   
##  Residual               622.6    24.95   
## Number of obs: 224, groups:  individual, 37
## 
## Fixed effects:
##                   Estimate Std. Error       df t value Pr(>|t|)    
## (Intercept)        40.9020     4.1878  82.5110   9.767 2.03e-15 ***
## week               -1.9455     0.6317 210.3601  -3.080  0.00235 ** 
## treatmentdhq        0.3379     7.2324  90.1519   0.047  0.96284    
## week:treatmentdhq  -0.9283     1.2219 218.9971  -0.760  0.44822    
## ---
## Signif. codes:  0 '***' 0.001 '**' 0.01 '*' 0.05 '.' 0.1 ' ' 1
## 
## Correlation of Fixed Effects:
##             (Intr) week   trtmnt
## week        -0.669              
## treatmntdhq -0.579  0.387       
## wk:trtmntdh  0.346 -0.517 -0.658
```

##### Statistics for Mobility rate

```
desc = Summarize(mob_rate ~ treatment + week, data=movement05)
kbl(desc, caption = "Descriptive statistics for Mobility rate in 0.5% Experiment") %>% kable_paper("hover", full_width = F)
```

Descriptive statistics for Mobility rate in 0.5% Experiment

| treatment | week | n | nvalid | mean | sd | min | Q1 | median | Q3 | max | percZero |
| --- | --- | --- | --- | --- | --- | --- | --- | --- | --- | --- | --- |
| control | 0 | 27 | 26 | 0.0372846 | 0.0294964 | 0.0010 | 0.010025 | 0.03180 | 0.058525 | 0.1006 | 0.000000 |
| dhq | 0 | 27 | 27 | 0.0304963 | 0.0296016 | 0.0006 | 0.014050 | 0.02470 | 0.035700 | 0.1300 | 0.000000 |
| control | 1 | 26 | 26 | 0.0526115 | 0.0441110 | 0.0021 | 0.019050 | 0.04165 | 0.079875 | 0.1453 | 0.000000 |
| dhq | 1 | 26 | 26 | 0.0365731 | 0.0257054 | 0.0006 | 0.014575 | 0.03890 | 0.048550 | 0.0872 | 0.000000 |
| control | 2 | 27 | 27 | 0.0470481 | 0.0440657 | 0.0000 | 0.017350 | 0.03460 | 0.066200 | 0.1909 | 3.703704 |
| dhq | 2 | 24 | 24 | 0.0411958 | 0.0362971 | 0.0009 | 0.017950 | 0.03290 | 0.052625 | 0.1561 | 0.000000 |
| control | 3 | 24 | 24 | 0.0382250 | 0.0425726 | 0.0000 | 0.011375 | 0.02840 | 0.037500 | 0.1646 | 4.166667 |
| dhq | 3 | 22 | 22 | 0.0340909 | 0.0181805 | 0.0039 | 0.023100 | 0.03495 | 0.040025 | 0.0825 | 0.000000 |

```
ggplot(movement05, aes(mob_rate, fill = treatment)) + 
  geom_density(alpha = 0.5) + facet_wrap(~week) + ggtitle('0.5% DHQ experiment') +
  scale_fill_manual(values = c("dhq" = "#dc4600", "control" = "#0064c8")) + 
  scale_color_manual(values = c("dhq" = "#dc4600", "control" = "#0064c8"))
```

```
movement05 %>% group_by(treatment, week) %>% shapiro_test(mob_rate)
```

```
## # A tibble: 8 x 5
##   treatment  week variable statistic         p
##   <chr>     <int> <chr>        <dbl>     <dbl>
## 1 control       0 mob_rate     0.931 0.0804   
## 2 control       1 mob_rate     0.876 0.00479  
## 3 control       2 mob_rate     0.834 0.000566 
## 4 control       3 mob_rate     0.767 0.0000907
## 5 dhq           0 mob_rate     0.769 0.0000423
## 6 dhq           1 mob_rate     0.948 0.205    
## 7 dhq           2 mob_rate     0.874 0.00621  
## 8 dhq           3 mob_rate     0.945 0.249
```

```
lmer1 <- lmer(mob_rate ~ week * treatment + tank + (1 | individual), movement05)
lmer2 <- lmer(mob_rate ~ week * treatment + (1 | individual), movement05)
anova(lmer1, lmer2)
```

```
## Data: movement05
## Models:
## lmer2: mob_rate ~ week * treatment + (1 | individual)
## lmer1: mob_rate ~ week * treatment + tank + (1 | individual)
##       npar     AIC     BIC logLik deviance  Chisq Df Pr(>Chisq)
## lmer2    6 -776.29 -756.44 394.14  -788.29                     
## lmer1   19 -767.69 -704.83 402.84  -805.69 17.402 13     0.1816
```

```
mob.lmer = lmer2
summary(mob.lmer)
```

```
## Linear mixed model fit by REML. t-tests use Satterthwaite's method [
## lmerModLmerTest]
## Formula: mob_rate ~ week * treatment + (1 | individual)
##    Data: movement05
## 
## REML criterion at convergence: -750.3
## 
## Scaled residuals: 
##     Min      1Q  Median      3Q     Max 
## -1.4959 -0.6363 -0.2558  0.3474  3.6168 
## 
## Random effects:
##  Groups     Name        Variance  Std.Dev.
##  individual (Intercept) 0.0001821 0.0135  
##  Residual               0.0010565 0.0325  
## Number of obs: 202, groups:  individual, 54
## 
## Fixed effects:
##                     Estimate Std. Error         df t value Pr(>|t|)    
## (Intercept)        4.391e-02  5.954e-03  1.547e+02   7.375 9.36e-12 ***
## week              -1.853e-04  2.911e-03  1.501e+02  -0.064    0.949    
## treatmentdhq      -1.066e-02  8.381e-03  1.540e+02  -1.272    0.205    
## week:treatmentdhq  1.930e-03  4.156e-03  1.525e+02   0.464    0.643    
## ---
## Signif. codes:  0 '***' 0.001 '**' 0.01 '*' 0.05 '.' 0.1 ' ' 1
## 
## Correlation of Fixed Effects:
##             (Intr) week   trtmnt
## week        -0.720              
## treatmntdhq -0.710  0.512       
## wk:trtmntdh  0.505 -0.700 -0.712
```

```
desc = Summarize(mob_rate ~ treatment + week, data=movement10)
kbl(desc, caption = "Descriptive statistics for Mobility rate in 1.0% Experiment") %>% kable_paper("hover", full_width = F)
```

Descriptive statistics for Mobility rate in 1.0% Experiment

| treatment | week | n | mean | sd | min | Q1 | median | Q3 | max |
| --- | --- | --- | --- | --- | --- | --- | --- | --- | --- |
| control | 0 | 24 | 0.1013417 | 0.0650294 | 0.0200000 | 0.0512500 | 0.0813500 | 0.1447500 | 0.2420000 |
| dhq | 0 | 13 | 0.1409623 | 0.0666904 | 0.0284000 | 0.1082000 | 0.1154000 | 0.1936000 | 0.2438000 |
| control | 1 | 14 | 0.0718758 | 0.0706979 | 0.0012387 | 0.0171108 | 0.0417178 | 0.1316641 | 0.2240000 |
| dhq | 1 | 7 | 0.0688081 | 0.0597481 | 0.0003000 | 0.0238329 | 0.0472000 | 0.1248456 | 0.1368000 |
| control | 2 | 18 | 0.1002232 | 0.0863817 | 0.0002000 | 0.0174250 | 0.0790108 | 0.1883000 | 0.2211462 |
| dhq | 2 | 11 | 0.1587031 | 0.0893078 | 0.0265000 | 0.0869682 | 0.1524175 | 0.2255500 | 0.3046000 |
| control | 3 | 4 | 0.1419250 | 0.0922574 | 0.0379000 | 0.0822250 | 0.1425500 | 0.2022500 | 0.2447000 |
| control | 4 | 16 | 0.0772343 | 0.0477542 | 0.0150000 | 0.0361250 | 0.0726537 | 0.1164250 | 0.1562975 |
| dhq | 4 | 8 | 0.0812837 | 0.0870760 | 0.0020000 | 0.0256000 | 0.0560000 | 0.0906450 | 0.2638760 |
| control | 5 | 15 | 0.1145733 | 0.0757989 | 0.0008000 | 0.0525500 | 0.1206000 | 0.1742500 | 0.2344000 |
| dhq | 5 | 7 | 0.1076717 | 0.0809804 | 0.0318000 | 0.0545500 | 0.0748000 | 0.1547008 | 0.2286000 |
| control | 6 | 11 | 0.0494545 | 0.0352923 | 0.0047000 | 0.0256500 | 0.0398000 | 0.0702500 | 0.1178000 |
| dhq | 6 | 4 | 0.1504750 | 0.0536107 | 0.0859000 | 0.1174750 | 0.1565500 | 0.1895500 | 0.2029000 |
| control | 7 | 17 | 0.1007118 | 0.0700623 | 0.0256000 | 0.0433000 | 0.0811000 | 0.1552000 | 0.2563000 |
| dhq | 7 | 7 | 0.1024143 | 0.0381616 | 0.0622000 | 0.0744000 | 0.0999000 | 0.1204500 | 0.1651000 |
| control | 8 | 16 | 0.0838091 | 0.0717465 | 0.0056000 | 0.0350750 | 0.0714500 | 0.1082250 | 0.2528000 |
| dhq | 8 | 6 | 0.0935000 | 0.0907491 | 0.0080000 | 0.0520750 | 0.0728500 | 0.0881500 | 0.2689000 |
| control | 9 | 13 | 0.0573834 | 0.0607627 | 0.0017000 | 0.0197000 | 0.0414000 | 0.0614000 | 0.2287000 |
| dhq | 9 | 4 | 0.1152000 | 0.0629271 | 0.0384000 | 0.0779250 | 0.1221500 | 0.1594250 | 0.1781000 |
| control | 10 | 11 | 0.0394818 | 0.0380481 | 0.0010000 | 0.0123500 | 0.0349000 | 0.0478000 | 0.1200000 |
| dhq | 10 | 2 | 0.0411000 | 0.0350725 | 0.0163000 | 0.0287000 | 0.0411000 | 0.0535000 | 0.0659000 |

```
ggplot(movement10, aes(mob_rate, fill = treatment)) + 
  geom_density(alpha = 0.5) + facet_wrap(~week) + ggtitle('1.0% DHQ experiment')  +
  scale_fill_manual(values = c("dhq" = "#f0aa00", "control" = "#3caac8")) + 
  scale_color_manual(values = c("dhq" = "#f0aa00", "control" = "#3caac8"))
```

```
#movement10 %>% group_by(treatment, week) %>% shapiro_test(mob_rate)
lmer1 <- lmer(mob_rate ~ week * treatment + tank + (1 | individual), movement10)
lmer2 <- lmer(mob_rate ~ week * treatment + (1 | individual), movement10)
anova(lmer1, lmer2)
```

```
## Data: movement10
## Models:
## lmer2: mob_rate ~ week * treatment + (1 | individual)
## lmer1: mob_rate ~ week * treatment + tank + (1 | individual)
##       npar     AIC     BIC logLik deviance  Chisq Df Pr(>Chisq)
## lmer2    6 -557.01 -536.43  284.5  -569.01                     
## lmer1   14 -549.79 -501.78  288.9  -577.79 8.7834  8     0.3609
```

```
mob.lmer = lmer2
summary(mob.lmer)
```

```
## Linear mixed model fit by REML. t-tests use Satterthwaite's method [
## lmerModLmerTest]
## Formula: mob_rate ~ week * treatment + (1 | individual)
##    Data: movement10
## 
## REML criterion at convergence: -532.3
## 
## Scaled residuals: 
##     Min      1Q  Median      3Q     Max 
## -1.7281 -0.7029 -0.2626  0.5755  2.6256 
## 
## Random effects:
##  Groups     Name        Variance  Std.Dev.
##  individual (Intercept) 0.0002751 0.01659 
##  Residual               0.0046797 0.06841 
## Number of obs: 228, groups:  individual, 37
## 
## Fixed effects:
##                     Estimate Std. Error         df t value Pr(>|t|)    
## (Intercept)        1.037e-01  1.008e-02  1.271e+02  10.294   <2e-16 ***
## week              -4.048e-03  1.684e-03  2.195e+02  -2.405    0.017 *  
## treatmentdhq       2.778e-02  1.736e-02  1.365e+02   1.601    0.112    
## week:treatmentdhq -6.334e-04  3.200e-03  2.240e+02  -0.198    0.843    
## ---
## Signif. codes:  0 '***' 0.001 '**' 0.01 '*' 0.05 '.' 0.1 ' ' 1
## 
## Correlation of Fixed Effects:
##             (Intr) week   trtmnt
## week        -0.761              
## treatmntdhq -0.581  0.442       
## wk:trtmntdh  0.400 -0.526 -0.740
```

##### Statistics for Mobility speed

```
desc = Summarize(mob_speed ~ treatment + week, data=movement05)
kbl(desc, caption = "Descriptive statistics for Mobility speed in 0.5% Experiment") %>% kable_paper("hover", full_width = F)
```

Descriptive statistics for Mobility speed in 0.5% Experiment

| treatment | week | n | nvalid | mean | sd | min | Q1 | median | Q3 | max | percZero |
| --- | --- | --- | --- | --- | --- | --- | --- | --- | --- | --- | --- |
| control | 0 | 27 | 26 | 23.81885 | 7.053486 | 13.95 | 19.225 | 22.320 | 26.2525 | 44.77 | 0.000000 |
| dhq | 0 | 27 | 27 | 25.92111 | 10.074434 | 10.98 | 19.410 | 24.630 | 28.6600 | 56.40 | 0.000000 |
| control | 1 | 26 | 26 | 28.28308 | 19.168028 | 12.94 | 19.385 | 23.885 | 29.2175 | 113.98 | 0.000000 |
| dhq | 1 | 26 | 26 | 20.22385 | 6.983047 | 11.55 | 14.845 | 18.965 | 23.7675 | 35.13 | 0.000000 |
| control | 2 | 27 | 27 | 23.70444 | 8.562183 | 0.00 | 20.185 | 23.390 | 26.6300 | 47.04 | 3.703704 |
| dhq | 2 | 24 | 24 | 22.90792 | 10.450445 | 13.59 | 17.435 | 20.555 | 24.7675 | 61.69 | 0.000000 |
| control | 3 | 24 | 24 | 19.94333 | 9.119554 | 0.00 | 14.525 | 18.450 | 22.8325 | 48.12 | 4.166667 |
| dhq | 3 | 22 | 22 | 20.76636 | 4.576549 | 12.98 | 18.015 | 21.450 | 22.8150 | 30.79 | 0.000000 |

```
ggplot(movement05, aes(mob_speed, fill = treatment)) + 
  geom_density(alpha = 0.5) + facet_wrap(~week) + ggtitle('0.5% DHQ experiment') +
  scale_fill_manual(values = c("dhq" = "#dc4600", "control" = "#0064c8")) + 
  scale_color_manual(values = c("dhq" = "#dc4600", "control" = "#0064c8"))
```

```
movement05 %>% group_by(treatment, week) %>% shapiro_test(mob_speed)
```

```
## # A tibble: 8 x 5
##   treatment  week variable  statistic            p
##   <chr>     <int> <chr>         <dbl>        <dbl>
## 1 control       0 mob_speed     0.888 0.00855     
## 2 control       1 mob_speed     0.555 0.0000000926
## 3 control       2 mob_speed     0.920 0.0402      
## 4 control       3 mob_speed     0.887 0.0116      
## 5 dhq           0 mob_speed     0.895 0.0103      
## 6 dhq           1 mob_speed     0.925 0.0583      
## 7 dhq           2 mob_speed     0.717 0.0000171   
## 8 dhq           3 mob_speed     0.952 0.338
```

```
lmer1 <- lmer(mob_speed ~ week * treatment + tank + (1 | individual), movement05)
lmer2 <- lmer(mob_speed ~ week * treatment + (1 | individual), movement05)
anova(lmer1, lmer2)
```

```
## Data: movement05
## Models:
## lmer2: mob_speed ~ week * treatment + (1 | individual)
## lmer1: mob_speed ~ week * treatment + tank + (1 | individual)
##       npar    AIC    BIC  logLik deviance  Chisq Df Pr(>Chisq)
## lmer2    6 1528.6 1548.5 -758.30   1516.6                     
## lmer1   19 1548.7 1611.5 -755.34   1510.7 5.9314 13     0.9486
```

```
speed.lmer = lmer2
summary(speed.lmer)
```

```
## Linear mixed model fit by REML. t-tests use Satterthwaite's method [
## lmerModLmerTest]
## Formula: mob_speed ~ week * treatment + (1 | individual)
##    Data: movement05
## 
## REML criterion at convergence: 1508.9
## 
## Scaled residuals: 
##     Min      1Q  Median      3Q     Max 
## -2.5308 -0.4735 -0.1137  0.2978  7.9084 
## 
## Random effects:
##  Groups     Name        Variance Std.Dev.
##  individual (Intercept) 16.08    4.010   
##  Residual               95.60    9.777   
## Number of obs: 202, groups:  individual, 54
## 
## Fixed effects:
##                     Estimate Std. Error         df t value Pr(>|t|)    
## (Intercept)        26.227962   1.786710 149.023390  14.679   <2e-16 ***
## week               -1.468584   0.875663 143.307340  -1.677   0.0957 .  
## treatmentdhq       -1.762134   2.514910 148.205163  -0.701   0.4846    
## week:treatmentdhq   0.008194   1.250143 145.885786   0.007   0.9948    
## ---
## Signif. codes:  0 '***' 0.001 '**' 0.01 '*' 0.05 '.' 0.1 ' ' 1
## 
## Correlation of Fixed Effects:
##             (Intr) week   trtmnt
## week        -0.722              
## treatmntdhq -0.710  0.513       
## wk:trtmntdh  0.506 -0.700 -0.713
```

```
desc = Summarize(mob_speed ~ treatment + week, data=movement10)
kbl(desc, caption = "Descriptive statistics for Mobility speed in 1.0% Experiment") %>% kable_paper("hover", full_width = F)
```

Descriptive statistics for Mobility speed in 1.0% Experiment

| treatment | week | n | nvalid | mean | sd | min | Q1 | median | Q3 | max |
| --- | --- | --- | --- | --- | --- | --- | --- | --- | --- | --- |
| control | 0 | 24 | 23 | 21.19696 | 9.237443 | 12.390000 | 15.30500 | 20.63000 | 23.14000 | 51.15 |
| dhq | 0 | 13 | 13 | 22.46342 | 11.710825 | 9.530000 | 13.63000 | 17.53000 | 30.95000 | 46.06 |
| control | 1 | 14 | 14 | 18.66695 | 11.270192 | 5.580868 | 11.15680 | 13.75107 | 27.88388 | 43.67 |
| dhq | 1 | 7 | 5 | 12.64333 | 2.140151 | 9.899660 | 10.79000 | 13.80000 | 14.13699 | 14.59 |
| control | 2 | 18 | 18 | 17.83793 | 9.034114 | 5.683640 | 9.88000 | 17.56236 | 26.96140 | 30.04 |
| dhq | 2 | 11 | 11 | 18.60934 | 4.389175 | 13.230000 | 15.43238 | 18.14000 | 22.11247 | 25.60 |
| control | 3 | 4 | 3 | 21.51667 | 7.726010 | 14.780000 | 17.30000 | 19.82000 | 24.88500 | 29.95 |
| control | 4 | 16 | 16 | 18.89507 | 5.188135 | 6.184644 | 16.06500 | 18.96125 | 21.96250 | 28.01 |
| dhq | 4 | 8 | 8 | 19.03420 | 7.210049 | 6.310000 | 14.90999 | 19.73500 | 25.10750 | 26.35 |
| control | 5 | 15 | 15 | 22.35133 | 7.008732 | 10.890000 | 18.23000 | 21.49000 | 26.20000 | 38.57 |
| dhq | 5 | 7 | 7 | 17.27526 | 7.417036 | 10.266835 | 13.80000 | 14.81000 | 17.69000 | 32.87 |
| control | 6 | 11 | 11 | 20.90818 | 7.207417 | 6.170000 | 18.40500 | 18.98000 | 25.32500 | 34.20 |
| dhq | 6 | 4 | 4 | 18.46750 | 4.658107 | 14.120000 | 15.16250 | 17.65500 | 20.96000 | 24.44 |
| control | 7 | 17 | 17 | 18.93471 | 6.351969 | 11.140000 | 15.36000 | 16.22000 | 19.34000 | 33.93 |
| dhq | 7 | 7 | 7 | 17.13143 | 3.534645 | 13.580000 | 14.64000 | 15.30000 | 19.57500 | 22.61 |
| control | 8 | 16 | 16 | 24.57506 | 8.894824 | 13.220000 | 17.68000 | 23.27000 | 31.75500 | 42.60 |
| dhq | 8 | 6 | 6 | 21.84333 | 7.284426 | 15.890000 | 16.99250 | 19.24000 | 23.78250 | 35.15 |
| control | 9 | 13 | 13 | 21.54015 | 6.378713 | 13.180000 | 16.02000 | 21.72000 | 25.86000 | 32.57 |
| dhq | 9 | 4 | 4 | 17.20750 | 3.421904 | 12.560000 | 15.68750 | 18.10500 | 19.62500 | 20.06 |
| control | 10 | 11 | 11 | 25.78909 | 6.650915 | 18.440000 | 20.58500 | 22.55000 | 31.05000 | 38.63 |
| dhq | 10 | 2 | 2 | 27.21500 | 15.224009 | 16.450000 | 21.83250 | 27.21500 | 32.59750 | 37.98 |

```
ggplot(movement10, aes(mob_speed, fill = treatment)) + 
  geom_density(alpha = 0.5) + facet_wrap(~week) + ggtitle('1.0% DHQ experiment')  +
  scale_fill_manual(values = c("dhq" = "#f0aa00", "control" = "#3caac8")) + 
  scale_color_manual(values = c("dhq" = "#f0aa00", "control" = "#3caac8"))
```

```
#movement10 %>% group_by(treatment, week) %>% shapiro_test(mob_speed)
lmer1 <- lmer(mob_speed ~ week * treatment + tank + (1 | individual), movement10)
lmer2 <- lmer(mob_speed ~ week * treatment + (1 | individual), movement10)
anova(lmer1, lmer2)
```

```
## Data: movement10
## Models:
## lmer2: mob_speed ~ week * treatment + (1 | individual)
## lmer1: mob_speed ~ week * treatment + tank + (1 | individual)
##       npar    AIC    BIC  logLik deviance  Chisq Df Pr(>Chisq)
## lmer2    6 1568.9 1589.4 -778.45   1556.9                     
## lmer1   14 1575.2 1623.0 -773.60   1547.2 9.6908  8     0.2874
```

```
speed.lmer = lmer2
summary(speed.lmer)
```

```
## Linear mixed model fit by REML. t-tests use Satterthwaite's method [
## lmerModLmerTest]
## Formula: mob_speed ~ week * treatment + (1 | individual)
##    Data: movement10
## 
## REML criterion at convergence: 1555.5
## 
## Scaled residuals: 
##     Min      1Q  Median      3Q     Max 
## -2.0391 -0.6490 -0.2185  0.5647  3.9229 
## 
## Random effects:
##  Groups     Name        Variance Std.Dev.
##  individual (Intercept)  4.732   2.175   
##  Residual               58.375   7.640   
## Number of obs: 224, groups:  individual, 37
## 
## Fixed effects:
##                   Estimate Std. Error       df t value Pr(>|t|)    
## (Intercept)        19.0468     1.1633 126.2992  16.373   <2e-16 ***
## week                0.3978     0.1903 215.4687   2.091   0.0377 *  
## treatmentdhq        0.3397     2.0202 136.5705   0.168   0.8667    
## week:treatmentdhq  -0.4790     0.3645 219.9971  -1.314   0.1901    
## ---
## Signif. codes:  0 '***' 0.001 '**' 0.01 '*' 0.05 '.' 0.1 ' ' 1
## 
## Correlation of Fixed Effects:
##             (Intr) week   trtmnt
## week        -0.746              
## treatmntdhq -0.576  0.429       
## wk:trtmntdh  0.389 -0.522 -0.730
```

### Feeding trials

#### Behaviors

We provide Table feeding05. Note that this table only contains data for the 0.5% DHQ experiment, as measuring behaviors for the 1.0% DHQ experiment was not possible due to the use of food in flakes.

```
feeding05 = read.csv('D:/STANFORD/DHQ_FEEDING/code/SM_tables/feeding05.csv', header=TRUE, sep=',')
kbl(head(feeding05), caption = "Head of feeding05 table") %>% kable_paper("hover", full_width = F)
```

Head of feeding05 table

| id | individual | treatment | week | tank | behavior | modifier | number | duration\_total | duration\_mean | duration\_stdev | inter\_event\_mean | inter\_event\_stdev | total\_percent | boris\_time |
| --- | --- | --- | --- | --- | --- | --- | --- | --- | --- | --- | --- | --- | --- | --- |
| CT04W0 | CT04 | control | 0 | B10 | breathing | bubble | 1 | 1.254 | 1.254 | NA | NA | NA | 0.1 | 30 |
| CT04W0 | CT04 | control | 0 | B10 | contact | NA | 0 | 0.000 | 0.000 | NA | NA | NA | 0.0 | 30 |
| CT04W0 | CT04 | control | 0 | B10 | eating | NA | 0 | 0.000 | 0.000 | NA | NA | NA | 0.0 | 30 |
| CT04W0 | CT04 | control | 0 | B10 | food | NA | 1 | NA | NA | NA | NA | NA | NA | 30 |
| CT04W0 | CT04 | control | 0 | B10 | moving | NA | 0 | 0.000 | 0.000 | NA | NA | NA | 0.0 | 30 |
| CT04W0 | CT04 | control | 0 | B10 | near | NA | 0 | 0.000 | 0.000 | NA | NA | NA | 0.0 | 30 |

```
info05 = subset(feeding05, select=c("id","individual","treatment","week","tank"))
info05 = unique(info05)
```

We usually evaluated 30 min of video but for some videos we evaluated less because there was an interruption (e.g., loud noise, hit the arena, sudden light change). Hence, we only use videos with 30 min ‘boris\_time’ for our behavior analysis.

```
feeding05 = subset(feeding05, boris_time == 30)
```

We subset four dataframes for number of events, percent of duration of events, and mean of duration of events.

##### Number of events: plots

Subset number of events, remove outliers, and obtain plots.

```
# subset dataframe
number = subset(feeding05, select = c('id', 'number', 'behavior'))
number = subset(number, behavior != 'breathing' & behavior != 'moving' & behavior != 'turning')
number = spread(number, behavior, number)
number = merge(info05, number, by = 'id')
breathing = subset(feeding05, behavior == 'breathing' & modifier != 'NA')
breathing.number = aggregate(breathing$number, by = list('id'=breathing$id), FUN = sum)
names(breathing.number)[names(breathing.number) == "x"] <- "breathing"
number = merge(number, breathing.number, by = 'id')

# remove outliers
number$eating = ifelse(number$eating > 20, NA, number$eating)
number$breathing = ifelse(number$breathing > 8, NA, number$breathing)

# plots
number.gather = gather(number, type, measurement, breathing, smelling,near,testing, eating)
ggplot(number.gather, aes(week, measurement, colour = treatment, group = individual)) +
  geom_smooth(method=loess, aes(group = treatment, fill=treatment), size = 2) +
  theme_gdocs() + ggtitle('0.5% DHQ experiment') +
  facet_wrap(~type, scales = "free_y") + ylab('Number of events') +
  scale_fill_manual(values = c("dhq" = "#dc4600", "control" = "#0064c8")) + 
  scale_color_manual(values = c("dhq" = "#dc4600", "control" = "#0064c8"))
```

##### Number of events: statistics per behavior

Breathing:

```
desc = Summarize(breathing ~ treatment + week, data=number)
kbl(desc, caption = "Descriptive statistics for Number of breaths in 0.5% Experiment") %>% kable_paper("hover", full_width = F)
```

Descriptive statistics for Number of breaths in 0.5% Experiment

| treatment | week | n | nvalid | mean | sd | min | Q1 | median | Q3 | max |
| --- | --- | --- | --- | --- | --- | --- | --- | --- | --- | --- |
| control | 0 | 20 | 19 | 2.105263 | 1.629408 | 1 | 1.00 | 2 | 2.50 | 8 |
| dhq | 0 | 19 | 19 | 3.473684 | 2.037657 | 1 | 2.00 | 3 | 5.00 | 8 |
| control | 1 | 19 | 18 | 3.722222 | 1.742397 | 1 | 2.25 | 4 | 5.50 | 6 |
| dhq | 1 | 17 | 15 | 2.200000 | 1.698739 | 1 | 1.00 | 2 | 2.50 | 6 |
| control | 2 | 16 | 14 | 2.642857 | 1.215739 | 1 | 2.00 | 2 | 3.75 | 5 |
| dhq | 2 | 15 | 13 | 2.615385 | 1.609268 | 1 | 1.00 | 2 | 4.00 | 5 |
| control | 3 | 14 | 13 | 2.692308 | 1.182132 | 1 | 2.00 | 2 | 3.00 | 5 |
| dhq | 3 | 13 | 13 | 1.846154 | 1.214232 | 1 | 1.00 | 1 | 2.00 | 5 |

```
ggplot(number, aes(breathing, fill = treatment)) + 
  geom_density(alpha = 0.5) + facet_wrap(~week) +
  scale_fill_manual(values = c("dhq" = "#dc4600", "control" = "#0064c8")) + 
  scale_color_manual(values = c("dhq" = "#dc4600", "control" = "#0064c8"))
```

```
number %>% group_by(treatment, week) %>% shapiro_test(breathing)
```

```
## # A tibble: 8 x 5
##   treatment  week variable  statistic         p
##   <chr>     <int> <chr>         <dbl>     <dbl>
## 1 control       0 breathing     0.643 0.0000127
## 2 control       1 breathing     0.893 0.0435   
## 3 control       2 breathing     0.894 0.0928   
## 4 control       3 breathing     0.824 0.0133   
## 5 dhq           0 breathing     0.913 0.0858   
## 6 dhq           1 breathing     0.714 0.000351 
## 7 dhq           2 breathing     0.835 0.0182   
## 8 dhq           3 breathing     0.743 0.00154
```

```
number$trans.breathing = log10(number$breathing)
number %>% group_by(treatment, week) %>% shapiro_test(trans.breathing)
```

```
## # A tibble: 8 x 5
##   treatment  week variable        statistic       p
##   <chr>     <int> <chr>               <dbl>   <dbl>
## 1 control       0 trans.breathing     0.823 0.00251
## 2 control       1 trans.breathing     0.871 0.0186 
## 3 control       2 trans.breathing     0.896 0.0979 
## 4 control       3 trans.breathing     0.879 0.0701 
## 5 dhq           0 trans.breathing     0.933 0.197  
## 6 dhq           1 trans.breathing     0.809 0.00479
## 7 dhq           2 trans.breathing     0.816 0.0105 
## 8 dhq           3 trans.breathing     0.787 0.00477
```

```
ggplot(number, aes(trans.breathing, fill = treatment)) + 
  geom_density(alpha = 0.5) + facet_wrap(~week) +
  scale_fill_manual(values = c("dhq" = "#dc4600", "control" = "#0064c8")) + 
  scale_color_manual(values = c("dhq" = "#dc4600", "control" = "#0064c8"))
```

```
lmer1 <- lmer(trans.breathing ~ week * treatment + tank + (1 | individual), number)
lmer2 <- lmer(trans.breathing ~ week * treatment + (1 | individual), number)
anova(lmer1, lmer2)
```

```
## Data: number
## Models:
## lmer2: trans.breathing ~ week * treatment + (1 | individual)
## lmer1: trans.breathing ~ week * treatment + tank + (1 | individual)
##       npar    AIC    BIC  logLik deviance  Chisq Df Pr(>Chisq)  
## lmer2    6 26.408 43.330 -7.2039  14.4079                       
## lmer1   18 27.680 78.446  4.1597  -8.3195 22.727 12    0.03013 *
## ---
## Signif. codes:  0 '***' 0.001 '**' 0.01 '*' 0.05 '.' 0.1 ' ' 1
```

```
breathing.num.lmer = lmer1
summary(breathing.num.lmer)
```

```
## Linear mixed model fit by REML. t-tests use Satterthwaite's method [
## lmerModLmerTest]
## Formula: trans.breathing ~ week * treatment + tank + (1 | individual)
##    Data: number
## 
## REML criterion at convergence: 46.4
## 
## Scaled residuals: 
##      Min       1Q   Median       3Q      Max 
## -1.87442 -0.68458 -0.06998  0.70125  2.12956 
## 
## Random effects:
##  Groups     Name        Variance Std.Dev.
##  individual (Intercept) 0.003559 0.05965 
##  Residual               0.060166 0.24529 
## Number of obs: 124, groups:  individual, 47
## 
## Fixed effects:
##                   Estimate Std. Error       df t value Pr(>|t|)  
## (Intercept)        0.16830    0.11074 25.69964   1.520   0.1408  
## week               0.04020    0.02826 93.32231   1.422   0.1583  
## treatmentdhq       0.09051    0.07143 90.09308   1.267   0.2083  
## tankA07            0.10944    0.15894 22.74770   0.689   0.4981  
## tankA09           -0.04057    0.20617 71.54205  -0.197   0.8446  
## tankA15           -0.02418    0.14056 18.26794  -0.172   0.8653  
## tankB07            0.22211    0.12064 22.89471   1.841   0.0786 .
## tankB10            0.18598    0.14056 18.26794   1.323   0.2021  
## tankB15            0.23211    0.11512 23.29320   2.016   0.0555 .
## tankB16            0.31802    0.15599 27.34757   2.039   0.0513 .
## tankC07            0.02406    0.13189 24.79636   0.182   0.8567  
## tankC10            0.02915    0.12788 21.72547   0.228   0.8218  
## tankC11            0.28700    0.15597 27.17927   1.840   0.0767 .
## tankC13            0.25747    0.12682 20.83927   2.030   0.0553 .
## tankC15            0.09301    0.14470 20.38520   0.643   0.5275  
## week:treatmentdhq -0.10287    0.04010 91.49998  -2.565   0.0119 *
## ---
## Signif. codes:  0 '***' 0.001 '**' 0.01 '*' 0.05 '.' 0.1 ' ' 1
```

Smelling:

```
desc = Summarize(smelling ~ treatment + week, data=number)
kbl(desc, caption = "Descriptive statistics for Number of smelling events in 0.5% Experiment") %>% kable_paper("hover", full_width = F)
```

Descriptive statistics for Number of smelling events in 0.5% Experiment

| treatment | week | n | mean | sd | min | Q1 | median | Q3 | max | percZero |
| --- | --- | --- | --- | --- | --- | --- | --- | --- | --- | --- |
| control | 0 | 20 | 12.100000 | 13.96952 | 0 | 3.0 | 7.5 | 14.00 | 55 | 10.000000 |
| dhq | 0 | 19 | 9.421053 | 11.96437 | 0 | 2.0 | 4.0 | 8.00 | 41 | 5.263158 |
| control | 1 | 19 | 16.894737 | 12.74055 | 0 | 8.5 | 15.0 | 20.50 | 50 | 5.263158 |
| dhq | 1 | 17 | 12.705882 | 10.65812 | 1 | 4.0 | 10.0 | 16.00 | 34 | 0.000000 |
| control | 2 | 16 | 13.375000 | 15.15201 | 0 | 3.0 | 6.5 | 15.75 | 51 | 6.250000 |
| dhq | 2 | 15 | 12.133333 | 10.35696 | 0 | 4.0 | 9.0 | 18.50 | 36 | 13.333333 |
| control | 3 | 14 | 16.928571 | 14.20977 | 0 | 3.5 | 14.5 | 26.75 | 38 | 14.285714 |
| dhq | 3 | 13 | 10.615385 | 11.31031 | 0 | 2.0 | 9.0 | 13.00 | 36 | 15.384615 |

```
ggplot(number, aes(smelling, fill = treatment)) + 
  geom_density(alpha = 0.5) + facet_wrap(~week) +
  scale_fill_manual(values = c("dhq" = "#dc4600", "control" = "#0064c8")) + 
  scale_color_manual(values = c("dhq" = "#dc4600", "control" = "#0064c8"))
```

```
number %>% group_by(treatment, week) %>% shapiro_test(smelling)
```

```
## # A tibble: 8 x 5
##   treatment  week variable statistic         p
##   <chr>     <int> <chr>        <dbl>     <dbl>
## 1 control       0 smelling     0.768 0.000297 
## 2 control       1 smelling     0.901 0.0513   
## 3 control       2 smelling     0.775 0.00128  
## 4 control       3 smelling     0.897 0.102    
## 5 dhq           0 smelling     0.717 0.0000888
## 6 dhq           1 smelling     0.886 0.0406   
## 7 dhq           2 smelling     0.923 0.217    
## 8 dhq           3 smelling     0.859 0.0370
```

```
number$trans.smelling = sqrt(number$smelling)
number %>% group_by(treatment, week) %>% shapiro_test(trans.smelling)
```

```
## # A tibble: 8 x 5
##   treatment  week variable       statistic      p
##   <chr>     <int> <chr>              <dbl>  <dbl>
## 1 control       0 trans.smelling     0.951 0.383 
## 2 control       1 trans.smelling     0.979 0.933 
## 3 control       2 trans.smelling     0.926 0.214 
## 4 control       3 trans.smelling     0.916 0.195 
## 5 dhq           0 trans.smelling     0.890 0.0322
## 6 dhq           1 trans.smelling     0.949 0.435 
## 7 dhq           2 trans.smelling     0.963 0.739 
## 8 dhq           3 trans.smelling     0.965 0.830
```

```
ggplot(number, aes(trans.breathing, fill = treatment)) + 
  geom_density(alpha = 0.5) + facet_wrap(~week) +
  scale_fill_manual(values = c("dhq" = "#dc4600", "control" = "#0064c8")) + 
  scale_color_manual(values = c("dhq" = "#dc4600", "control" = "#0064c8"))
```

```
lmer1 <- lmer(trans.smelling ~ week * treatment + tank + (1 | individual), number)
lmer2 <- lmer(trans.smelling ~ week * treatment + (1 | individual), number)
anova(lmer1, lmer2)
```

```
## Data: number
## Models:
## lmer2: trans.smelling ~ week * treatment + (1 | individual)
## lmer1: trans.smelling ~ week * treatment + tank + (1 | individual)
##       npar    AIC    BIC  logLik deviance  Chisq Df Pr(>Chisq)
## lmer2    6 535.49 552.83 -261.74   523.49                     
## lmer1   18 545.70 597.73 -254.85   509.70 13.785 12     0.3147
```

```
smelling.num.lmer = lmer2
summary(smelling.num.lmer)
```

```
## Linear mixed model fit by REML. t-tests use Satterthwaite's method [
## lmerModLmerTest]
## Formula: trans.smelling ~ week * treatment + (1 | individual)
##    Data: number
## 
## REML criterion at convergence: 528.3
## 
## Scaled residuals: 
##      Min       1Q   Median       3Q      Max 
## -2.09014 -0.63152 -0.03565  0.69163  2.03998 
## 
## Random effects:
##  Groups     Name        Variance Std.Dev.
##  individual (Intercept) 0.7227   0.8501  
##  Residual               2.5209   1.5877  
## Number of obs: 133, groups:  individual, 47
## 
## Fixed effects:
##                    Estimate Std. Error        df t value Pr(>|t|)    
## (Intercept)         3.16327    0.35390 102.09181   8.938  1.8e-14 ***
## week                0.12481    0.17706  98.94962   0.705    0.483    
## treatmentdhq       -0.40828    0.50628 105.02110  -0.806    0.422    
## week:treatmentdhq  -0.06328    0.25484  99.88743  -0.248    0.804    
## ---
## Signif. codes:  0 '***' 0.001 '**' 0.01 '*' 0.05 '.' 0.1 ' ' 1
## 
## Correlation of Fixed Effects:
##             (Intr) week   trtmnt
## week        -0.665              
## treatmntdhq -0.699  0.465       
## wk:trtmntdh  0.462 -0.695 -0.663
```

Near the food:

```
desc = Summarize(near ~ treatment + week, data=number)
kbl(desc, caption = "Descriptive statistics for Number of times approaching the food in 0.5% Experiment") %>% kable_paper("hover", full_width = F)
```

Descriptive statistics for Number of times approaching the food in 0.5% Experiment

| treatment | week | n | mean | sd | min | Q1 | median | Q3 | max | percZero |
| --- | --- | --- | --- | --- | --- | --- | --- | --- | --- | --- |
| control | 0 | 20 | 5.200000 | 7.331260 | 0 | 1.0 | 3.0 | 5.00 | 30 | 20.00000 |
| dhq | 0 | 19 | 5.473684 | 7.640306 | 0 | 0.0 | 2.0 | 6.00 | 27 | 36.84211 |
| control | 1 | 19 | 5.263158 | 5.970689 | 0 | 0.5 | 3.0 | 8.00 | 19 | 26.31579 |
| dhq | 1 | 17 | 3.882353 | 5.914837 | 0 | 0.0 | 2.0 | 5.00 | 24 | 41.17647 |
| control | 2 | 16 | 13.187500 | 19.204926 | 0 | 0.0 | 6.0 | 14.25 | 70 | 31.25000 |
| dhq | 2 | 15 | 4.400000 | 4.468940 | 0 | 1.0 | 3.0 | 7.00 | 13 | 20.00000 |
| control | 3 | 14 | 9.000000 | 9.663572 | 0 | 3.0 | 7.5 | 10.75 | 33 | 21.42857 |
| dhq | 3 | 13 | 2.769231 | 3.515533 | 0 | 0.0 | 1.0 | 6.00 | 10 | 46.15385 |

```
ggplot(number, aes(near, fill = treatment)) + 
  geom_density(alpha = 0.5) + facet_wrap(~week) +
  scale_fill_manual(values = c("dhq" = "#dc4600", "control" = "#0064c8")) + 
  scale_color_manual(values = c("dhq" = "#dc4600", "control" = "#0064c8"))
```

```
number %>% group_by(treatment, week) %>% shapiro_test(near)
```

```
## # A tibble: 8 x 5
##   treatment  week variable statistic         p
##   <chr>     <int> <chr>        <dbl>     <dbl>
## 1 control       0 near         0.670 0.0000173
## 2 control       1 near         0.833 0.00364  
## 3 control       2 near         0.721 0.000289 
## 4 control       3 near         0.809 0.00642  
## 5 dhq           0 near         0.712 0.0000775
## 6 dhq           1 near         0.664 0.0000466
## 7 dhq           2 near         0.858 0.0230   
## 8 dhq           3 near         0.792 0.00545
```

```
number$trans.near = sqrt(number$near)
number %>% group_by(treatment, week) %>% shapiro_test(trans.near)
```

```
## # A tibble: 8 x 5
##   treatment  week variable   statistic      p
##   <chr>     <int> <chr>          <dbl>  <dbl>
## 1 control       0 trans.near     0.906 0.0546
## 2 control       1 trans.near     0.923 0.126 
## 3 control       2 trans.near     0.902 0.0870
## 4 control       3 trans.near     0.927 0.278 
## 5 dhq           0 trans.near     0.864 0.0114
## 6 dhq           1 trans.near     0.848 0.0101
## 7 dhq           2 trans.near     0.938 0.362 
## 8 dhq           3 trans.near     0.818 0.0111
```

```
ggplot(number, aes(trans.breathing, fill = treatment)) + 
  geom_density(alpha = 0.5) + facet_wrap(~week) +
  scale_fill_manual(values = c("dhq" = "#dc4600", "control" = "#0064c8")) + 
  scale_color_manual(values = c("dhq" = "#dc4600", "control" = "#0064c8"))
```

```
lmer1 <- lmer(trans.near ~ week * treatment + tank + (1 | individual), number)
lmer2 <- lmer(trans.near ~ week * treatment + (1 | individual), number)
anova(lmer1, lmer2)
```

```
## Data: number
## Models:
## lmer2: trans.near ~ week * treatment + (1 | individual)
## lmer1: trans.near ~ week * treatment + tank + (1 | individual)
##       npar    AIC    BIC  logLik deviance  Chisq Df Pr(>Chisq)
## lmer2    6 511.87 529.21 -249.94   499.87                     
## lmer1   18 521.71 573.73 -242.85   485.71 14.167 12     0.2902
```

```
near.num.lmer = lmer2
summary(near.num.lmer)
```

```
## Linear mixed model fit by REML. t-tests use Satterthwaite's method [
## lmerModLmerTest]
## Formula: trans.near ~ week * treatment + (1 | individual)
##    Data: number
## 
## REML criterion at convergence: 505.7
## 
## Scaled residuals: 
##     Min      1Q  Median      3Q     Max 
## -1.7150 -0.8179 -0.0165  0.5633  3.3934 
## 
## Random effects:
##  Groups     Name        Variance Std.Dev.
##  individual (Intercept) 0.3162   0.5623  
##  Residual               2.3110   1.5202  
## Number of obs: 133, groups:  individual, 47
## 
## Fixed effects:
##                    Estimate Std. Error        df t value Pr(>|t|)    
## (Intercept)         1.74199    0.31429 112.86437   5.543 1.98e-07 ***
## week                0.29861    0.16818 102.90859   1.775   0.0788 .  
## treatmentdhq       -0.07587    0.45066 115.18862  -0.168   0.8666    
## week:treatmentdhq  -0.42124    0.24190 103.60484  -1.741   0.0846 .  
## ---
## Signif. codes:  0 '***' 0.001 '**' 0.01 '*' 0.05 '.' 0.1 ' ' 1
## 
## Correlation of Fixed Effects:
##             (Intr) week   trtmnt
## week        -0.715              
## treatmntdhq -0.697  0.499       
## wk:trtmntdh  0.497 -0.695 -0.713
```

Testing:

```
desc = Summarize(testing ~ treatment + week, data=number)
kbl(desc, caption = "Descriptive statistics for Number of testing events in 0.5% Experiment") %>% kable_paper("hover", full_width = F)
```

Descriptive statistics for Number of testing events in 0.5% Experiment

| treatment | week | n | mean | sd | min | Q1 | median | Q3 | max | percZero |
| --- | --- | --- | --- | --- | --- | --- | --- | --- | --- | --- |
| control | 0 | 20 | 1.0000000 | 2.339591 | 0 | 0 | 0.0 | 0.25 | 8 | 75.00000 |
| dhq | 0 | 19 | 1.6315789 | 3.148378 | 0 | 0 | 0.0 | 1.50 | 11 | 57.89474 |
| control | 1 | 19 | 1.5263158 | 1.806421 | 0 | 0 | 1.0 | 3.00 | 5 | 47.36842 |
| dhq | 1 | 17 | 1.5882353 | 2.450990 | 0 | 0 | 0.0 | 3.00 | 7 | 64.70588 |
| control | 2 | 16 | 3.5000000 | 4.163332 | 0 | 0 | 1.5 | 5.50 | 13 | 31.25000 |
| dhq | 2 | 15 | 1.1333333 | 1.884776 | 0 | 0 | 0.0 | 1.50 | 7 | 53.33333 |
| control | 3 | 14 | 3.2857143 | 4.268463 | 0 | 0 | 2.0 | 5.50 | 14 | 42.85714 |
| dhq | 3 | 13 | 0.8461538 | 1.214232 | 0 | 0 | 0.0 | 1.00 | 4 | 53.84615 |

```
ggplot(number, aes(testing, fill = treatment)) + 
  geom_density(alpha = 0.5) + facet_wrap(~week) +
  scale_fill_manual(values = c("dhq" = "#dc4600", "control" = "#0064c8")) + 
  scale_color_manual(values = c("dhq" = "#dc4600", "control" = "#0064c8"))
```

```
number %>% group_by(treatment, week) %>% shapiro_test(testing)
```

```
## # A tibble: 8 x 5
##   treatment  week variable statistic           p
##   <chr>     <int> <chr>        <dbl>       <dbl>
## 1 control       0 testing      0.496 0.000000293
## 2 control       1 testing      0.805 0.00134    
## 3 control       2 testing      0.826 0.00610    
## 4 control       3 testing      0.792 0.00397    
## 5 dhq           0 testing      0.588 0.00000339 
## 6 dhq           1 testing      0.697 0.000107   
## 7 dhq           2 testing      0.662 0.000101   
## 8 dhq           3 testing      0.743 0.00154
```

```
number$trans.testing = sqrt(number$testing)
number %>% group_by(treatment, week) %>% shapiro_test(trans.testing)
```

```
## # A tibble: 8 x 5
##   treatment  week variable      statistic          p
##   <chr>     <int> <chr>             <dbl>      <dbl>
## 1 control       0 trans.testing     0.575 0.00000163
## 2 control       1 trans.testing     0.797 0.00103   
## 3 control       2 trans.testing     0.903 0.0891    
## 4 control       3 trans.testing     0.852 0.0239    
## 5 dhq           0 trans.testing     0.736 0.000154  
## 6 dhq           1 trans.testing     0.687 0.0000835 
## 7 dhq           2 trans.testing     0.791 0.00284   
## 8 dhq           3 trans.testing     0.782 0.00425
```

```
ggplot(number, aes(trans.breathing, fill = treatment)) + 
  geom_density(alpha = 0.5) + facet_wrap(~week) +
  scale_fill_manual(values = c("dhq" = "#dc4600", "control" = "#0064c8")) + 
  scale_color_manual(values = c("dhq" = "#dc4600", "control" = "#0064c8"))
```

```
lmer1 <- lmer(trans.testing ~ week * treatment + tank + (1 | individual), number)
lmer2 <- lmer(trans.testing ~ week * treatment + (1 | individual), number)
anova(lmer1, lmer2)
```

```
## Data: number
## Models:
## lmer2: trans.testing ~ week * treatment + (1 | individual)
## lmer1: trans.testing ~ week * treatment + tank + (1 | individual)
##       npar    AIC    BIC  logLik deviance  Chisq Df Pr(>Chisq)
## lmer2    6 388.08 405.42 -188.04   376.08                     
## lmer1   18 404.21 456.24 -184.11   368.21 7.8663 12     0.7955
```

```
testing.num.lmer = lmer2
summary(testing.num.lmer)
```

```
## Linear mixed model fit by REML. t-tests use Satterthwaite's method [
## lmerModLmerTest]
## Formula: trans.testing ~ week * treatment + (1 | individual)
##    Data: number
## 
## REML criterion at convergence: 385.6
## 
## Scaled residuals: 
##     Min      1Q  Median      3Q     Max 
## -1.4522 -0.7127 -0.3743  0.6966  2.4208 
## 
## Random effects:
##  Groups     Name        Variance Std.Dev.
##  individual (Intercept) 0.1089   0.3299  
##  Residual               0.9235   0.9610  
## Number of obs: 133, groups:  individual, 47
## 
## Fixed effects:
##                   Estimate Std. Error       df t value Pr(>|t|)   
## (Intercept)         0.5290     0.1966 112.6696   2.691  0.00821 **
## week                0.3329     0.1062 100.9497   3.136  0.00225 **
## treatmentdhq        0.2160     0.2820 115.0998   0.766  0.44520   
## week:treatmentdhq  -0.3787     0.1527 101.6492  -2.480  0.01479 * 
## ---
## Signif. codes:  0 '***' 0.001 '**' 0.01 '*' 0.05 '.' 0.1 ' ' 1
## 
## Correlation of Fixed Effects:
##             (Intr) week   trtmnt
## week        -0.722              
## treatmntdhq -0.697  0.504       
## wk:trtmntdh  0.502 -0.695 -0.721
```

Eating:

```
desc = Summarize(eating ~ treatment + week, data=number)
kbl(desc, caption = "Descriptive statistics for Number of eating events in 0.5% Experiment") %>% kable_paper("hover", full_width = F)
```

Descriptive statistics for Number of eating events in 0.5% Experiment

| treatment | week | n | nvalid | mean | sd | min | Q1 | median | Q3 | max | percZero |
| --- | --- | --- | --- | --- | --- | --- | --- | --- | --- | --- | --- |
| control | 0 | 20 | 20 | 1.1500000 | 2.058998 | 0 | 0 | 0 | 2.0 | 7 | 65.00000 |
| dhq | 0 | 19 | 19 | 2.5789474 | 4.259836 | 0 | 0 | 0 | 3.0 | 15 | 52.63158 |
| control | 1 | 19 | 19 | 2.9473684 | 4.648461 | 0 | 0 | 1 | 3.5 | 18 | 42.10526 |
| dhq | 1 | 17 | 17 | 1.5882353 | 2.346775 | 0 | 0 | 0 | 3.0 | 6 | 64.70588 |
| control | 2 | 16 | 15 | 4.4666667 | 5.356794 | 0 | 0 | 3 | 8.0 | 15 | 40.00000 |
| dhq | 2 | 15 | 15 | 1.3333333 | 1.397276 | 0 | 0 | 1 | 2.0 | 5 | 33.33333 |
| control | 3 | 14 | 13 | 4.0769231 | 5.219539 | 0 | 0 | 1 | 9.0 | 14 | 46.15385 |
| dhq | 3 | 13 | 13 | 0.8461538 | 1.405119 | 0 | 0 | 0 | 1.0 | 5 | 53.84615 |

```
ggplot(number, aes(eating, fill = treatment)) + 
  geom_density(alpha = 0.5) + facet_wrap(~week) +
  scale_fill_manual(values = c("dhq" = "#dc4600", "control" = "#0064c8")) + 
  scale_color_manual(values = c("dhq" = "#dc4600", "control" = "#0064c8"))
```

```
number %>% group_by(treatment, week) %>% shapiro_test(eating)
```

```
## # A tibble: 8 x 5
##   treatment  week variable statistic          p
##   <chr>     <int> <chr>        <dbl>      <dbl>
## 1 control       0 eating       0.632 0.00000640
## 2 control       1 eating       0.683 0.0000349 
## 3 control       2 eating       0.806 0.00445   
## 4 control       3 eating       0.780 0.00397   
## 5 dhq           0 eating       0.675 0.0000287 
## 6 dhq           1 eating       0.690 0.0000899 
## 7 dhq           2 eating       0.843 0.0139    
## 8 dhq           3 eating       0.649 0.000178
```

```
number$trans.eating = sqrt(number$eating)
number %>% group_by(treatment, week) %>% shapiro_test(trans.eating)
```

```
## # A tibble: 8 x 5
##   treatment  week variable     statistic         p
##   <chr>     <int> <chr>            <dbl>     <dbl>
## 1 control       0 trans.eating     0.696 0.0000347
## 2 control       1 trans.eating     0.855 0.00808  
## 3 control       2 trans.eating     0.849 0.0170   
## 4 control       3 trans.eating     0.805 0.00787  
## 5 dhq           0 trans.eating     0.792 0.000876 
## 6 dhq           1 trans.eating     0.670 0.0000538
## 7 dhq           2 trans.eating     0.867 0.0302   
## 8 dhq           3 trans.eating     0.775 0.00351
```

```
ggplot(number, aes(trans.breathing, fill = treatment)) + 
  geom_density(alpha = 0.5) + facet_wrap(~week) +
  scale_fill_manual(values = c("dhq" = "#dc4600", "control" = "#0064c8")) + 
  scale_color_manual(values = c("dhq" = "#dc4600", "control" = "#0064c8"))
```

```
lmer1 <- lmer(trans.eating ~ week * treatment + tank + (1 | individual), number)
lmer2 <- lmer(trans.eating ~ week * treatment + (1 | individual), number)
anova(lmer1, lmer2)
```

```
## Data: number
## Models:
## lmer2: trans.eating ~ week * treatment + (1 | individual)
## lmer1: trans.eating ~ week * treatment + tank + (1 | individual)
##       npar   AIC    BIC  logLik deviance Chisq Df Pr(>Chisq)
## lmer2    6 413.5 430.76 -200.75    401.5                    
## lmer1   18 428.2 479.95 -196.10    392.2 9.304 12     0.6768
```

```
eating.num.lmer = lmer2
summary(eating.num.lmer)
```

```
## Linear mixed model fit by REML. t-tests use Satterthwaite's method [
## lmerModLmerTest]
## Formula: trans.eating ~ week * treatment + (1 | individual)
##    Data: number
## 
## REML criterion at convergence: 409.9
## 
## Scaled residuals: 
##     Min      1Q  Median      3Q     Max 
## -1.4230 -0.7480 -0.2932  0.7053  2.5410 
## 
## Random effects:
##  Groups     Name        Variance Std.Dev.
##  individual (Intercept) 0.212    0.4604  
##  Residual               1.115    1.0561  
## Number of obs: 131, groups:  individual, 47
## 
## Fixed effects:
##                   Estimate Std. Error       df t value Pr(>|t|)   
## (Intercept)         0.7220     0.2251 106.2587   3.208  0.00177 **
## week                0.3219     0.1199  99.8681   2.685  0.00850 **
## treatmentdhq        0.2399     0.3221 108.8691   0.745  0.45806   
## week:treatmentdhq  -0.4336     0.1705  99.6304  -2.543  0.01253 * 
## ---
## Signif. codes:  0 '***' 0.001 '**' 0.01 '*' 0.05 '.' 0.1 ' ' 1
## 
## Correlation of Fixed Effects:
##             (Intr) week   trtmnt
## week        -0.690              
## treatmntdhq -0.699  0.482       
## wk:trtmntdh  0.485 -0.703 -0.691
```

##### Percent of duration of events: plots and descriptive statistics

Subset percent duration of events, and obtain plots.

```
# subset dataframe
percent = subset(feeding05, select = c('id', 'total_percent', 'behavior'))
percent = subset(percent, behavior != 'breathing' & behavior != 'moving' & behavior != 'turning')
percent = spread(percent, behavior, total_percent)
percent = merge(info05, percent, by = 'id')
breathing = subset(feeding05, behavior == 'breathing' & modifier != 'NA')
breathing.percent = aggregate(breathing$total_percent, by = list('id'=breathing$id), FUN = sum)
names(breathing.percent)[names(breathing.percent) == "x"] <- "breathing"
percent = merge(percent, breathing.percent, by = 'id')

# plots
percent.gather = gather(percent, type, measurement, breathing, smelling,near,testing, eating)
ggplot(percent.gather, aes(week, measurement, colour = treatment, group = individual)) +
  geom_smooth(method=loess, aes(group = treatment, fill=treatment), size = 2) +
  theme_gdocs() + ggtitle('0.5% DHQ experiment') +
  facet_wrap(~type, scales = "free_y") + ylab('% Total time') +
  scale_fill_manual(values = c("dhq" = "#dc4600", "control" = "#0064c8")) + 
  scale_color_manual(values = c("dhq" = "#dc4600", "control" = "#0064c8"))
```

##### Percent of duration of events: statistics per behavior

Breathing:

```
desc = Summarize(breathing ~ treatment + week, data=percent)
kbl(desc, caption = "Descriptive statistics for Percent of time breathing in 0.5% Experiment") %>% kable_paper("hover", full_width = F)
```

Descriptive statistics for Percent of time breathing in 0.5% Experiment

| treatment | week | n | mean | sd | min | Q1 | median | Q3 | max | percZero |
| --- | --- | --- | --- | --- | --- | --- | --- | --- | --- | --- |
| control | 0 | 20 | 0.2300000 | 0.2556725 | 0.0 | 0.10 | 0.20 | 0.200 | 1.2 | 5.000000 |
| dhq | 0 | 19 | 0.2947368 | 0.2120631 | 0.1 | 0.10 | 0.30 | 0.350 | 0.9 | 0.000000 |
| control | 1 | 19 | 0.3894737 | 0.2884725 | 0.1 | 0.20 | 0.30 | 0.500 | 1.2 | 0.000000 |
| dhq | 1 | 17 | 0.3235294 | 0.3011009 | 0.0 | 0.10 | 0.20 | 0.400 | 1.0 | 5.882353 |
| control | 2 | 16 | 0.5125000 | 0.6396614 | 0.1 | 0.20 | 0.35 | 0.600 | 2.8 | 0.000000 |
| dhq | 2 | 15 | 0.4733333 | 0.4233652 | 0.1 | 0.15 | 0.30 | 0.750 | 1.4 | 0.000000 |
| control | 3 | 14 | 0.3357143 | 0.1945691 | 0.1 | 0.20 | 0.35 | 0.475 | 0.7 | 0.000000 |
| dhq | 3 | 13 | 0.2538462 | 0.1808101 | 0.1 | 0.10 | 0.20 | 0.400 | 0.6 | 0.000000 |

```
ggplot(percent, aes(breathing, fill = treatment)) + 
  geom_density(alpha = 0.5) + facet_wrap(~week) +
  scale_fill_manual(values = c("dhq" = "#dc4600", "control" = "#0064c8")) + 
  scale_color_manual(values = c("dhq" = "#dc4600", "control" = "#0064c8"))
```

```
percent %>% group_by(treatment, week) %>% shapiro_test(breathing)
```

```
## # A tibble: 8 x 5
##   treatment  week variable  statistic          p
##   <chr>     <int> <chr>         <dbl>      <dbl>
## 1 control       0 breathing     0.605 0.00000332
## 2 control       1 breathing     0.824 0.00262   
## 3 control       2 breathing     0.550 0.00000564
## 4 control       3 breathing     0.926 0.270     
## 5 dhq           0 breathing     0.805 0.00136   
## 6 dhq           1 breathing     0.829 0.00517   
## 7 dhq           2 breathing     0.834 0.0104    
## 8 dhq           3 breathing     0.808 0.00844
```

```
percent$trans.breathing = sqrt(percent$breathing)
percent %>% group_by(treatment, week) %>% shapiro_test(trans.breathing)
```

```
## # A tibble: 8 x 5
##   treatment  week variable        statistic        p
##   <chr>     <int> <chr>               <dbl>    <dbl>
## 1 control       0 trans.breathing     0.818 0.00163 
## 2 control       1 trans.breathing     0.898 0.0444  
## 3 control       2 trans.breathing     0.762 0.000899
## 4 control       3 trans.breathing     0.926 0.265   
## 5 dhq           0 trans.breathing     0.874 0.0167  
## 6 dhq           1 trans.breathing     0.925 0.176   
## 7 dhq           2 trans.breathing     0.882 0.0505  
## 8 dhq           3 trans.breathing     0.809 0.00875
```

```
ggplot(percent, aes(trans.breathing, fill = treatment)) + 
  geom_density(alpha = 0.5) + facet_wrap(~week) +
  scale_fill_manual(values = c("dhq" = "#dc4600", "control" = "#0064c8")) + 
  scale_color_manual(values = c("dhq" = "#dc4600", "control" = "#0064c8"))
```

```
lmer1 <- lmer(trans.breathing ~ week * treatment + tank + (1 | individual), percent)
lmer2 <- lmer(trans.breathing ~ week * treatment + (1 | individual), percent)
anova(lmer1, lmer2)
```

```
## Data: percent
## Models:
## lmer2: trans.breathing ~ week * treatment + (1 | individual)
## lmer1: trans.breathing ~ week * treatment + tank + (1 | individual)
##       npar    AIC    BIC  logLik deviance  Chisq Df Pr(>Chisq)  
## lmer2    6 2.4591 19.801  4.7704  -9.5409                       
## lmer1   18 6.2425 58.269 14.8788 -29.7575 20.217 12     0.0631 .
## ---
## Signif. codes:  0 '***' 0.001 '**' 0.01 '*' 0.05 '.' 0.1 ' ' 1
```

```
breathing.percent.lmer = lmer1
summary(breathing.percent.lmer)
```

```
## Linear mixed model fit by REML. t-tests use Satterthwaite's method [
## lmerModLmerTest]
## Formula: trans.breathing ~ week * treatment + tank + (1 | individual)
##    Data: percent
## 
## REML criterion at convergence: 28.1
## 
## Scaled residuals: 
##     Min      1Q  Median      3Q     Max 
## -1.6766 -0.5694 -0.2410  0.6238  4.0115 
## 
## Random effects:
##  Groups     Name        Variance Std.Dev.
##  individual (Intercept) 0.00445  0.06671 
##  Residual               0.04988  0.22334 
## Number of obs: 133, groups:  individual, 47
## 
## Fixed effects:
##                   Estimate Std. Error       df t value Pr(>|t|)   
## (Intercept)        0.35169    0.10422 26.25315   3.375  0.00231 **
## week               0.04556    0.02477 96.52627   1.839  0.06899 . 
## treatmentdhq       0.02972    0.06466 90.72099   0.460  0.64691   
## tankA07            0.07270    0.15051 24.25646   0.483  0.63341   
## tankA09            0.05115    0.19167 72.47934   0.267  0.79034   
## tankA15            0.07629    0.13373 19.42777   0.570  0.57488   
## tankB07            0.27646    0.11232 22.28454   2.461  0.02204 * 
## tankB10            0.01038    0.13373 19.42777   0.078  0.93893   
## tankB15            0.16245    0.10847 23.80935   1.498  0.14737   
## tankB16            0.24134    0.13701 21.24703   1.761  0.09254 . 
## tankC07            0.02254    0.12476 25.86788   0.181  0.85802   
## tankC10            0.02381    0.11999 21.80340   0.198  0.84452   
## tankC11            0.20445    0.14730 28.20433   1.388  0.17600   
## tankC13            0.17072    0.11917 21.20252   1.433  0.16657   
## tankC15            0.15756    0.13737 21.52302   1.147  0.26399   
## week:treatmentdhq -0.03446    0.03578 96.18833  -0.963  0.33788   
## ---
## Signif. codes:  0 '***' 0.001 '**' 0.01 '*' 0.05 '.' 0.1 ' ' 1
```

Smelling:

```
ggplot(percent, aes(smelling, fill = treatment)) + 
  geom_density(alpha = 0.5) + facet_wrap(~week) +
  scale_fill_manual(values = c("dhq" = "#dc4600", "control" = "#0064c8")) + 
  scale_color_manual(values = c("dhq" = "#dc4600", "control" = "#0064c8"))
```

```
percent %>% group_by(treatment, week) %>% shapiro_test(smelling)
```

```
## # A tibble: 8 x 5
##   treatment  week variable statistic          p
##   <chr>     <int> <chr>        <dbl>      <dbl>
## 1 control       0 smelling     0.785 0.000691  
## 2 control       1 smelling     0.957 0.509     
## 3 control       2 smelling     0.835 0.00827   
## 4 control       3 smelling     0.904 0.152     
## 5 dhq           0 smelling     0.547 0.00000135
## 6 dhq           1 smelling     0.818 0.00356   
## 7 dhq           2 smelling     0.927 0.280     
## 8 dhq           3 smelling     0.834 0.0237
```

```
percent$trans.smelling = sqrt(percent$smelling)
percent %>% group_by(treatment, week) %>% shapiro_test(trans.smelling)
```

```
## # A tibble: 8 x 5
##   treatment  week variable       statistic       p
##   <chr>     <int> <chr>              <dbl>   <dbl>
## 1 control       0 trans.smelling     0.944 0.307  
## 2 control       1 trans.smelling     0.935 0.217  
## 3 control       2 trans.smelling     0.941 0.361  
## 4 control       3 trans.smelling     0.898 0.126  
## 5 dhq           0 trans.smelling     0.807 0.00145
## 6 dhq           1 trans.smelling     0.884 0.0368 
## 7 dhq           2 trans.smelling     0.919 0.209  
## 8 dhq           3 trans.smelling     0.909 0.207
```

```
ggplot(percent, aes(trans.breathing, fill = treatment)) + 
  geom_density(alpha = 0.5) + facet_wrap(~week) +
  scale_fill_manual(values = c("dhq" = "#dc4600", "control" = "#0064c8")) + 
  scale_color_manual(values = c("dhq" = "#dc4600", "control" = "#0064c8"))
```

```
lmer1 <- lmer(trans.smelling ~ week * treatment + tank + (1 | individual), percent)
lmer2 <- lmer(trans.smelling ~ week * treatment + (1 | individual), percent)
anova(lmer1, lmer2)
```

```
## Data: percent
## Models:
## lmer2: trans.smelling ~ week * treatment + (1 | individual)
## lmer1: trans.smelling ~ week * treatment + tank + (1 | individual)
##       npar    AIC    BIC  logLik deviance  Chisq Df Pr(>Chisq)
## lmer2    6 377.23 394.39 -182.61   365.23                     
## lmer1   18 388.74 440.22 -176.37   352.74 12.486 12     0.4075
```

```
smelling.percent.lmer = lmer2
summary(smelling.percent.lmer)
```

```
## Linear mixed model fit by REML. t-tests use Satterthwaite's method [
## lmerModLmerTest]
## Formula: trans.smelling ~ week * treatment + (1 | individual)
##    Data: percent
## 
## REML criterion at convergence: 374.4
## 
## Scaled residuals: 
##     Min      1Q  Median      3Q     Max 
## -1.9552 -0.7611  0.0156  0.6625  3.7198 
## 
## Random effects:
##  Groups     Name        Variance Std.Dev.
##  individual (Intercept) 0.1969   0.4437  
##  Residual               0.8631   0.9291  
## Number of obs: 129, groups:  individual, 46
## 
## Fixed effects:
##                    Estimate Std. Error        df t value Pr(>|t|)    
## (Intercept)         1.44722    0.20562 104.17487   7.038 2.13e-10 ***
## week                0.16803    0.10682 100.37762   1.573    0.119    
## treatmentdhq       -0.05194    0.29190 105.68838  -0.178    0.859    
## week:treatmentdhq  -0.05739    0.15287  99.89790  -0.375    0.708    
## ---
## Signif. codes:  0 '***' 0.001 '**' 0.01 '*' 0.05 '.' 0.1 ' ' 1
## 
## Correlation of Fixed Effects:
##             (Intr) week   trtmnt
## week        -0.691              
## treatmntdhq -0.704  0.487       
## wk:trtmntdh  0.483 -0.699 -0.681
```

Near the food:

```
desc = Summarize(near ~ treatment + week, data=percent)
kbl(desc, caption = "Descriptive statistics for Percent of time spent close to the food in 0.5% Experiment") %>% kable_paper("hover", full_width = F)
```

Descriptive statistics for Percent of time spent close to the food in 0.5% Experiment

| treatment | week | n | nvalid | mean | sd | min | Q1 | median | Q3 | max | percZero |
| --- | --- | --- | --- | --- | --- | --- | --- | --- | --- | --- | --- |
| control | 0 | 20 | 19 | 8.278947 | 11.19611 | 0 | 0.900 | 3.20 | 12.600 | 40.2 | 21.052632 |
| dhq | 0 | 19 | 17 | 10.105882 | 13.03963 | 0 | 0.000 | 1.40 | 13.200 | 45.4 | 29.411765 |
| control | 1 | 19 | 16 | 12.368750 | 10.21089 | 0 | 2.450 | 11.35 | 20.425 | 29.6 | 12.500000 |
| dhq | 1 | 17 | 13 | 15.615385 | 16.36571 | 0 | 0.200 | 13.80 | 22.400 | 43.1 | 23.076923 |
| control | 2 | 16 | 14 | 18.492857 | 15.58394 | 0 | 4.700 | 17.55 | 30.625 | 45.0 | 21.428571 |
| dhq | 2 | 15 | 13 | 13.261539 | 13.26188 | 0 | 4.300 | 10.40 | 19.200 | 43.0 | 7.692308 |
| control | 3 | 14 | 13 | 21.169231 | 17.93945 | 0 | 7.100 | 16.10 | 35.400 | 51.2 | 15.384615 |
| dhq | 3 | 13 | 10 | 11.980000 | 17.07850 | 0 | 0.625 | 6.10 | 16.150 | 56.1 | 30.000000 |

```
ggplot(percent, aes(near, fill = treatment)) + 
  geom_density(alpha = 0.5) + facet_wrap(~week) +
  scale_fill_manual(values = c("dhq" = "#dc4600", "control" = "#0064c8")) + 
  scale_color_manual(values = c("dhq" = "#dc4600", "control" = "#0064c8"))
```

```
percent %>% group_by(treatment, week) %>% shapiro_test(near)
```

```
## # A tibble: 8 x 5
##   treatment  week variable statistic        p
##   <chr>     <int> <chr>        <dbl>    <dbl>
## 1 control       0 near         0.741 0.000178
## 2 control       1 near         0.920 0.168   
## 3 control       2 near         0.922 0.233   
## 4 control       3 near         0.885 0.0834  
## 5 dhq           0 near         0.784 0.00124 
## 6 dhq           1 near         0.846 0.0256  
## 7 dhq           2 near         0.868 0.0491  
## 8 dhq           3 near         0.724 0.00173
```

```
percent$trans.near = sqrt(percent$near)
percent %>% group_by(treatment, week) %>% shapiro_test(trans.near)
```

```
## # A tibble: 8 x 5
##   treatment  week variable   statistic      p
##   <chr>     <int> <chr>          <dbl>  <dbl>
## 1 control       0 trans.near     0.901 0.0514
## 2 control       1 trans.near     0.919 0.165 
## 3 control       2 trans.near     0.898 0.107 
## 4 control       3 trans.near     0.922 0.268 
## 5 dhq           0 trans.near     0.861 0.0156
## 6 dhq           1 trans.near     0.894 0.110 
## 7 dhq           2 trans.near     0.972 0.921 
## 8 dhq           3 trans.near     0.916 0.323
```

```
ggplot(percent, aes(trans.breathing, fill = treatment)) + 
  geom_density(alpha = 0.5) + facet_wrap(~week) +
  scale_fill_manual(values = c("dhq" = "#dc4600", "control" = "#0064c8")) + 
  scale_color_manual(values = c("dhq" = "#dc4600", "control" = "#0064c8"))
```

```
lmer1 <- lmer(trans.near ~ week * treatment + tank + (1 | individual), percent)
lmer2 <- lmer(trans.near ~ week * treatment + (1 | individual), percent)
anova(lmer1, lmer2)
```

```
## Data: percent
## Models:
## lmer2: trans.near ~ week * treatment + (1 | individual)
## lmer1: trans.near ~ week * treatment + tank + (1 | individual)
##       npar   AIC    BIC  logLik deviance  Chisq Df Pr(>Chisq)
## lmer2    6 507.9 524.37 -247.95    495.9                     
## lmer1   18 521.6 571.01 -242.80    485.6 10.305 12     0.5892
```

```
near.percent.lmer = lmer2
summary(near.percent.lmer)
```

```
## Linear mixed model fit by REML. t-tests use Satterthwaite's method [
## lmerModLmerTest]
## Formula: trans.near ~ week * treatment + (1 | individual)
##    Data: percent
## 
## REML criterion at convergence: 498.8
## 
## Scaled residuals: 
##      Min       1Q   Median       3Q      Max 
## -1.82322 -0.73200 -0.09154  0.69323  2.16603 
## 
## Random effects:
##  Groups     Name        Variance Std.Dev.
##  individual (Intercept) 0.8386   0.9158  
##  Residual               3.8232   1.9553  
## Number of obs: 115, groups:  individual, 45
## 
## Fixed effects:
##                   Estimate Std. Error      df t value Pr(>|t|)    
## (Intercept)         2.2740     0.4386 94.0565   5.184 1.24e-06 ***
## week                0.6522     0.2261 88.0434   2.885  0.00492 ** 
## treatmentdhq        0.2965     0.6425 94.6467   0.461  0.64551    
## week:treatmentdhq  -0.5084     0.3370 91.4682  -1.509  0.13485    
## ---
## Signif. codes:  0 '***' 0.001 '**' 0.01 '*' 0.05 '.' 0.1 ' ' 1
## 
## Correlation of Fixed Effects:
##             (Intr) week   trtmnt
## week        -0.683              
## treatmntdhq -0.683  0.466       
## wk:trtmntdh  0.458 -0.671 -0.684
```

Testing:

```
desc = Summarize(testing ~ treatment + week, data=percent)
kbl(desc, caption = "Descriptive statistics for Percent of time testing in 0.5% Experiment") %>% kable_paper("hover", full_width = F)
```

Descriptive statistics for Percent of time testing in 0.5% Experiment

| treatment | week | n | nvalid | mean | sd | min | Q1 | median | Q3 | max | percZero |
| --- | --- | --- | --- | --- | --- | --- | --- | --- | --- | --- | --- |
| control | 0 | 20 | 13 | 0.6461538 | 1.1565865 | 0 | 0.000 | 0.00 | 0.800 | 3.5 | 61.53846 |
| dhq | 0 | 19 | 16 | 0.9937500 | 1.4521105 | 0 | 0.000 | 0.05 | 1.600 | 4.0 | 50.00000 |
| control | 1 | 19 | 16 | 0.7187500 | 0.9253603 | 0 | 0.000 | 0.20 | 1.025 | 2.6 | 37.50000 |
| dhq | 1 | 17 | 12 | 1.8833333 | 2.9522975 | 0 | 0.000 | 0.10 | 2.700 | 9.3 | 50.00000 |
| control | 2 | 16 | 14 | 1.0285714 | 1.0957460 | 0 | 0.125 | 0.75 | 1.350 | 3.3 | 21.42857 |
| dhq | 2 | 15 | 12 | 1.5500000 | 4.0241318 | 0 | 0.000 | 0.20 | 0.700 | 14.2 | 41.66667 |
| control | 3 | 14 | 11 | 1.0909091 | 1.0222079 | 0 | 0.200 | 0.70 | 1.950 | 2.7 | 27.27273 |
| dhq | 3 | 13 | 10 | 0.5200000 | 1.0747610 | 0 | 0.000 | 0.20 | 0.275 | 3.5 | 40.00000 |

```
ggplot(percent, aes(testing, fill = treatment)) + 
  geom_density(alpha = 0.5) + facet_wrap(~week) +
  scale_fill_manual(values = c("dhq" = "#dc4600", "control" = "#0064c8")) + 
  scale_color_manual(values = c("dhq" = "#dc4600", "control" = "#0064c8"))
```

```
percent %>% group_by(treatment, week) %>% shapiro_test(testing)
```

```
## # A tibble: 8 x 5
##   treatment  week variable statistic          p
##   <chr>     <int> <chr>        <dbl>      <dbl>
## 1 control       0 testing      0.641 0.000152  
## 2 control       1 testing      0.780 0.00150   
## 3 control       2 testing      0.858 0.0286    
## 4 control       3 testing      0.887 0.127     
## 5 dhq           0 testing      0.715 0.000247  
## 6 dhq           1 testing      0.721 0.00134   
## 7 dhq           2 testing      0.434 0.00000595
## 8 dhq           3 testing      0.536 0.00000958
```

```
percent$trans.testing = sqrt(percent$testing)
percent %>% group_by(treatment, week) %>% shapiro_test(trans.testing)
```

```
## # A tibble: 8 x 5
##   treatment  week variable      statistic        p
##   <chr>     <int> <chr>             <dbl>    <dbl>
## 1 control       0 trans.testing     0.722 0.000918
## 2 control       1 trans.testing     0.853 0.0152  
## 3 control       2 trans.testing     0.937 0.382   
## 4 control       3 trans.testing     0.887 0.128   
## 5 dhq           0 trans.testing     0.774 0.00125 
## 6 dhq           1 trans.testing     0.803 0.0100  
## 7 dhq           2 trans.testing     0.692 0.000704
## 8 dhq           3 trans.testing     0.782 0.00868
```

```
ggplot(percent, aes(trans.breathing, fill = treatment)) + 
  geom_density(alpha = 0.5) + facet_wrap(~week) +
  scale_fill_manual(values = c("dhq" = "#dc4600", "control" = "#0064c8")) + 
  scale_color_manual(values = c("dhq" = "#dc4600", "control" = "#0064c8"))
```

```
lmer1 <- lmer(trans.testing ~ week * treatment + tank + (1 | individual), percent)
lmer2 <- lmer(trans.testing ~ week * treatment + (1 | individual), percent)
anova(lmer1, lmer2)
```

```
## Data: percent
## Models:
## lmer2: trans.testing ~ week * treatment + (1 | individual)
## lmer1: trans.testing ~ week * treatment + tank + (1 | individual)
##       npar    AIC    BIC  logLik deviance  Chisq Df Pr(>Chisq)
## lmer2    6 247.20 263.07 -117.60   235.20                     
## lmer1   18 261.34 308.94 -112.67   225.34 9.8628 12      0.628
```

```
testing.percent.lmer = lmer2
summary(testing.percent.lmer)
```

```
## Linear mixed model fit by REML. t-tests use Satterthwaite's method [
## lmerModLmerTest]
## Formula: trans.testing ~ week * treatment + (1 | individual)
##    Data: percent
## 
## REML criterion at convergence: 245.9
## 
## Scaled residuals: 
##     Min      1Q  Median      3Q     Max 
## -1.1948 -0.6876 -0.2125  0.4847  4.2102 
## 
## Random effects:
##  Groups     Name        Variance Std.Dev.
##  individual (Intercept) 0.08175  0.2859  
##  Residual               0.51264  0.7160  
## Number of obs: 104, groups:  individual, 44
## 
## Fixed effects:
##                   Estimate Std. Error       df t value Pr(>|t|)   
## (Intercept)        0.48469    0.17814 91.06109   2.721   0.0078 **
## week               0.14913    0.09305 78.96308   1.603   0.1130   
## treatmentdhq       0.26690    0.24764 89.49234   1.078   0.2840   
## week:treatmentdhq -0.19995    0.13174 84.09917  -1.518   0.1328   
## ---
## Signif. codes:  0 '***' 0.001 '**' 0.01 '*' 0.05 '.' 0.1 ' ' 1
## 
## Correlation of Fixed Effects:
##             (Intr) week   trtmnt
## week        -0.748              
## treatmntdhq -0.719  0.538       
## wk:trtmntdh  0.528 -0.706 -0.729
```

Eating:

```
desc = Summarize(eating ~ treatment + week, data=percent)
kbl(desc, caption = "Descriptive statistics for Percent of time eating in 0.5% Experiment") %>% kable_paper("hover", full_width = F)
```

Descriptive statistics for Percent of time eating in 0.5% Experiment

| treatment | week | n | nvalid | mean | sd | min | Q1 | median | Q3 | max | percZero |
| --- | --- | --- | --- | --- | --- | --- | --- | --- | --- | --- | --- |
| control | 0 | 20 | 14 | 2.671429 | 4.260269 | 0 | 0.000 | 0.35 | 3.500 | 14.2 | 50.00000 |
| dhq | 0 | 19 | 16 | 2.893750 | 4.179229 | 0 | 0.000 | 1.50 | 3.625 | 15.8 | 50.00000 |
| control | 1 | 19 | 16 | 5.125000 | 5.759109 | 0 | 0.000 | 3.15 | 7.075 | 17.5 | 31.25000 |
| dhq | 1 | 17 | 12 | 6.083333 | 9.542139 | 0 | 0.000 | 0.25 | 9.350 | 30.7 | 50.00000 |
| control | 2 | 16 | 13 | 9.607692 | 7.784757 | 0 | 0.100 | 9.80 | 14.100 | 21.6 | 23.07692 |
| dhq | 2 | 15 | 12 | 5.108333 | 8.237879 | 0 | 0.375 | 2.90 | 4.500 | 29.3 | 16.66667 |
| control | 3 | 14 | 11 | 9.027273 | 9.613542 | 0 | 0.900 | 8.40 | 10.950 | 31.7 | 27.27273 |
| dhq | 3 | 13 | 9 | 7.522222 | 13.459270 | 0 | 0.000 | 0.10 | 10.800 | 40.6 | 33.33333 |

```
ggplot(percent, aes(eating, fill = treatment)) + 
  geom_density(alpha = 0.5) + facet_wrap(~week) +
  scale_fill_manual(values = c("dhq" = "#dc4600", "control" = "#0064c8")) + 
  scale_color_manual(values = c("dhq" = "#dc4600", "control" = "#0064c8"))
```

```
percent %>% group_by(treatment, week) %>% shapiro_test(eating)
```

```
## # A tibble: 8 x 5
##   treatment  week variable statistic        p
##   <chr>     <int> <chr>        <dbl>    <dbl>
## 1 control       0 eating       0.691 0.000298
## 2 control       1 eating       0.827 0.00627 
## 3 control       2 eating       0.892 0.105   
## 4 control       3 eating       0.852 0.0453  
## 5 dhq           0 eating       0.714 0.000245
## 6 dhq           1 eating       0.714 0.00116 
## 7 dhq           2 eating       0.639 0.000231
## 8 dhq           3 eating       0.650 0.000373
```

```
percent$trans.eating = sqrt(percent$eating)
percent %>% group_by(treatment, week) %>% shapiro_test(trans.eating)
```

```
## # A tibble: 8 x 5
##   treatment  week variable     statistic       p
##   <chr>     <int> <chr>            <dbl>   <dbl>
## 1 control       0 trans.eating     0.806 0.00600
## 2 control       1 trans.eating     0.900 0.0817 
## 3 control       2 trans.eating     0.818 0.0111 
## 4 control       3 trans.eating     0.914 0.269  
## 5 dhq           0 trans.eating     0.802 0.00287
## 6 dhq           1 trans.eating     0.797 0.00856
## 7 dhq           2 trans.eating     0.898 0.151  
## 8 dhq           3 trans.eating     0.801 0.0207
```

```
ggplot(percent, aes(trans.breathing, fill = treatment)) + 
  geom_density(alpha = 0.5) + facet_wrap(~week) +
  scale_fill_manual(values = c("dhq" = "#dc4600", "control" = "#0064c8")) + 
  scale_color_manual(values = c("dhq" = "#dc4600", "control" = "#0064c8"))
```

```
lmer1 <- lmer(trans.eating ~ week * treatment + tank + (1 | individual), percent)
lmer2 <- lmer(trans.eating ~ week * treatment + (1 | individual), percent)
anova(lmer1, lmer2)
```

```
## Data: percent
## Models:
## lmer2: trans.eating ~ week * treatment + (1 | individual)
## lmer1: trans.eating ~ week * treatment + tank + (1 | individual)
##       npar    AIC    BIC  logLik deviance  Chisq Df Pr(>Chisq)  
## lmer2    6 392.69 408.50 -190.34   380.69                       
## lmer1   18 394.01 441.44 -179.01   358.01 22.678 12    0.03058 *
## ---
## Signif. codes:  0 '***' 0.001 '**' 0.01 '*' 0.05 '.' 0.1 ' ' 1
```

```
eating.percent.lmer = lmer1
summary(eating.percent.lmer)
```

```
## Linear mixed model fit by REML. t-tests use Satterthwaite's method [
## lmerModLmerTest]
## Formula: trans.eating ~ week * treatment + tank + (1 | individual)
##    Data: percent
## 
## REML criterion at convergence: 347.6
## 
## Scaled residuals: 
##     Min      1Q  Median      3Q     Max 
## -2.0292 -0.5454 -0.0292  0.5866  2.0365 
## 
## Random effects:
##  Groups     Name        Variance Std.Dev.
##  individual (Intercept) 0.8137   0.9021  
##  Residual               1.6893   1.2997  
## Number of obs: 103, groups:  individual, 41
## 
## Fixed effects:
##                   Estimate Std. Error       df t value Pr(>|t|)   
## (Intercept)        3.17924    1.37697 32.37040   2.309  0.02748 * 
## week               0.55740    0.17212 72.69735   3.238  0.00181 **
## treatmentdhq      -0.07598    0.51971 55.07405  -0.146  0.88430   
## tankA07           -1.61819    1.60100 29.34070  -1.011  0.32041   
## tankA09           -1.49844    1.73065 41.33051  -0.866  0.39159   
## tankA15           -3.33824    1.53049 26.37226  -2.181  0.03827 * 
## tankB07            0.15377    1.44899 32.57728   0.106  0.91613   
## tankB10           -1.74247    1.53049 26.37226  -1.139  0.26515   
## tankB15           -2.20177    1.38111 30.98766  -1.594  0.12104   
## tankB16           -2.21751    1.54033 26.98688  -1.440  0.16147   
## tankC07           -0.89498    1.53089 32.78318  -0.585  0.56281   
## tankC10           -1.85936    1.45174 29.84623  -1.281  0.21013   
## tankC11           -2.25620    1.57005 28.99173  -1.437  0.16142   
## tankC13           -2.37739    1.43166 28.72989  -1.661  0.10767   
## tankC15           -2.13569    1.53890 26.95463  -1.388  0.17656   
## week:treatmentdhq -0.43382    0.25198 74.31383  -1.722  0.08930 . 
## ---
## Signif. codes:  0 '***' 0.001 '**' 0.01 '*' 0.05 '.' 0.1 ' ' 1
```

##### Mean duration of events: plots

Subset mean duration of events, and obtain plots.

```
# subset dataframe
mean = subset(feeding05, select = c('id', 'duration_mean', 'behavior'))
mean = subset(mean, behavior != 'breathing' & behavior != 'moving' & behavior != 'turning')
mean = spread(mean, behavior, duration_mean)
mean = merge(info05, mean, by = 'id')
breathing = subset(feeding05, behavior == 'breathing' & modifier != 'NA')
breathing.mean = aggregate(breathing$duration_mean, by = list('id'=breathing$id), FUN = sum)
names(breathing.mean)[names(breathing.mean) == "x"] <- "breathing"
mean = merge(mean, breathing.mean, by = 'id')

# remove outliers
mean$near = ifelse(mean$near > 500, NA, mean$near)
mean$smelling = ifelse(mean$smelling > 60, NA, mean$smelling)

# plots
mean.gather = gather(mean, type, measurement, breathing, smelling,near,testing, eating)
ggplot(mean.gather, aes(week, measurement, colour = treatment, group = individual)) +
  geom_smooth(method=loess, aes(group = treatment, fill=treatment), size = 2) +
  theme_gdocs() + ggtitle('0.5% DHQ experiment') +
  facet_wrap(~type, scales = "free_y") + ylab('Mean duration of events (s)') +
  scale_fill_manual(values = c("dhq" = "#dc4600", "control" = "#0064c8")) + 
  scale_color_manual(values = c("dhq" = "#dc4600", "control" = "#0064c8"))
```

##### Mean duration of events: statistics per behavior

Breathing:

```
desc = Summarize(breathing ~ treatment + week, data=mean)
kbl(desc, caption = "Descriptive statistics for Average time per breath in 0.5% Experiment") %>% kable_paper("hover", full_width = F)
```

Descriptive statistics for Average time per breath in 0.5% Experiment

| treatment | week | n | mean | sd | min | Q1 | median | Q3 | max |
| --- | --- | --- | --- | --- | --- | --- | --- | --- | --- |
| control | 0 | 20 | 1.913850 | 0.804769 | 0.747 | 1.3125 | 1.8725 | 2.30025 | 3.444 |
| dhq | 0 | 19 | 1.983368 | 1.093581 | 0.625 | 1.3895 | 1.6660 | 2.27050 | 4.671 |
| control | 1 | 19 | 2.398842 | 1.270959 | 0.668 | 1.6325 | 2.1280 | 3.18600 | 6.079 |
| dhq | 1 | 17 | 2.268412 | 1.397443 | 0.750 | 1.5000 | 1.7530 | 2.74900 | 6.398 |
| control | 2 | 16 | 2.830812 | 1.585723 | 1.233 | 1.4140 | 2.4235 | 3.59100 | 6.002 |
| dhq | 2 | 15 | 3.789400 | 2.097930 | 1.266 | 2.1245 | 3.0000 | 5.26700 | 9.158 |
| control | 3 | 14 | 2.631571 | 1.216290 | 1.249 | 1.5790 | 2.6455 | 3.26325 | 5.392 |
| dhq | 3 | 13 | 3.499154 | 2.353812 | 0.998 | 1.7500 | 2.7490 | 4.99600 | 8.703 |

```
ggplot(mean, aes(breathing, fill = treatment)) + 
  geom_density(alpha = 0.5) + facet_wrap(~week) +
  scale_fill_manual(values = c("dhq" = "#dc4600", "control" = "#0064c8")) + 
  scale_color_manual(values = c("dhq" = "#dc4600", "control" = "#0064c8"))
```

```
mean %>% group_by(treatment, week) %>% shapiro_test(breathing)
```

```
## # A tibble: 8 x 5
##   treatment  week variable  statistic       p
##   <chr>     <int> <chr>         <dbl>   <dbl>
## 1 control       0 breathing     0.945 0.297  
## 2 control       1 breathing     0.906 0.0633 
## 3 control       2 breathing     0.872 0.0290 
## 4 control       3 breathing     0.917 0.196  
## 5 dhq           0 breathing     0.849 0.00657
## 6 dhq           1 breathing     0.818 0.00358
## 7 dhq           2 breathing     0.890 0.0670 
## 8 dhq           3 breathing     0.871 0.0534
```

```
mean$trans.breathing = log10(mean$breathing)
mean %>% group_by(treatment, week) %>% shapiro_test(trans.breathing)
```

```
## # A tibble: 8 x 5
##   treatment  week variable        statistic     p
##   <chr>     <int> <chr>               <dbl> <dbl>
## 1 control       0 trans.breathing     0.959 0.523
## 2 control       1 trans.breathing     0.977 0.902
## 3 control       2 trans.breathing     0.916 0.145
## 4 control       3 trans.breathing     0.945 0.490
## 5 dhq           0 trans.breathing     0.972 0.814
## 6 dhq           1 trans.breathing     0.968 0.785
## 7 dhq           2 trans.breathing     0.971 0.874
## 8 dhq           3 trans.breathing     0.964 0.810
```

```
ggplot(mean, aes(trans.breathing, fill = treatment)) + 
  geom_density(alpha = 0.5) + facet_wrap(~week) +
  scale_fill_manual(values = c("dhq" = "#dc4600", "control" = "#0064c8")) + 
  scale_color_manual(values = c("dhq" = "#dc4600", "control" = "#0064c8"))
```

```
lmer1 <- lmer(trans.breathing ~ week * treatment + tank + (1 | individual), mean)
lmer2 <- lmer(trans.breathing ~ week * treatment + (1 | individual), mean)
anova(lmer1, lmer2)
```

```
## Data: mean
## Models:
## lmer2: trans.breathing ~ week * treatment + (1 | individual)
## lmer1: trans.breathing ~ week * treatment + tank + (1 | individual)
##       npar     AIC    BIC logLik deviance Chisq Df Pr(>Chisq)
## lmer2    6 -9.6822  7.660 10.841  -21.682                    
## lmer1   18  1.6982 53.724 17.151  -34.302 12.62 12     0.3973
```

```
breathing.mean.lmer = lmer2
summary(breathing.mean.lmer)
```

```
## Linear mixed model fit by REML. t-tests use Satterthwaite's method [
## lmerModLmerTest]
## Formula: trans.breathing ~ week * treatment + (1 | individual)
##    Data: mean
## 
## REML criterion at convergence: -0.3
## 
## Scaled residuals: 
##     Min      1Q  Median      3Q     Max 
## -2.2021 -0.6136 -0.1114  0.7510  2.1255 
## 
## Random effects:
##  Groups     Name        Variance Std.Dev.
##  individual (Intercept) 0.009361 0.09675 
##  Residual               0.043536 0.20865 
## Number of obs: 133, groups:  individual, 47
## 
## Fixed effects:
##                    Estimate Std. Error        df t value Pr(>|t|)    
## (Intercept)         0.26437    0.04494 102.54469   5.883  5.1e-08 ***
## week                0.04478    0.02319  95.29062   1.931   0.0564 .  
## treatmentdhq       -0.02151    0.06435 105.67428  -0.334   0.7389    
## week:treatmentdhq   0.04526    0.03337  96.24293   1.356   0.1782    
## ---
## Signif. codes:  0 '***' 0.001 '**' 0.01 '*' 0.05 '.' 0.1 ' ' 1
## 
## Correlation of Fixed Effects:
##             (Intr) week   trtmnt
## week        -0.687              
## treatmntdhq -0.698  0.480       
## wk:trtmntdh  0.478 -0.695 -0.686
```

Smelling:

```
desc = Summarize(smelling ~ treatment + week, data=mean)
kbl(desc, caption = "Descriptive statistics for Average time per smelling event in 0.5% Experiment") %>% kable_paper("hover", full_width = F)
```

Descriptive statistics for Average time per smelling event in 0.5% Experiment

| treatment | week | n | nvalid | mean | sd | min | Q1 | median | Q3 | max | percZero |
| --- | --- | --- | --- | --- | --- | --- | --- | --- | --- | --- | --- |
| control | 0 | 20 | 20 | 4.465150 | 5.874627 | 0.000 | 1.85875 | 2.4665 | 3.69425 | 23.249 | 10.000000 |
| dhq | 0 | 19 | 19 | 4.704842 | 4.895194 | 0.000 | 1.62250 | 3.0040 | 4.89600 | 16.390 | 5.263158 |
| control | 1 | 19 | 19 | 4.499000 | 3.317262 | 0.000 | 2.13050 | 3.6320 | 6.73450 | 12.723 | 5.263158 |
| dhq | 1 | 17 | 16 | 3.822000 | 1.812646 | 1.499 | 2.29375 | 3.5660 | 5.20200 | 6.750 | 0.000000 |
| control | 2 | 16 | 16 | 5.687688 | 7.769852 | 0.000 | 2.65550 | 3.6030 | 5.23750 | 33.404 | 6.250000 |
| dhq | 2 | 15 | 15 | 6.861333 | 9.087097 | 0.000 | 2.69100 | 5.2710 | 7.00250 | 37.376 | 13.333333 |
| control | 3 | 14 | 14 | 6.001143 | 5.031011 | 0.000 | 1.78750 | 5.7185 | 8.36425 | 15.315 | 14.285714 |
| dhq | 3 | 13 | 13 | 5.783385 | 6.876941 | 0.000 | 0.87700 | 2.3990 | 6.50300 | 20.292 | 15.384615 |

```
ggplot(mean, aes(smelling, fill = treatment)) + 
  geom_density(alpha = 0.5) + facet_wrap(~week) +
  scale_fill_manual(values = c("dhq" = "#dc4600", "control" = "#0064c8")) + 
  scale_color_manual(values = c("dhq" = "#dc4600", "control" = "#0064c8"))
```

```
mean %>% group_by(treatment, week) %>% shapiro_test(smelling)
```

```
## # A tibble: 8 x 5
##   treatment  week variable statistic          p
##   <chr>     <int> <chr>        <dbl>      <dbl>
## 1 control       0 smelling     0.630 0.00000613
## 2 control       1 smelling     0.920 0.114     
## 3 control       2 smelling     0.559 0.00000681
## 4 control       3 smelling     0.928 0.288     
## 5 dhq           0 smelling     0.793 0.000893  
## 6 dhq           1 smelling     0.906 0.0995    
## 7 dhq           2 smelling     0.631 0.0000494 
## 8 dhq           3 smelling     0.789 0.00511
```

```
mean$trans.smelling = sqrt(mean$smelling)
mean %>% group_by(treatment, week) %>% shapiro_test(trans.smelling)
```

```
## # A tibble: 8 x 5
##   treatment  week variable       statistic       p
##   <chr>     <int> <chr>              <dbl>   <dbl>
## 1 control       0 trans.smelling     0.853 0.00600
## 2 control       1 trans.smelling     0.979 0.932  
## 3 control       2 trans.smelling     0.835 0.00823
## 4 control       3 trans.smelling     0.946 0.497  
## 5 dhq           0 trans.smelling     0.944 0.314  
## 6 dhq           1 trans.smelling     0.926 0.210  
## 7 dhq           2 trans.smelling     0.899 0.0906 
## 8 dhq           3 trans.smelling     0.930 0.345
```

```
ggplot(mean, aes(trans.breathing, fill = treatment)) + 
  geom_density(alpha = 0.5) + facet_wrap(~week) +
  scale_fill_manual(values = c("dhq" = "#dc4600", "control" = "#0064c8")) + 
  scale_color_manual(values = c("dhq" = "#dc4600", "control" = "#0064c8"))
```

```
lmer1 <- lmer(trans.smelling ~ week * treatment + tank + (1 | individual), mean)
lmer2 <- lmer(trans.smelling ~ week * treatment + (1 | individual), mean)
anova(lmer1, lmer2)
```

```
## Data: mean
## Models:
## lmer2: trans.smelling ~ week * treatment + (1 | individual)
## lmer1: trans.smelling ~ week * treatment + tank + (1 | individual)
##       npar    AIC    BIC  logLik deviance  Chisq Df Pr(>Chisq)
## lmer2    6 403.77 421.07 -195.88   391.77                     
## lmer1   18 417.31 469.20 -190.66   381.31 10.456 12      0.576
```

```
smelling.mean.lmer = lmer2
summary(smelling.mean.lmer)
```

```
## Linear mixed model fit by REML. t-tests use Satterthwaite's method [
## lmerModLmerTest]
## Formula: trans.smelling ~ week * treatment + (1 | individual)
##    Data: mean
## 
## REML criterion at convergence: 400.3
## 
## Scaled residuals: 
##     Min      1Q  Median      3Q     Max 
## -2.0589 -0.4208 -0.0308  0.3823  3.5209 
## 
## Random effects:
##  Groups     Name        Variance Std.Dev.
##  individual (Intercept) 0.3766   0.6137  
##  Residual               0.8987   0.9480  
## Number of obs: 132, groups:  individual, 47
## 
## Fixed effects:
##                   Estimate Std. Error       df t value Pr(>|t|)    
## (Intercept)        1.81461    0.22376 91.36537   8.110 2.22e-12 ***
## week               0.11800    0.10623 93.55150   1.111    0.270    
## treatmentdhq       0.06522    0.32015 95.13259   0.204    0.839    
## week:treatmentdhq -0.08063    0.15298 94.71271  -0.527    0.599    
## ---
## Signif. codes:  0 '***' 0.001 '**' 0.01 '*' 0.05 '.' 0.1 ' ' 1
## 
## Correlation of Fixed Effects:
##             (Intr) week   trtmnt
## week        -0.629              
## treatmntdhq -0.699  0.439       
## wk:trtmntdh  0.437 -0.694 -0.626
```

Near the food:

```
desc = Summarize(near ~ treatment + week, data=mean)
kbl(desc, caption = "Descriptive statistics for Average time per event close to the food in 0.5% Experiment") %>% kable_paper("hover", full_width = F)
```

Descriptive statistics for Average time per event close to the food in 0.5% Experiment

| treatment | week | n | mean | sd | min | Q1 | median | Q3 | max | percZero |
| --- | --- | --- | --- | --- | --- | --- | --- | --- | --- | --- |
| control | 0 | 20 | 25.63760 | 29.09933 | 0 | 1.17775 | 15.9890 | 40.7765 | 84.125 | 20.00000 |
| dhq | 0 | 19 | 18.17500 | 23.38172 | 0 | 0.00000 | 5.7640 | 37.8370 | 74.292 | 36.84211 |
| control | 1 | 19 | 35.40105 | 37.46696 | 0 | 1.37450 | 26.0070 | 62.5240 | 119.479 | 26.31579 |
| dhq | 1 | 17 | 49.12235 | 66.97309 | 0 | 0.00000 | 3.5010 | 100.5850 | 179.969 | 41.17647 |
| control | 2 | 16 | 38.99744 | 63.00797 | 0 | 0.00000 | 18.2905 | 52.0720 | 249.471 | 31.25000 |
| dhq | 2 | 15 | 49.98680 | 58.25343 | 0 | 8.13050 | 24.3440 | 81.3125 | 193.568 | 20.00000 |
| control | 3 | 14 | 51.55036 | 77.78169 | 0 | 16.08075 | 29.3105 | 56.7740 | 305.753 | 21.42857 |
| dhq | 3 | 13 | 46.83785 | 91.41621 | 0 | 0.00000 | 19.0470 | 43.1520 | 336.487 | 46.15385 |

```
ggplot(mean, aes(near, fill = treatment)) + 
  geom_density(alpha = 0.5) + facet_wrap(~week) +
  scale_fill_manual(values = c("dhq" = "#dc4600", "control" = "#0064c8")) + 
  scale_color_manual(values = c("dhq" = "#dc4600", "control" = "#0064c8"))
```

```
mean %>% group_by(treatment, week) %>% shapiro_test(near)
```

```
## # A tibble: 8 x 5
##   treatment  week variable statistic         p
##   <chr>     <int> <chr>        <dbl>     <dbl>
## 1 control       0 near         0.819 0.00171  
## 2 control       1 near         0.864 0.0115   
## 3 control       2 near         0.647 0.0000471
## 4 control       3 near         0.603 0.0000435
## 5 dhq           0 near         0.782 0.000628 
## 6 dhq           1 near         0.739 0.000332 
## 7 dhq           2 near         0.829 0.00896  
## 8 dhq           3 near         0.557 0.0000285
```

```
mean$trans.near = sqrt(mean$near)
mean %>% group_by(treatment, week) %>% shapiro_test(trans.near)
```

```
## # A tibble: 8 x 5
##   treatment  week variable   statistic       p
##   <chr>     <int> <chr>          <dbl>   <dbl>
## 1 control       0 trans.near     0.919 0.0941 
## 2 control       1 trans.near     0.916 0.0946 
## 3 control       2 trans.near     0.886 0.0490 
## 4 control       3 trans.near     0.878 0.0540 
## 5 dhq           0 trans.near     0.861 0.0102 
## 6 dhq           1 trans.near     0.798 0.00191
## 7 dhq           2 trans.near     0.942 0.404  
## 8 dhq           3 trans.near     0.801 0.00698
```

```
ggplot(mean, aes(trans.breathing, fill = treatment)) + 
  geom_density(alpha = 0.5) + facet_wrap(~week) +
  scale_fill_manual(values = c("dhq" = "#dc4600", "control" = "#0064c8")) + 
  scale_color_manual(values = c("dhq" = "#dc4600", "control" = "#0064c8"))
```

```
lmer1 <- lmer(trans.near ~ week * treatment + tank + (1 | individual), mean)
lmer2 <- lmer(trans.near ~ week * treatment + (1 | individual), mean)
anova(lmer1, lmer2)
```

```
## Data: mean
## Models:
## lmer2: trans.near ~ week * treatment + (1 | individual)
## lmer1: trans.near ~ week * treatment + tank + (1 | individual)
##       npar    AIC    BIC  logLik deviance  Chisq Df Pr(>Chisq)
## lmer2    6 767.49 784.83 -377.74   755.49                     
## lmer1   18 782.89 834.92 -373.45   746.89 8.5974 12     0.7369
```

```
near.mean.lmer = lmer2
summary(near.mean.lmer)
```

```
## Linear mixed model fit by REML. t-tests use Satterthwaite's method [
## lmerModLmerTest]
## Formula: trans.near ~ week * treatment + (1 | individual)
##    Data: mean
## 
## REML criterion at convergence: 753.7
## 
## Scaled residuals: 
##     Min      1Q  Median      3Q     Max 
## -1.3135 -0.8365 -0.1266  0.6342  2.8404 
## 
## Random effects:
##  Groups     Name        Variance Std.Dev.
##  individual (Intercept)  1.186   1.089   
##  Residual               16.589   4.073   
## Number of obs: 133, groups:  individual, 47
## 
## Fixed effects:
##                    Estimate Std. Error        df t value Pr(>|t|)    
## (Intercept)         4.00079    0.81109 117.31940   4.933  2.7e-06 ***
## week                0.51071    0.44843 104.04513   1.139    0.257    
## treatmentdhq       -0.39848    1.16443 119.30739  -0.342    0.733    
## week:treatmentdhq   0.04079    0.64479 104.58515   0.063    0.950    
## ---
## Signif. codes:  0 '***' 0.001 '**' 0.01 '*' 0.05 '.' 0.1 ' ' 1
## 
## Correlation of Fixed Effects:
##             (Intr) week   trtmnt
## week        -0.741              
## treatmntdhq -0.697  0.516       
## wk:trtmntdh  0.516 -0.695 -0.740
```

Testing:

```
desc = Summarize(testing ~ treatment + week, data=mean)
kbl(desc, caption = "Descriptive statistics for Average time per testing event in 0.5% Experiment") %>% kable_paper("hover", full_width = F)
```

Descriptive statistics for Average time per testing event in 0.5% Experiment

| treatment | week | n | mean | sd | min | Q1 | median | Q3 | max | percZero |
| --- | --- | --- | --- | --- | --- | --- | --- | --- | --- | --- |
| control | 0 | 20 | 2.228650 | 4.916750 | 0 | 0 | 0.0000 | 1.18775 | 19.879 | 75.00000 |
| dhq | 0 | 19 | 6.956263 | 15.673598 | 0 | 0 | 0.0000 | 5.86250 | 61.781 | 57.89474 |
| control | 1 | 19 | 4.005474 | 5.797689 | 0 | 0 | 0.6160 | 6.79800 | 18.334 | 47.36842 |
| dhq | 1 | 17 | 4.767471 | 9.005681 | 0 | 0 | 0.0000 | 8.91500 | 33.549 | 64.70588 |
| control | 2 | 16 | 3.777438 | 3.422177 | 0 | 0 | 3.9415 | 5.67300 | 11.000 | 31.25000 |
| dhq | 2 | 15 | 8.727867 | 21.759912 | 0 | 0 | 0.0000 | 5.30450 | 84.954 | 53.33333 |
| control | 3 | 14 | 3.317857 | 3.423179 | 0 | 0 | 3.4220 | 6.08425 | 9.748 | 42.85714 |
| dhq | 3 | 13 | 3.318846 | 5.352103 | 0 | 0 | 0.0000 | 3.51600 | 15.526 | 53.84615 |

```
ggplot(mean, aes(testing, fill = treatment)) + 
  geom_density(alpha = 0.5) + facet_wrap(~week) +
  scale_fill_manual(values = c("dhq" = "#dc4600", "control" = "#0064c8")) + 
  scale_color_manual(values = c("dhq" = "#dc4600", "control" = "#0064c8"))
```

```
mean %>% group_by(treatment, week) %>% shapiro_test(testing)
```

```
## # A tibble: 8 x 5
##   treatment  week variable statistic           p
##   <chr>     <int> <chr>        <dbl>       <dbl>
## 1 control       0 testing      0.534 0.000000658
## 2 control       1 testing      0.735 0.000151   
## 3 control       2 testing      0.913 0.129      
## 4 control       3 testing      0.856 0.0265     
## 5 dhq           0 testing      0.507 0.000000578
## 6 dhq           1 testing      0.613 0.0000140  
## 7 dhq           2 testing      0.455 0.00000156 
## 8 dhq           3 testing      0.666 0.000257
```

```
mean$trans.testing = sqrt(mean$testing)
mean %>% group_by(treatment, week) %>% shapiro_test(trans.testing)
```

```
## # A tibble: 8 x 5
##   treatment  week variable      statistic          p
##   <chr>     <int> <chr>             <dbl>      <dbl>
## 1 control       0 trans.testing     0.598 0.00000284
## 2 control       1 trans.testing     0.811 0.00164   
## 3 control       2 trans.testing     0.861 0.0198    
## 4 control       3 trans.testing     0.800 0.00506   
## 5 dhq           0 trans.testing     0.707 0.0000670 
## 6 dhq           1 trans.testing     0.695 0.000103  
## 7 dhq           2 trans.testing     0.705 0.000280  
## 8 dhq           3 trans.testing     0.772 0.00320
```

```
ggplot(mean, aes(trans.breathing, fill = treatment)) + 
  geom_density(alpha = 0.5) + facet_wrap(~week) +
  scale_fill_manual(values = c("dhq" = "#dc4600", "control" = "#0064c8")) + 
  scale_color_manual(values = c("dhq" = "#dc4600", "control" = "#0064c8"))
```

```
lmer1 <- lmer(trans.testing ~ week * treatment + tank + (1 | individual), mean)
lmer2 <- lmer(trans.testing ~ week * treatment + (1 | individual), mean)
anova(lmer1, lmer2)
```

```
## Data: mean
## Models:
## lmer2: trans.testing ~ week * treatment + (1 | individual)
## lmer1: trans.testing ~ week * treatment + tank + (1 | individual)
##       npar    AIC    BIC  logLik deviance  Chisq Df Pr(>Chisq)
## lmer2    6 531.69 549.03 -259.84   519.69                     
## lmer1   18 547.34 599.37 -255.67   511.34 8.3479 12     0.7574
```

```
testing.mean.lmer = lmer2
summary(testing.mean.lmer)
```

```
## Linear mixed model fit by REML. t-tests use Satterthwaite's method [
## lmerModLmerTest]
## Formula: trans.testing ~ week * treatment + (1 | individual)
##    Data: mean
## 
## REML criterion at convergence: 525.2
## 
## Scaled residuals: 
##     Min      1Q  Median      3Q     Max 
## -0.9221 -0.7749 -0.5127  0.6067  4.5435 
## 
## Random effects:
##  Groups     Name        Variance Std.Dev.
##  individual (Intercept) 0.02712  0.1647  
##  Residual               2.97792  1.7257  
## Number of obs: 133, groups:  individual, 47
## 
## Fixed effects:
##                   Estimate Std. Error       df t value Pr(>|t|)   
## (Intercept)         0.8959     0.3304 120.8075   2.712  0.00767 **
## week                0.2354     0.1889 102.8819   1.246  0.21547   
## treatmentdhq        0.5418     0.4749 122.5443   1.141  0.25619   
## week:treatmentdhq  -0.2824     0.2715 103.2265  -1.040  0.30066   
## ---
## Signif. codes:  0 '***' 0.001 '**' 0.01 '*' 0.05 '.' 0.1 ' ' 1
## 
## Correlation of Fixed Effects:
##             (Intr) week   trtmnt
## week        -0.770              
## treatmntdhq -0.696  0.536       
## wk:trtmntdh  0.536 -0.696 -0.769
```

Eating:

```
desc = Summarize(eating ~ treatment + week, data=mean)
kbl(desc, caption = "Descriptive statistics for Average time per eating event in 0.5% Experiment") %>% kable_paper("hover", full_width = F)
```

Descriptive statistics for Average time per eating event in 0.5% Experiment

| treatment | week | n | mean | sd | min | Q1 | median | Q3 | max | percZero |
| --- | --- | --- | --- | --- | --- | --- | --- | --- | --- | --- |
| control | 0 | 20 | 9.17695 | 13.68311 | 0 | 0 | 0.0000 | 20.74150 | 36.398 | 65.00000 |
| dhq | 0 | 19 | 11.08626 | 17.78254 | 0 | 0 | 0.0000 | 16.71900 | 60.734 | 52.63158 |
| control | 1 | 19 | 38.11574 | 78.44880 | 0 | 0 | 5.6890 | 34.03700 | 311.053 | 42.10526 |
| dhq | 1 | 17 | 15.40765 | 27.54082 | 0 | 0 | 0.0000 | 28.20000 | 92.017 | 64.70588 |
| control | 2 | 16 | 19.89044 | 25.42398 | 0 | 0 | 10.3745 | 28.01425 | 84.717 | 37.50000 |
| dhq | 2 | 15 | 36.66347 | 66.77989 | 0 | 0 | 17.0260 | 42.50250 | 263.984 | 33.33333 |
| control | 3 | 14 | 19.82279 | 24.99174 | 0 | 0 | 9.6970 | 32.49525 | 83.726 | 42.85714 |
| dhq | 3 | 13 | 50.51292 | 108.86308 | 0 | 0 | 0.0000 | 42.49800 | 364.972 | 53.84615 |

```
ggplot(mean, aes(eating, fill = treatment)) + 
  geom_density(alpha = 0.5) + facet_wrap(~week) +
  scale_fill_manual(values = c("dhq" = "#dc4600", "control" = "#0064c8")) + 
  scale_color_manual(values = c("dhq" = "#dc4600", "control" = "#0064c8"))
```

```
mean %>% group_by(treatment, week) %>% shapiro_test(eating)
```

```
## # A tibble: 8 x 5
##   treatment  week variable statistic          p
##   <chr>     <int> <chr>        <dbl>      <dbl>
## 1 control       0 eating       0.686 0.0000267 
## 2 control       1 eating       0.535 0.00000103
## 3 control       2 eating       0.802 0.00289   
## 4 control       3 eating       0.807 0.00610   
## 5 dhq           0 eating       0.695 0.0000479 
## 6 dhq           1 eating       0.644 0.0000289 
## 7 dhq           2 eating       0.571 0.0000140 
## 8 dhq           3 eating       0.551 0.0000255
```

```
mean$trans.eating = sqrt(mean$eating)
mean %>% group_by(treatment, week) %>% shapiro_test(trans.eating)
```

```
## # A tibble: 8 x 5
##   treatment  week variable     statistic         p
##   <chr>     <int> <chr>            <dbl>     <dbl>
## 1 control       0 trans.eating     0.668 0.0000165
## 2 control       1 trans.eating     0.784 0.000666 
## 3 control       2 trans.eating     0.870 0.0267   
## 4 control       3 trans.eating     0.849 0.0216   
## 5 dhq           0 trans.eating     0.767 0.000392 
## 6 dhq           1 trans.eating     0.686 0.0000806
## 7 dhq           2 trans.eating     0.851 0.0178   
## 8 dhq           3 trans.eating     0.692 0.000463
```

```
ggplot(mean, aes(trans.breathing, fill = treatment)) + 
  geom_density(alpha = 0.5) + facet_wrap(~week) +
  scale_fill_manual(values = c("dhq" = "#dc4600", "control" = "#0064c8")) + 
  scale_color_manual(values = c("dhq" = "#dc4600", "control" = "#0064c8"))
```

```
lmer1 <- lmer(trans.eating ~ week * treatment + tank + (1 | individual), mean)
lmer2 <- lmer(trans.eating ~ week * treatment + (1 | individual), mean)
anova(lmer1, lmer2)
```

```
## Data: mean
## Models:
## lmer2: trans.eating ~ week * treatment + (1 | individual)
## lmer1: trans.eating ~ week * treatment + tank + (1 | individual)
##       npar    AIC    BIC  logLik deviance  Chisq Df Pr(>Chisq)
## lmer2    6 746.57 763.91 -367.28   734.57                     
## lmer1   18 760.75 812.78 -362.38   724.75 9.8151 12     0.6322
```

```
eating.mean.lmer = lmer2
summary(eating.mean.lmer)
```

```
## Linear mixed model fit by REML. t-tests use Satterthwaite's method [
## lmerModLmerTest]
## Formula: trans.eating ~ week * treatment + (1 | individual)
##    Data: mean
## 
## REML criterion at convergence: 733.5
## 
## Scaled residuals: 
##     Min      1Q  Median      3Q     Max 
## -1.1193 -0.6965 -0.4306  0.6598  3.8251 
## 
## Random effects:
##  Groups     Name        Variance Std.Dev.
##  individual (Intercept)  0.5271  0.726   
##  Residual               14.6100  3.822   
## Number of obs: 133, groups:  individual, 47
## 
## Fixed effects:
##                   Estimate Std. Error       df t value Pr(>|t|)   
## (Intercept)         2.4311     0.7447 119.8717   3.265  0.00143 **
## week                0.4138     0.4195 104.8962   0.986  0.32622   
## treatmentdhq       -0.4996     1.0699 121.6185  -0.467  0.64139   
## week:treatmentdhq   0.3314     0.6031 105.3198   0.549  0.58388   
## ---
## Signif. codes:  0 '***' 0.001 '**' 0.01 '*' 0.05 '.' 0.1 ' ' 1
## 
## Correlation of Fixed Effects:
##             (Intr) week   trtmnt
## week        -0.757              
## treatmntdhq -0.696  0.527       
## wk:trtmntdh  0.527 -0.696 -0.756
```

#### Latency

The latency05 table, for the 0.5% DHQ experiment, looks like this:

```
latency05 = read.table("D:/STANFORD/DHQ_FEEDING/code/SM_tables/latency05.csv", header=TRUE, sep=',')
kbl(head(latency05), caption = "Head of latency05 table") %>% kable_paper("hover", full_width = F)
```

Head of latency05 table

| id | individual | treatment | week | behavior | latency | censored |
| --- | --- | --- | --- | --- | --- | --- |
| CT04W0 | CT04 | control | 0 | eating | 1800.0 | 1 |
| CT04W0 | CT04 | control | 0 | near | 1800.0 | 1 |
| CT04W0 | CT04 | control | 0 | smelling | 314.1 | 2 |
| CT04W0 | CT04 | control | 0 | testing | 1800.0 | 1 |
| CT04W2 | CT04 | control | 2 | eating | 811.3 | 2 |
| CT04W2 | CT04 | control | 2 | near | 697.8 | 2 |

We subset this dataframe by behavior for further analysis.

##### Latency to smell

```
smell.lat = subset(latency05, behavior == 'smelling')

smell.lat.fit <- survfit(Surv(latency, censored) ~ treatment + week, data = smell.lat)
ggsurvplot(smell.lat.fit, data=smell.lat, xlab = "seconds", ylab = 'Cummulative events', title = '0.5% DHQ experiment', fun="event", ggtheme = theme_gdocs())
```

```
for (w in 0:3){
  print(cat("WEEK: ",w))
  week.subset = subset(smell.lat, week == w)
  a = survdiff(Surv(latency, censored) ~ treatment, data = week.subset)
  print(a)}
```

```
## WEEK:  0NULL
## Call:
## survdiff(formula = Surv(latency, censored) ~ treatment, data = week.subset)
## 
##                    N Observed Expected (O-E)^2/E (O-E)^2/V
## treatment=control 24       21     20.6   0.00940    0.0189
## treatment=dhq     24       20     20.4   0.00945    0.0189
## 
##  Chisq= 0  on 1 degrees of freedom, p= 0.9 
## WEEK:  1NULL
## Call:
## survdiff(formula = Surv(latency, censored) ~ treatment, data = week.subset)
## 
##                    N Observed Expected (O-E)^2/E (O-E)^2/V
## treatment=control 22       20     16.1     0.947       1.7
## treatment=dhq     22       18     21.9     0.696       1.7
## 
##  Chisq= 1.7  on 1 degrees of freedom, p= 0.2 
## WEEK:  2NULL
## Call:
## survdiff(formula = Surv(latency, censored) ~ treatment, data = week.subset)
## 
##                    N Observed Expected (O-E)^2/E (O-E)^2/V
## treatment=control 24       22     15.5      2.75      4.71
## treatment=dhq     22       16     22.5      1.89      4.71
## 
##  Chisq= 4.7  on 1 degrees of freedom, p= 0.03 
## WEEK:  3NULL
## Call:
## survdiff(formula = Surv(latency, censored) ~ treatment, data = week.subset)
## 
##                    N Observed Expected (O-E)^2/E (O-E)^2/V
## treatment=control 21       15     15.8    0.0399    0.0816
## treatment=dhq     20       16     15.2    0.0415    0.0816
## 
##  Chisq= 0.1  on 1 degrees of freedom, p= 0.8
```

##### Latency to approach the food

```
near.lat = subset(latency05, behavior == 'near')

near.lat.fit <- survfit(Surv(latency, censored) ~ treatment + week, data = near.lat)
ggsurvplot(near.lat.fit, data=smell.lat, xlab = "seconds", ylab = 'Cummulative events', title = '0.5% DHQ experiment', fun="event", ggtheme = theme_gdocs())
```

```
for (w in 0:3){
  print(cat("WEEK: ",w))
  week.subset = subset(near.lat, week == w)
  a = survdiff(Surv(latency, censored) ~ treatment, data = week.subset)
  print(a)}
```

```
## WEEK:  0NULL
## Call:
## survdiff(formula = Surv(latency, censored) ~ treatment, data = week.subset)
## 
##                    N Observed Expected (O-E)^2/E (O-E)^2/V
## treatment=control 24       18     17.1    0.0471    0.0982
## treatment=dhq     24       15     15.9    0.0507    0.0982
## 
##  Chisq= 0.1  on 1 degrees of freedom, p= 0.8 
## WEEK:  1NULL
## Call:
## survdiff(formula = Surv(latency, censored) ~ treatment, data = week.subset)
## 
##                    N Observed Expected (O-E)^2/E (O-E)^2/V
## treatment=control 22       15     11.2     1.265      2.24
## treatment=dhq     22       11     14.8     0.962      2.24
## 
##  Chisq= 2.2  on 1 degrees of freedom, p= 0.1 
## WEEK:  2NULL
## Call:
## survdiff(formula = Surv(latency, censored) ~ treatment, data = week.subset)
## 
##                    N Observed Expected (O-E)^2/E (O-E)^2/V
## treatment=control 24       16     14.2     0.234     0.447
## treatment=dhq     22       14     15.8     0.210     0.447
## 
##  Chisq= 0.4  on 1 degrees of freedom, p= 0.5 
## WEEK:  3NULL
## Call:
## survdiff(formula = Surv(latency, censored) ~ treatment, data = week.subset)
## 
##                    N Observed Expected (O-E)^2/E (O-E)^2/V
## treatment=control 21       13     11.7     0.148      0.29
## treatment=dhq     20       11     12.3     0.141      0.29
## 
##  Chisq= 0.3  on 1 degrees of freedom, p= 0.6
```

##### Latency to test the food

```
test.lat = subset(latency05, behavior == 'testing')

test.lat.fit <- survfit(Surv(latency, censored) ~ treatment + week, data = test.lat)
ggsurvplot(test.lat.fit, data=test.lat, xlab = "seconds", ylab = 'Cummulative events', title = '0.5% DHQ experiment', fun="event", ggtheme = theme_gdocs())
```

```
for (w in 0:3){
  print(cat("WEEK: ",w))
  week.subset = subset(test.lat, week == w)
  a = survdiff(Surv(latency, censored) ~ treatment, data = week.subset)
  print(a)}
```

```
## WEEK:  0NULL
## Call:
## survdiff(formula = Surv(latency, censored) ~ treatment, data = week.subset)
## 
##                    N Observed Expected (O-E)^2/E (O-E)^2/V
## treatment=control 24        6     8.39     0.683      1.44
## treatment=dhq     24       10     7.61     0.754      1.44
## 
##  Chisq= 1.4  on 1 degrees of freedom, p= 0.2 
## WEEK:  1NULL
## Call:
## survdiff(formula = Surv(latency, censored) ~ treatment, data = week.subset)
## 
##                    N Observed Expected (O-E)^2/E (O-E)^2/V
## treatment=control 22       13     8.69      2.13      3.96
## treatment=dhq     22        6    10.31      1.80      3.96
## 
##  Chisq= 4  on 1 degrees of freedom, p= 0.05 
## WEEK:  2NULL
## Call:
## survdiff(formula = Surv(latency, censored) ~ treatment, data = week.subset)
## 
##                    N Observed Expected (O-E)^2/E (O-E)^2/V
## treatment=control 24       15     10.7      1.75      3.29
## treatment=dhq     22        8     12.3      1.52      3.29
## 
##  Chisq= 3.3  on 1 degrees of freedom, p= 0.07 
## WEEK:  3NULL
## Call:
## survdiff(formula = Surv(latency, censored) ~ treatment, data = week.subset)
## 
##                    N Observed Expected (O-E)^2/E (O-E)^2/V
## treatment=control 21       10     8.91     0.134     0.253
## treatment=dhq     20        9    10.09     0.119     0.253
## 
##  Chisq= 0.3  on 1 degrees of freedom, p= 0.6
```

##### Latency to eat

```
eat.lat = subset(latency05, behavior == 'eating')

eat.lat.fit <- survfit(Surv(latency, censored) ~ treatment + week, data = eat.lat)
ggsurvplot(eat.lat.fit, data=eat.lat, xlab = "seconds", ylab = 'Cummulative events', title = '0.5% DHQ experiment', fun="event", ggtheme = theme_gdocs())
```

```
for (w in 0:3){
  print(cat("WEEK: ",w))
  week.subset = subset(eat.lat, week == w)
  a = survdiff(Surv(latency, censored) ~ treatment, data = week.subset)
  print(a)}
```

```
## WEEK:  0NULL
## Call:
## survdiff(formula = Surv(latency, censored) ~ treatment, data = week.subset)
## 
##                    N Observed Expected (O-E)^2/E (O-E)^2/V
## treatment=control 24        8    10.33     0.524      1.15
## treatment=dhq     24       11     8.67     0.623      1.15
## 
##  Chisq= 1.1  on 1 degrees of freedom, p= 0.3 
## WEEK:  1NULL
## Call:
## survdiff(formula = Surv(latency, censored) ~ treatment, data = week.subset)
## 
##                    N Observed Expected (O-E)^2/E (O-E)^2/V
## treatment=control 22       13     8.59      2.26      4.15
## treatment=dhq     22        6    10.41      1.87      4.15
## 
##  Chisq= 4.1  on 1 degrees of freedom, p= 0.04 
## WEEK:  2NULL
## Call:
## survdiff(formula = Surv(latency, censored) ~ treatment, data = week.subset)
## 
##                    N Observed Expected (O-E)^2/E (O-E)^2/V
## treatment=control 24       15     12.3     0.570      1.09
## treatment=dhq     22       11     13.7     0.515      1.09
## 
##  Chisq= 1.1  on 1 degrees of freedom, p= 0.3 
## WEEK:  3NULL
## Call:
## survdiff(formula = Surv(latency, censored) ~ treatment, data = week.subset)
## 
##                    N Observed Expected (O-E)^2/E (O-E)^2/V
## treatment=control 21       10      9.1    0.0887     0.171
## treatment=dhq     20        9      9.9    0.0815     0.171
## 
##  Chisq= 0.2  on 1 degrees of freedom, p= 0.7
```
